# Empirical Models for Anatomical and Physiological Changes in a Human Mother and Fetus During Pregnancy and Gestation

**DOI:** 10.1101/438119

**Authors:** Dustin F. Kapraun, John F. Wambaugh, R. Woodrow Setzer, Richard S. Judson

## Abstract

Many parameters treated as constants in traditional physiologically based pharmacokinetic models must be formulated as time-varying quantities when modeling pregnancy and gestation due to the dramatic physiological and anatomical changes that occur during this period. While several collections of empirical models for such parameters have been published, each has shortcomings. We sought to create a repository of empirical models for tissue volumes, blood flow rates, and other quantities that undergo substantial changes in a human mother and her fetus during the time between conception and birth, and to address deficiencies with similar, previously published repositories. We used maximum likelihood estimation to calibrate various models for the time-varying quantities of interest, and then used the Akaike information criterion to select an optimal model for each quantity. For quantities of interest for which time-course data were not available, we constructed composite models using percentages and/or models describing related quantities. In this way, we developed a comprehensive collection of formulae describing parameters essential for constructing a PBPK model of a human mother and her fetus throughout the approximately 40 weeks of pregnancy and gestation. We included models describing blood flow rates through various fetal blood routes that have no counterparts in adults. Our repository of mathematical models for anatomical and physiological quantities of interest provides a basis for PBPK models of human pregnancy and gestation, and as such, it can ultimately be used to support decision-making with respect to optimal pharmacological dosing and risk assessment for pregnant women and their developing fetuses. *The views expressed in this article are those of the authors and do not necessarily represent the views or policies of the U.S. Environmental Protection Agency*.

**AUTHOR SUMMARY:** Physiologically based pharmacokinetic modeling is a well-known technique for making predictions about internal time-course concentrations of a substance that has entered an organism. This tool is widely used in both pharmaceutical research and human health risk assessment because it harnesses one of the fundamental tenets of both pharmacology and toxicology: it is the concentrations of an active chemical that reach internal target tissues, rather than externally applied “doses”, that govern the extent of the response (whether beneficial or adverse). Constructing physiologically based pharmacokinetic models for pregnancy and gestation presents a considerable challenge because many of the required parameters (such as blood flow rates or tissue volumes) that are typically assumed to be constant in adult models or short-duration simulations cannot be assumed to be constant when modeling pregnancy. Here we present models, stated as functions of gestational age, for anatomical and physiological changes that occur in a human mother and fetus during pregnancy and gestation. We evaluated and selected models by applying a consistent statistical technique, and where possible, we compared results produced by our models to those produced by previously-published models. The collection of pregnancy parameter models presented here represents the most comprehensive such collection to date.

## INTRODUCTION

Human health chemical risk assessments frequently consider pregnant women as a subpopulation of interest based on the relatively high exposure rates and/or susceptibility of this group to various compounds [1,2]. Unfortunately, pregnant women are generally underrepresented in pharmaceutical clinical studies [3] and non-therapeutic chemicals are rarely studied in humans at any life-stage [4–6]. In order to assess the risk posed by a chemical, pharmacokinetic (PK) modeling can be used to relate chemical exposure to potential toxicity in tissues [7]. PK models describe chemical absorption, distribution, metabolism, and elimination by the body [8,9]. Furthermore, such models allow one to quantify the tissue concentrations resulting from external doses, whether they are controlled (e.g., doses administered in a clinical trial or animal toxicity study [10]) or uncontrolled (e.g., through complex environmental exposures [11]).

During gestation there are windows of toxic susceptibility during which chemical insults may induce life-long adverse effects [2,9,12–14]. Mathematical PK models provide a means for predicting fetal tissue exposures to chemicals [12,15]. Given that PK data in pregnant women and in utero infants are unavailable for most chemicals, models are needed to estimate doses of concern based on data collected for non-pregnant adults or from animals [12]. Physiologically based PK (PBPK) models offer an attractive option for extrapolating information in applications such as human health risk assessments [16–18]. Furthermore, PBPK models can be used to understand and potentially replace some of the default uncertainty factors that are typically applied when using toxicity data to establish a reference dose [12,19].

Taking into account an external dose (such as an amount that is consumed orally, applied dermally, or inhaled from ambient air), a PBPK model utilizes information about absorption into the body, distribution and storage throughout the body’s tissues, metabolism within particular tissues, and excretion from the body to estimate internal doses (amounts or concentrations) at various sites within the body [20]. In other words, a PBPK model predicts concentrations in various tissues (e.g., adipose and brain tissues) represented by “compartments” of known volume that are connected by “flows”, most typically of blood [21,22]. Barton et al. [23] identified many ways in which PBPK models may be used for extrapolation, including across life-stages [15,24]. For the current manuscript, we constructed mathematical models describing anatomical and physiological changes associated with human pregnancy and gestation, as we contend that such models must be an integral part of any PBPK model that describes the kinetics of a chemical in a mother and her fetus over a significant portion of the pregnancy.

Several different collections of time-varying formulae describing quantities related to human pregnancy and gestation have already been published, and most recently Dallmann et al. [3] performed a meta-analysis of the literature in order to construct such a set of time-dependent formulae. Prior to this, Luecke and coauthors [9,15,25–27] developed a collection of formulae describing anatomical and physiological changes related to pregnancy and gestation. This latter collection of models has been particularly influential, with more than one hundred citations in peer-reviewed publications related to PBPK models. The following list describes some of the publications and research groups that have provided one or more formulae related to human gestation and pregnancy:

- Wosilait et al. [27] constructed an empirical model for human embryonic and fetal growth during in utero development that was based on four data sets covering various periods of gestation that were published between 1909 and 1975.
- Luecke et al. [15] published a general human pregnancy PBPK model that uses the fetal mass model of Wosilait et al. [27]. In the context of the PBPK model, Luecke et al. [15] provided a collection of models for changes in maternal and fetal tissue volumes and blood flow rates that are based upon allometric scaling of the fetal mass, but they did not describe the methods and data that were used to calibrate these models. Those authors cited a “submitted” article (“Luecke et al., ‘93b”) as the source of the models, but the manuscript of the same title that was ultimately published [25] only described the models and data sources for time-varying masses of fetal tissues (not for fetal blood flow rates, maternal tissue masses, or maternal blood flow rates).
- Abduljalil et al. [28] and Gaohua et al. [29] described empirical models for many of the anatomical and physiological changes that occur in a human mother during pregnancy. For their maternal models, these authors compiled an extensive list of published studies to build a large combined data set for various anatomical and physiological changes that occur in human, singleton, low-risk, normal pregnancies. Abduljalil et al. [28] also developed a model for fetal volume and Gaohua et al. [29] constructed models for the volume of and blood flow rates to a “fetoplacental unit”, which they consider as a single lumped compartment in their human pregnancy PBPK model. For each quantity of interest, Abduljalil et al. [28] used the coefficient of determination (*R*^2^) to select an optimal model from among several polynomial models, but using this measure of goodness tends to bias the model selection process by favoring models with more parameters [30].
- Dallmann et al. [3] curated a data set and used it to fit models for anatomical and physiological changes that occur in women and their fetuses during pregnancy. In this analysis, the authors considered changes to fetal mass, cord blood flow, and hematocrit, but they did not provide formulae for other fetal changes. In contrast to models constructed by others, the models of Dallmann et al. [3] use “fetal age” (i.e., time since fertilization of the ovum) instead of “gestational age” (time since the first day of the final menstrual period prior to conception) as the independent variable.
- Abduljalil et al. [31] described empirical models for organ and tissue volumes of a human fetus during gestation. To construct the models, these authors compiled a data set from published studies on embryonic and fetal tissue growth and composition during in utero development. Like Dallmann et al. [3], but in contrast to their own previous work [28], these authors used fetal age rather than gestational age as the independent variable in their models.

Here we have expanded on previous work by constructing empirical models for anatomical and physiological changes in a human mother and fetus that either (1) have not been previously described mathematically in the literature; (2) have been described mathematically, but by models that appear to give unreasonable values; or (3) have been described mathematically, but using models that rely on data that are sparse or outdated. Wherever possible, we have endeavored to compare our models to those presented elsewhere. Also, we include herein mathematical descriptions for “rest of body” volumes and blood flow rates that incorporate mass balance principles, whereas the aforementioned published collections do not. These efforts have resulted in a comprehensive collection of empirical models for important time-varying quantities that one should consider when constructing a PBPK model of human gestation and pregnancy.

## METHODS

### Data Sets

We used both curated (multiple-source) data sets and original (single-source) data sets to calibrate empirical models for various anatomical and physiological quantities that vary during gestation. In particular, we relied heavily on composite data sets published by Abduljalil et al. [28] and Abduljalil et al. [31]; for those quantities not described in these two sources (e.g., fetal blood flow rates), we located data from other published studies. Also, for cases in which data were only available in a graphical form, we used the WebPlotDigitizer data extraction tool [32] to convert the graphically-presented data into numerical data. The sources for the specific data sets used to calibrate models for each quantity of interest are identified in Table 1, and additional details are given in the Results section.

**Table 1.**
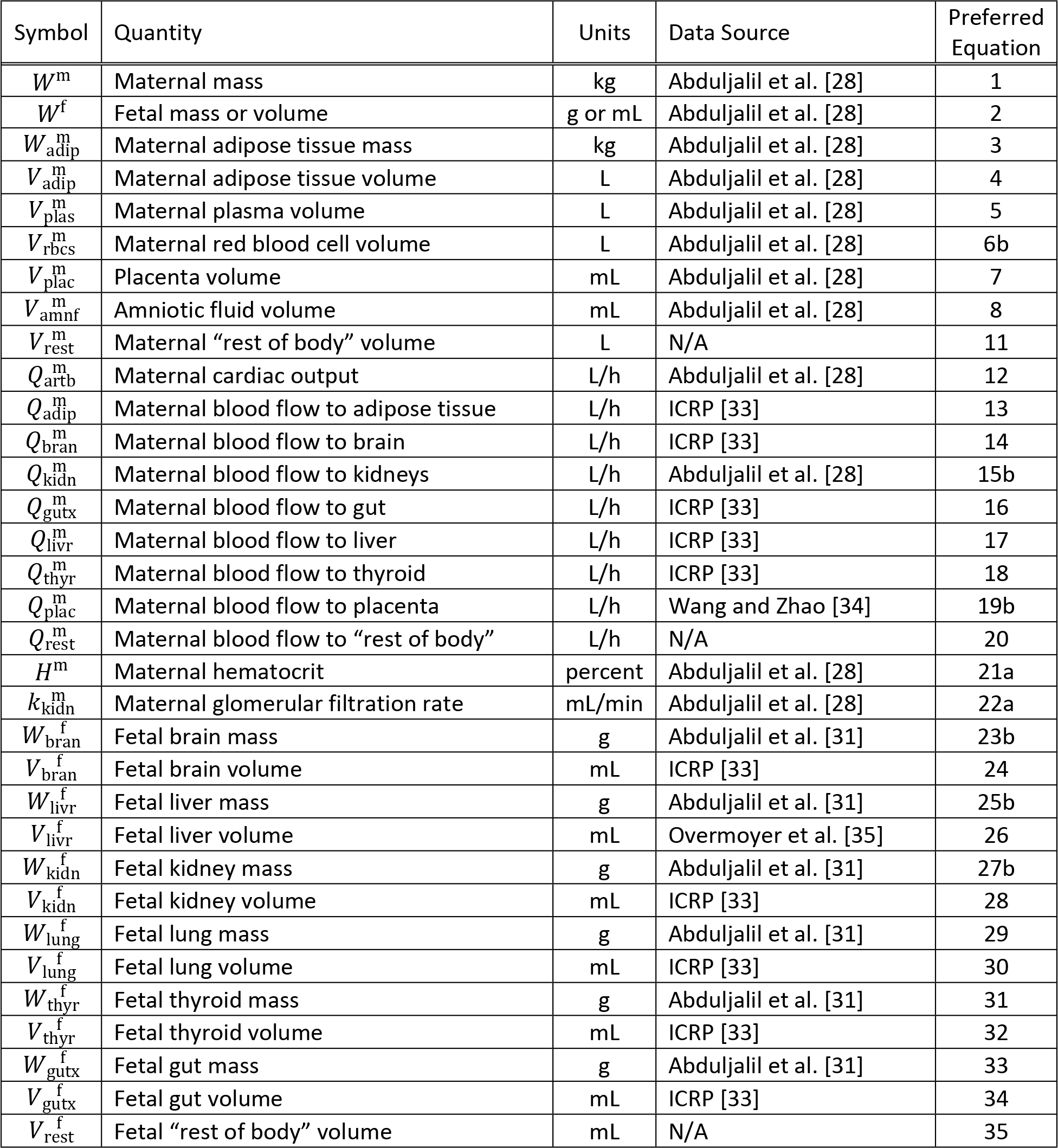
Data sources and preferred models for various quantities of interest.

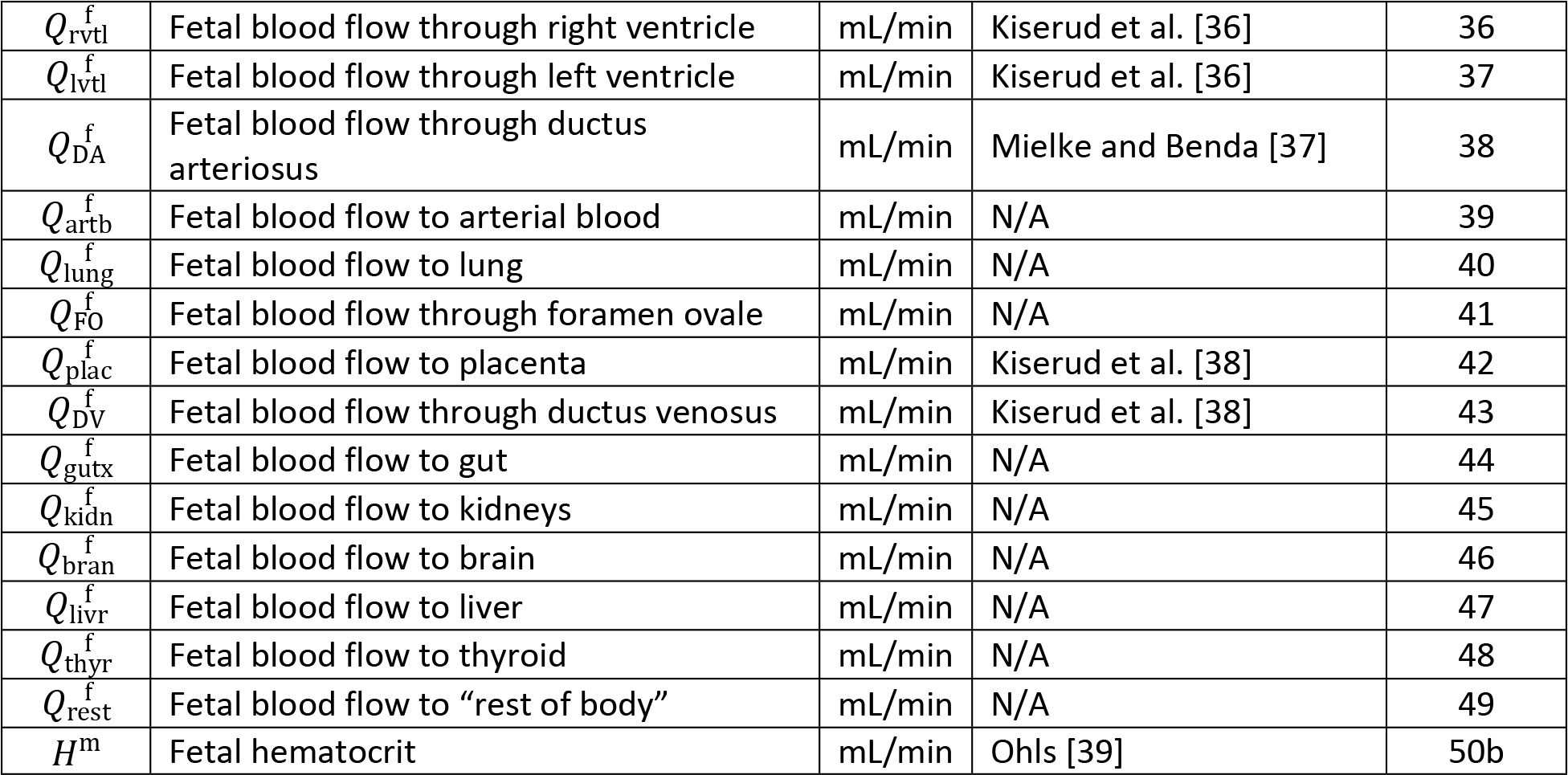

Some data sets are based on gestational age, or time since the last menstruation, whereas others are based on fetal age, or time since fertilization of the ovum. For purposes data analysis and model comparison, we assumed that gestational age equals fetal age plus two weeks [40] to convert data and models based on fetal age to time scales based on gestational age.

### Models

We examined four basic types of models to describe changes in masses, volumes, percentages (such as hematocrits) and rates (such as blood flow and glomerular filtration rates) that occur during the singleton pregnancy of an “average” healthy woman. In particular, we considered polynomial models (up to degree 3), Gompertz models, logistic models, and allometric power law models. For quantities expected to have initial values (i.e., values at the beginning of gestation) of zero (such as fetal body mass), we examined pure Gompertz and logistic growth models as candidate models, whereas for quantities expected to have initial values substantially greater than zero (such as maternal body mass) we examined modified Gompertz and logistic growth models that include an additional parameter describing the initial value. For some of the quantities considered, power law models were used to relate the quantity to a body mass (i.e., the maternal or fetal mass). When the quantity in question had an initial value substantially greater than zero but the related body mass did not (as for fetal body mass), we examined modified power law models that include an additional parameter describing the initial value. Formulas and references for all models we considered are provided in Table 2.

**Table 2.**
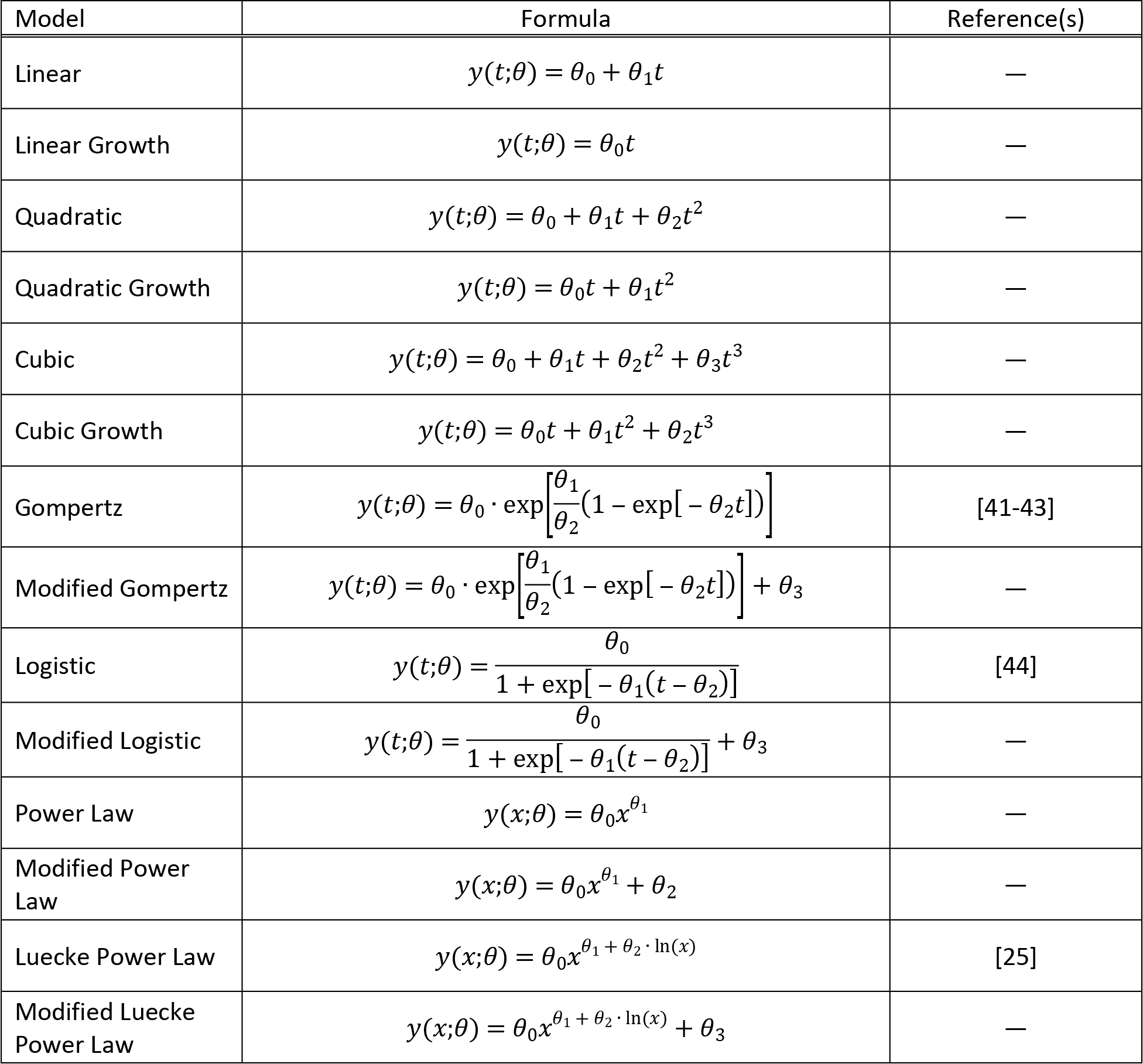
Types of models used to describe changes in masses, volumes, percentages, and physiological rates during pregnancy and gestation.

| Model | Formula | Reference(s) |
| --- | --- | --- |
| Linear | $y(t;\theta) = \theta_0 + \theta_1 t$ | — |
| Linear Growth | $y(t;\theta) = \theta_0 t$ | — |
| Quadratic | $y(t;\theta) = \theta_0 + \theta_1 t + \theta_2 t^2$ | — |
| Quadratic Growth | $y(t;\theta) = \theta_0 t + \theta_1 t^2$ | — |
| Cubic | $y(t;\theta) = \theta_0 + \theta_1 t + \theta_2 t^2 + \theta_3 t^3$ | — |
| Cubic Growth | $y(t;\theta) = \theta_0 t + \theta_1 t^2 + \theta_2 t^3$ | — |
| Gompertz | $y(t;\theta) = \theta_0 \cdot \exp\left[\frac{\theta_1}{\theta_2}(1 - \exp[-\theta_2 t])\right]$ | [41-43] |
| Modified Gompertz | $y(t;\theta) = \theta_0 \cdot \exp\left[\frac{\theta_1}{\theta_2}(1 - \exp[-\theta_2 t])\right] + \theta_3$ | — |
| Logistic | $y(t;\theta) = \frac{\theta_0}{1 + \exp[-\theta_1(t - \theta_2)]}$ | [44] |
| Modified Logistic | $y(t;\theta) = \frac{\theta_0}{1 + \exp[-\theta_1(t - \theta_2)]} + \theta_3$ | — |
| Power Law | $y(x;\theta) = \theta_0 x^{\theta_1}$ | — |
| Modified Power Law | $y(x;\theta) = \theta_0 x^{\theta_1} + \theta_2$ | — |
| Luecke Power Law | $y(x;\theta) = \theta_0 x^{\theta_1 + \theta_2 \cdot \ln(x)}$ | [25] |
| Modified Luecke Power Law | $y(x;\theta) = \theta_0 x^{\theta_1 + \theta_2 \cdot \ln(x)} + \theta_3$ | — |

### Model Calibration

In considering any particular model and any given quantity of interest (e.g., maternal body mass), we used data to identify the model parameters {*θ*_0_,*θ*_1_,…} that allow one to estimate the quantity of interest as a function *y* of an independent variable *t* (e.g., a gestational age or body mass) and said parameters (cf. Table 2). The data sets utilized included means, standard deviations, and sample sizes paired with times (gestational ages) or body masses. For data sets in which standard deviations and samples sizes were unavailable, we assumed a 20% coefficient of variation and a sample size of one for each data point. We used a standard maximum likelihood approach to obtain a maximum likelihood estimate (MLE) for the model parameters. Further details are provided in the Supporting Information (S1 Appendix).

### Model Selection

After obtaining the MLE for each model for a given data set and quantity of interest, we chose the most parsimonious model by applying the Akaike information criterion (AIC) [45]. We computed the AIC score of each model as

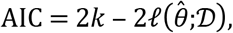

where *k* is the number of parameters in the model, 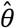 is the MLE, 𝒟 represents the data, and ℓ denotes the log-likelihood function. The model with the lowest AIC score was then selected as the most parsimonious model. For those cases in which the most parsimonious model gave infeasible values for the quantity of interest (e.g., negative values for a mass) at any point in the domain of applicability (e.g., for gestational ages from 0 to 42 weeks), the model with the next lowest AIC and that did not produce infeasible values was identified as the “preferred” model.

### Composite Models

For some important anatomical and physiological quantities, raw data expressed as values vs. gestational ages were not available. For such quantities (e.g., maternal blood flow to adipose tissue and fetal blood flow through the foramen ovale), we constructed composite models. That is, in each such case, we used information about relationships between the quantity of interest and other modeled quantities, possibly along with proportionality constants or relative percentages, to construct a model for the quantity of interest.

### Programming Details

For all analyses described herein, we used Python 3.6.4 with the NumPy (version 1.13.3), SciPy (version 0.19.1), Matplotlib (version 2.1.2), and PIL (version 5.0.0) packages. Scripts are available in the Supporting Information.

### Naming Conventions

We have used mathematical symbols to denote various quantities of interest throughout this manuscript. For example, 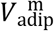 represents the volume of the adipose tissue in the mother. In general, a superscript on such a symbol will be “m” in the case a maternal quantity and “f” in the case of a fetal quantity. When present, the subscript on such a symbol is typically a four-letter code and it indicates a particular physiological compartment. A list of compartment codes is provided in Table 3. For symbols representing blood flow rates through temporary blood vessels or routes in the fetus, the subscript is an upper-case two-letter code indicating the specific blood vessel or route. In particular, “DA”, “DV”, and “FO” indicate the ductus arteriosus, ductus venosus, and foramen ovale, respectively.

**Table 3.** Four-letter codes used to represent compartments in a human mother or fetus. These codes appear as subscripts in mathematical symbols throughout this manuscript.

| Code | Compartment |
| --- | --- |
| artb | Arterial Blood |
| venb | Venous Blood |
| plas | Plasma |
| rbc | Red Blood Cells |
| adip | Adipose |
| lung | Lungs |
| thyr | Thyroid |
| kidn | Kidneys |
| gutx | Gut |
| livr | Liver |
| plac | Placenta |
| amnf | Amniotic Fluid |
| bran | Brain |
| ratm | Right Atrium |
| rvtl | Right Ventricle |
| latm | Left Atrium |
| lvtl | Left Ventricle |
| rest | Rest of Body |

## RESULTS

For each quantity of interest discussed in this section, we describe: (1) the data we used for model calibration, (2) the model selected by us after considering both parsimony and plausibility, and (3) previously published models for the same quantity of interest. Our recommended, or “preferred”, models for all quantities of interest are identified by equation number in Table 1.

### Maternal Mass

To identify an optimal model for the mass of an average woman throughout gestation, we used data curated by Abduljalil et al. [28] to calibrate linear, quadratic, and cubic functions (i.e., polynomial functions of degree 1, 2, and 3), as well as modified Gompertz and logistic functions. The cubic model, which had the lowest AIC, gives the mass (kg) of an average pregnant woman as

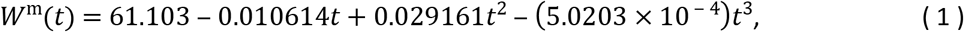

where *t* is the gestational age (weeks) of her fetus. Table 4 shows the maximum likelihood estimates of the parameter values for all models considered along with the associated log-likelihood and AIC values. The cubic model of Equation 1a is shown in Figure 1. *It is important to note that “maternal mass” includes the mass of the fetus, the placenta, and the amniotic fluid*.

**Table 4.** Maternal mass models (mass in kg vs. gestational age in weeks). For each model considered, the maximum likelihood parameter estimates 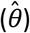, log-likelihood 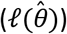, and AIC are provided. The row describing the selected model is shown in boldface.

| Model | $\hat{\theta}_0$ | $\hat{\theta}_1$ | $\hat{\theta}_2$ | $\hat{\theta}_3$ | $\ell(\hat{\theta})$ | AIC |
| --- | --- | --- | --- | --- | --- | --- |
| Linear | 61.093 | 0.35761 | — | — | -30314.7 | 60633.4 |
| Quadratic | 61.019 | 0.42287 | $-1.6429 \times 10^{-3}$ | — | -30312.4 | 60630.8 |
| **Cubic** | **61.103** | **-0.010614** | **0.029161** | **$5.0203 \times 10^{-4}$** | **-30298.0** | **60603.9** |
| Modified Gompertz | 0.078155 | 0.54404 | 0.10286 | 61.023 | -30299.2 | 60606.5 |
| Modified Logistic | 15.780 | 0.14502 | 19.055 | 60.166 | -30298.8 | 60605.6 |

**Figure 1.**
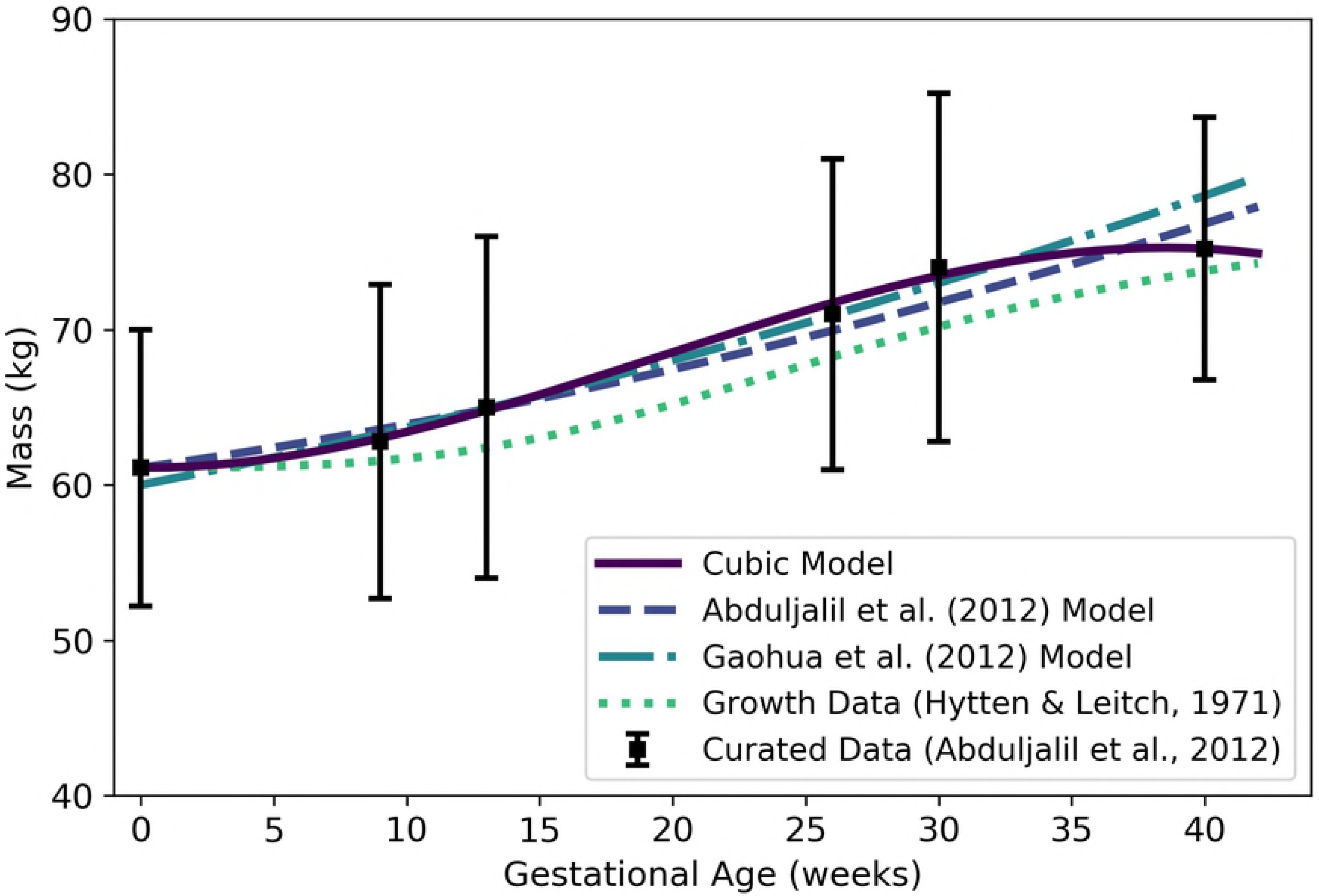
Mass of a human mother vs. gestational age. The cubic model (solid line) given by Equation 1 was selected as the most parsimonious model in our analysis. The models of Abduljalil et al. [28] and Gaohua et al. [29], both quadratic, were calibrated using the same curated data set [28] used by us. The maternal mass gain data (or “growth data”) depicted here were modified from the source [46] to account for an assumed initial mass as described in the text. Note that all models depicted here describe the mass of the entire maternal body plus the products of conception (including the fetus, placenta, and amniotic fluid).

Abduljalil et al. [28] and Gaohua et al. [29] both selected quadratic models to give the mass (in kg) of a pregnant woman as a function of gestational age *t* (in weeks). In both cases, the authors used the same maternal mass data used by us [28] in order to obtain their models. These two models are also shown in Figure 1.

Even without considering the starting mass of a woman, there is tremendous variability in mass *gain* trends of women during pregnancy. For example, some women have gained 23 kg or more at term, while others have actually experienced a *reduction* in total body mass [47]. Hytten and Leitch [46] depicted data summarizing the mean mass gain during pregnancy of 2868 normotensive primigravidae (i.e., women who were pregnant for the first time and who maintained blood pressure in the normal range throughout their pregnancies). Unfortunately, the actual data are not provided by Hytten and Leitch [46], so we were unable to calibrate models to that data as discussed in the Methods section. To compare the trend shown by Hytten and Leitch [46] with our model and the models of Abduljalil et al. [28] and Gaohua et al. [29], we captured data points from their graphical depiction of the data trend curve, calibrated a Gompertz growth curve to fit those points, then added an initial mass of 61.1 kg to match the initial mass of the Abduljalil et al. [28] model. The resulting curve is shown for purposes of comparison in Figure 1.

### Fetal Mass

To identify an optimal model for the *volume* of an average human fetus throughout gestation, we used data curated by Abduljalil et al. [28]. The Gompertz model, which had the lowest AIC, gives the volume (mL) of an average human fetus as

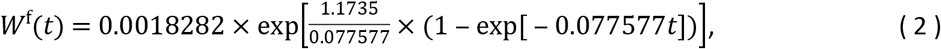

where *t* is the gestational age (weeks). Since the average density of a human fetus is approximately 1 g/mL throughout gestation [48], *W*^f^(*t*) also represents the *mass* (g) of the fetus at a given gestational age. Table 5 shows the maximum likelihood estimates of the parameter values for all models considered along with the associated log-likelihood and AIC values. Figure 2 shows the Gompertz model of Equation 2 and three published models for human fetal mass [3,27,28].

**Table 5.** Fetal mass models (mass in g vs. gestational age in weeks). For each model considered, the maximum likelihood parameter estimates 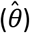, log-likelihood 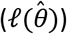, and AIC are provided. The row describing the selected model is shown in boldface.

| Model | $\hat{\theta}_0$ | $\hat{\theta}_1$ | $\hat{\theta}_2$ | $\hat{\theta}_3$ | $\ell(\hat{\theta})$ | AIC |
| --- | --- | --- | --- | --- | --- | --- |
| Linear Growth | 2.0609 | — | — | — | -1.3×10<sup>6</sup> | 2.6×10<sup>6</sup> |
| Quadratic Growth | -14.715 | 2.4326 | — | — | -288832 | 577668 |
| Cubic Growth | 0.11563 | 0.061459 | 0.045199 | — | -285330 | 570666 |
| **Gompertz** | **0.0018282** | **1.1735** | **0.077577** | — | **-258093** | **516193** |
| Logistic | 3680.3 | 0.29875 | 31.205 | — | -259854 | 519714 |

**Figure 2.**
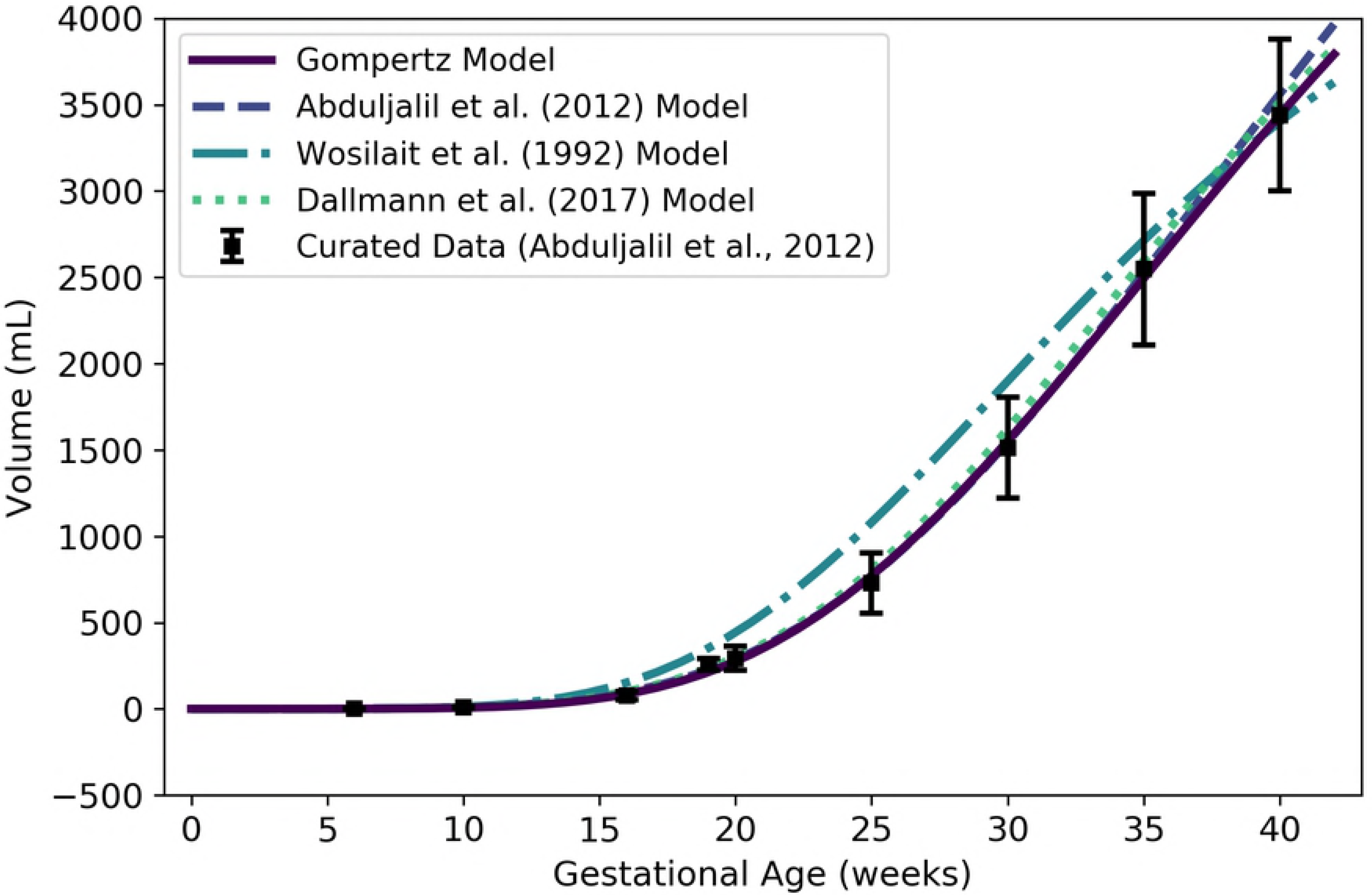
Volume (mL) or mass (g) of a human fetus vs. gestational age. The Gompertz model (solid line) given by Equation 2 was selected as the most parsimonious model in our analysis. The models of Abduljalil et al. [28] and Wosilait et al. [27] were also Gompertz models, though the model parameters used by those authors were different. Dallmann et al. [3] used a log-logistic model for fetal volume. The model of Abduljalil et al. [28] was calibrated using the same curated data set [28] used by us, while the models of Wosilait et al. [27] and Dallmann et al. [3] were calibrated using different data sets.

Like us, Abduljalil et al. [28] and Wosilait et al. [27] both selected Gompertz models to give the volume (mL) of a human fetus as a function of gestational age *t* (weeks). The model of Abduljalil et al. [28] was calibrated using the same curated data set [28] used by us, but we were unable to reproduce the Gompertz model parameters reported by them. Nevertheless, as illustrated in Figure 2, our Gompertz model and the Abduljalil et al. [28] model predict similar fetal volumes for most gestational ages. The model of Wosilait et al. [27], also depicted in Figure 2, was calibrated using a different data set; thus, it is not surprising that they obtained different Gompertz model parameters and different predicted fetal volumes. Dallmann et al. [3] used a log-logistic model for fetal volume that was calibrated with yet another curated data set.

### Maternal Compartment Volumes

While volumes of many maternal organs and tissues remain approximately constant throughout pregnancy, several of these undergo dramatic changes. In particular, the volumes of adipose tissue and blood components (plasma and red blood cells) generally increase as pregnancy progresses. In addition, the “products of conception”, which include the placenta and the amniotic fluid, increase in size as pregnancy progresses and therefore contribute to maternal mass gain [46].

#### Adipose Tissue

For the *mass* of the adipose (fat) tissue of an average woman throughout pregnancy, we used data curated by Abduljalil et al. [28] to calibrate various models. In calibrating the polynomial models and the modified Gompertz and logistic models, we used the total maternal fat mass (mean and standard deviation) vs. gestational age data exactly as tabulated by Abduljalil et al. [28]. In an alternative approach, we followed the working assumption of Luecke et al. [15] that maternal fat mass gain correlates with fetal mass; that is, we considered allometric relationships between these two quantities. Using Equation 2, we calculated a fetal mass for each gestational age data point that was tabulated [28] for total maternal fat mass. We then calibrated modified versions of the power law and Luecke power law models (cf. Table 2) using data for the total maternal fat mass (mean and standard deviation) vs. (calculated) fetal mass. Of all models considered, the linear model had the lowest AIC. This model gives the mass (kg) of the adipose tissue of an average pregnant woman as

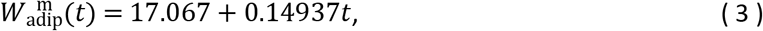

where *t* is the gestational age (weeks). Since the mean density of human adipose tissue is 0.950 kg/L [49], the *volume* of the maternal adipose tissue (L) can be computed as

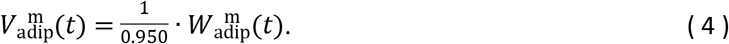

Table 6 shows the maximum likelihood estimates of the parameter values for all models considered along with the associated log-likelihood and AIC values. The linear model of Equation 3 and several other models for maternal fat mass are shown in Figure 3.

**Table 6.** Maternal fat mass models (mass in kg vs. fetal mass in kg for power law models, mass in kg vs. gestational age in weeks for all other models). For each model considered, the maximum likelihood parameter estimates 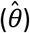, log-likelihood 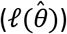, and AIC are provided. The row describing the selected model is shown in boldface.

| Model | $\hat{\theta}_0$ | $\hat{\theta}_1$ | $\hat{\theta}_2$ | $\hat{\theta}_3$ | $\ell(\hat{\theta})$ | AIC |
| --- | --- | --- | --- | --- | --- | --- |
| **Linear** | **17.067** | **0.14937** | — | — | **-3221.24** | **6446.48** |
| Quadratic | 17.249 | 0.11990 | $7.2110 \times 10^{-4}$ | — | -3221.13 | 6448.26 |
| Cubic | 17.165 | 0.19050 | -0.0046448 | $9.6148 \times 10^{-5}$ | -3220.99 | 6449.98 |
| Modified Gompertz | 20.402 | 0.0058803 | -0.0042400 | -3.1426 | -3221.12 | 6450.24 |
| Modified Logistic | 104.49 | 0.0070722 | 152.86 | -9.3414 | -3221.18 | 6450.36 |
| Modified Power Law | 4.0204 | 0.26566 | 17.224 | — | -3221.90 | 6449.80 |
| Modified Luecke Power Law | 4.3011 | 0.25439 | 0.0064128 | 16.904 | -3221.77 | 6451.54 |

**Figure 3.**
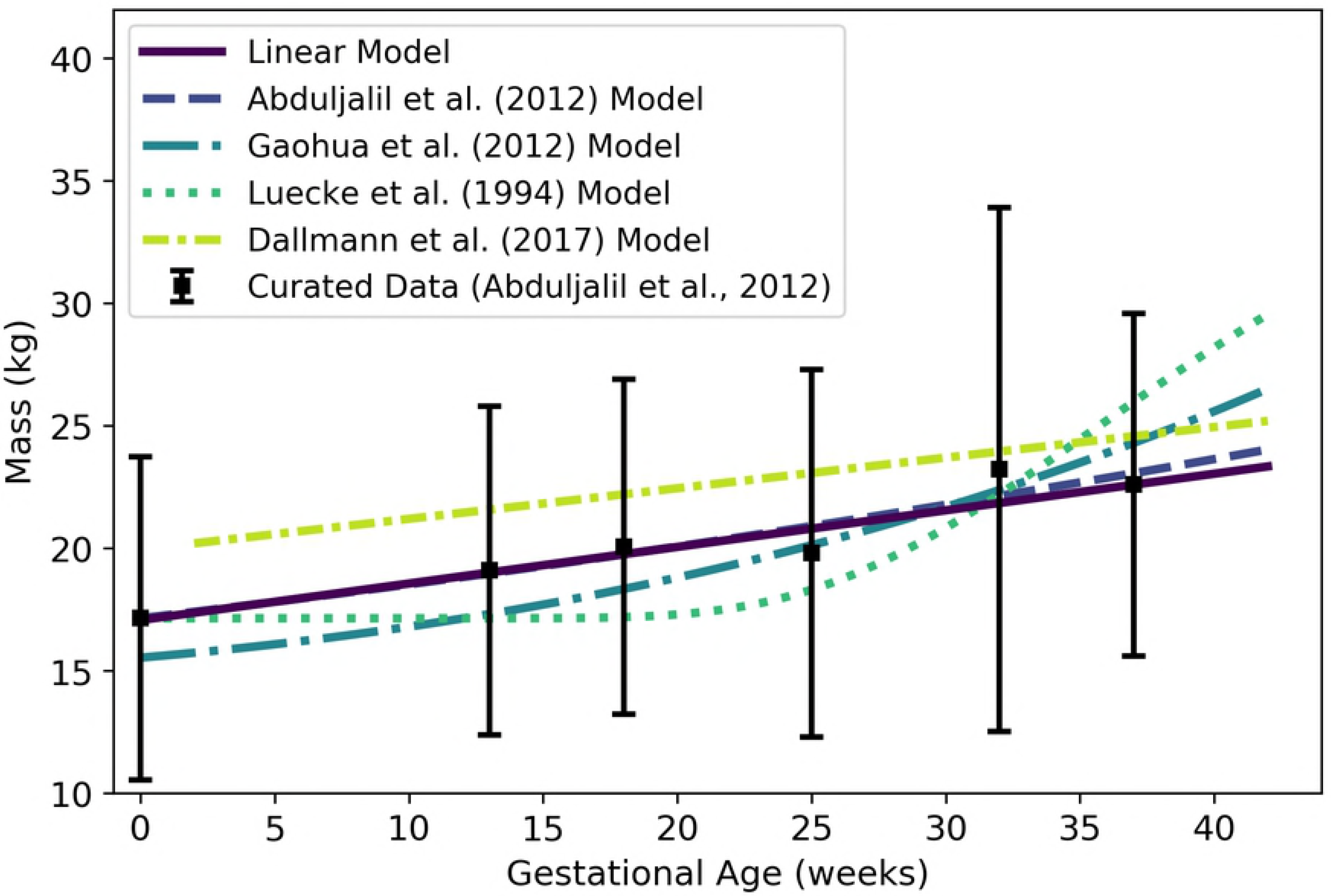
Adipose tissue mass of a human mother of vs. gestational age. The linear model (solid line) given by Equation 3 was selected as the most parsimonious model in our analysis. The models of Abduljalil et al. [28] and Gaohua et al. [29], both quadratic, were calibrated using the same curated data set [28] used by us. The latter of these models was modified as described in the text. The model of Luecke et al. [15] was calibrated using different data. It predicts maternal fat mass as a function of total fetal mass, and was interpreted as described in the text. Dallmann et al. [3] also selected a linear model, but they calibrated their model with different data.

Abduljalil et al. [28] and Gaohua et al. [29] each selected quadratic models to give the mass (kg) of adipose tissue in a pregnant woman as a function of gestational age *t* (weeks). In both cases, the authors used the same maternal total fat mass data used by us [28] to obtain their models. Gaohua et al. [29] claim that their model gives the total fat mass (kg) vs. gestational age, but the masses predicted for various gestational ages are about 50% larger than the mean data values upon which the model is based. In an effort to explain this discrepancy, we hypothesized that the Gaohua et al. [29] quadratic model for maternal fat mass is actually a model for the *percentage* of the mother’s body mass that is fat. Thus, to obtain fat mass values for comparison, we divided the values predicted by their model by 100 and then multiplied the resulting values by corresponding values from the maternal mass vs. gestational age model that these authors provide in the same manuscript [29]. That is, in Figure 3, the model labeled “Gaohua et al. [29] Model” differs from the model described by those authors.

Luecke et al. [15] constructed a model for maternal fat mass *gain*, so to compare its predictions with those of the other models, we added an initial mass of 17.14 kg (the mean value for non-pregnant women from the data set curated by Abduljalil et al. [28]) to the predicted mass gain values. The model of Luecke et al. [15], modified as just described to account for an initial maternal fat mass, is shown in Figure 3. For earlier gestational ages, this model predicts maternal fat masses considerably lower than the data values curated by Abduljalil et al. [28]. This may be because the model is a power law based on fetal mass, which does not increase substantially (from an initial mass of about zero) until about 15 weeks, whereas the data suggest that maternal fat mass *does* begin to increase substantially between conception and 13 weeks.

Note that the total fat mass predicted (by our preferred linear model) for a woman at conception, 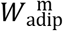 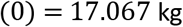, represents about 27.9% of the total body mass, *W*^m^(0) = 61.103 kg. This represents a considerable deviation from the figure of 37.5% reported in Table 7 for adipose body mass percentage for women, but note that that number represents an average for *all* women [33]. The individuals that have been included in studies of pregnancy [3,28], appear to have a lower mean body fat percentage than the population of all women, or even all women of childbearing age [50].

**Table 7.** Percent of total body mass and density (references listed) for various human organs and tissues.

| Compartment | Code | Percent (%) of Total Body Mass* |  | Density (kg/L) | Density Reference |
| --- | --- | --- | --- | --- | --- |
|  |  | Male | Female<sup>†</sup> |  |  |
| Adipose | adip | 19.9 | 37.5 | 0.95 | Martin et al. [49] |
| Brain | bran | 1.99 | 2.17 | 1.04 | ICRP [33] |
| Thyroid | thyr | 0.0274 | 0.0283 | 1.05 | ICRP [33] |
| Kidneys | kidn | 0.425 | 0.458 | 1.05 | ICRP [33] |
| Gut<sup>‡</sup> | gutx | 1.66 | 1.90 | 1.045 | ICRP [33] |
| Liver | livr | 2.47 | 2.33 | 1.05 | Overmoyer et al. [35] |
| Lungs | lung | 1.64 | 1.58 | 1.05 | ICRP [33] |
| Arterial Blood | artb | — | — | 1.06 | ICRP [33] |
| Venous Blood | venb | — | — | 1.06 | ICRP [33] |
| Placenta | plac | — | — | 1.02 | Del Nero et al. [51] |
| Amniotic Fluid | amnf | — | — | 1.01 | Uyeno [52] |

Percentages were computed using reference masses from Table 2.8 of ICRP (2002).
<sup>†</sup>Percentages listed in this column are for a non-pregnant woman.
<sup>‡</sup>The values listed for the “Gut” assume this compartment comprises the esophagus, stomach, small intestine, and large intestine.

#### Plasma

Plasma is one of the two major constituents of blood. We used the curated data of Abduljalil et al. [28] to calibrate various models for plasma volume in a human mother during pregnancy. The modified logistic model, which we found to be the best of the candidate models based on AIC, gives the plasma volume (L) of the mother as

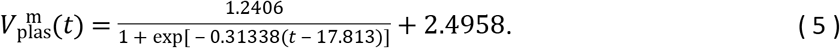

where *t* is the gestational age (weeks). Table 8 shows the maximum likelihood estimates of the parameter values for all models considered along with the associated log-likelihood and AIC values. We remark that the modified Luecke power law model yielded a lower AIC (365.07 vs. 365.13), but the AIC difference is too small to recommend one model over the other. Furthermore, the modified Luecke power law relates maternal plasma volume to fetal mass whereas the modified logistic model relates maternal plasma volume directly to gestational age; thus, the latter model has the advantage of not requiring an (intermediate) estimate of fetal mass at each time point of interest. Figure 4 shows the modified logistic model of Equation 5 and several other models for maternal plasma volume. (Figure 4 does not show the modified Luecke power law model, but values predicted by that model and the modified logistic model differ by less than 2% throughout the period from 0 to 42 gestational weeks.)

**Table 8.**
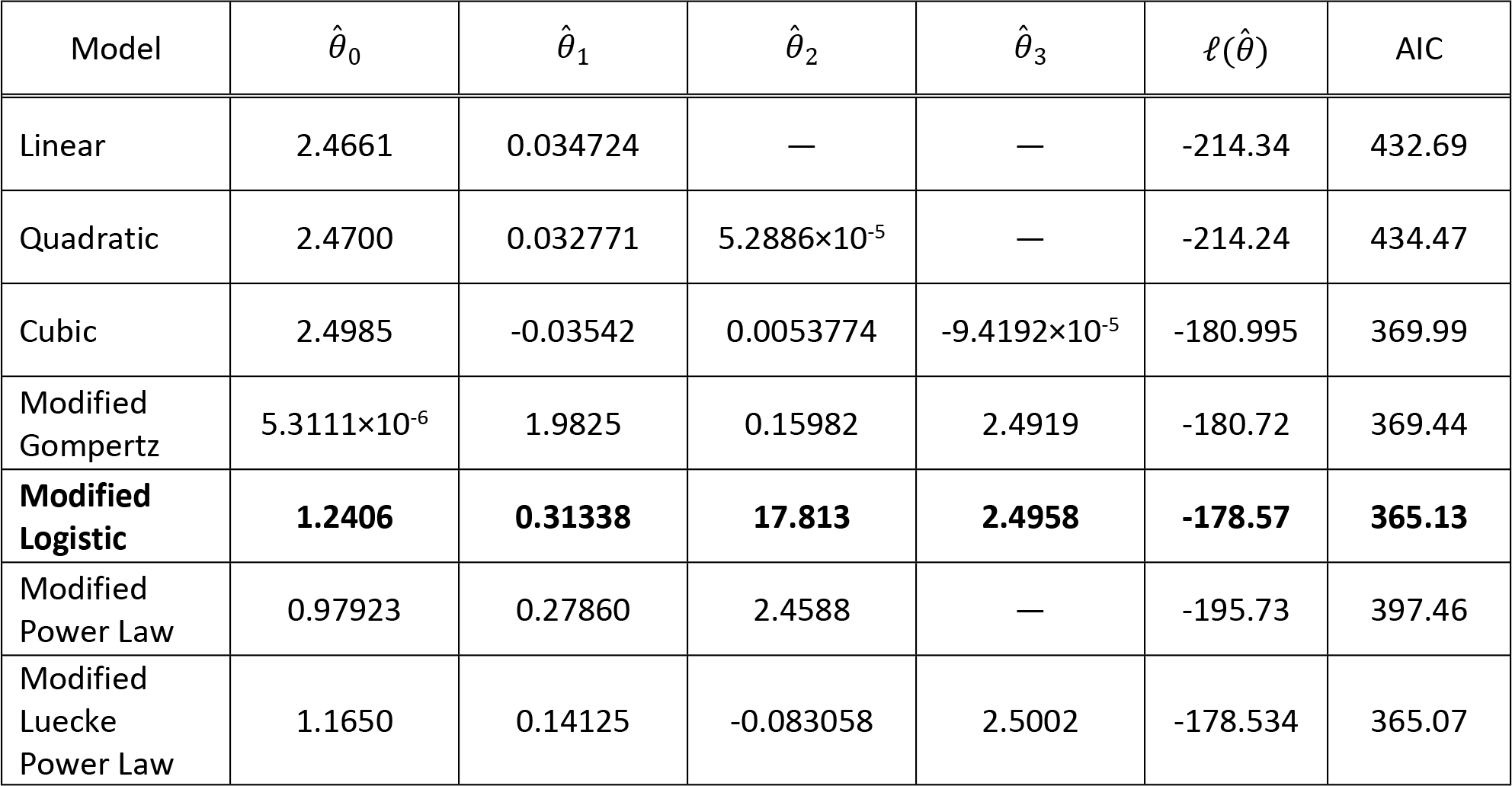
Maternal plasma volume models (volume in L vs. fetal mass in kg for power law models, volume in L vs. gestational age in weeks for all other models). For each model considered, the maximum likelihood parameter estimates 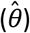, log-likelihood 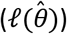, and AIC are provided. The row describing the selected model is shown in boldface.

**Figure 4.**
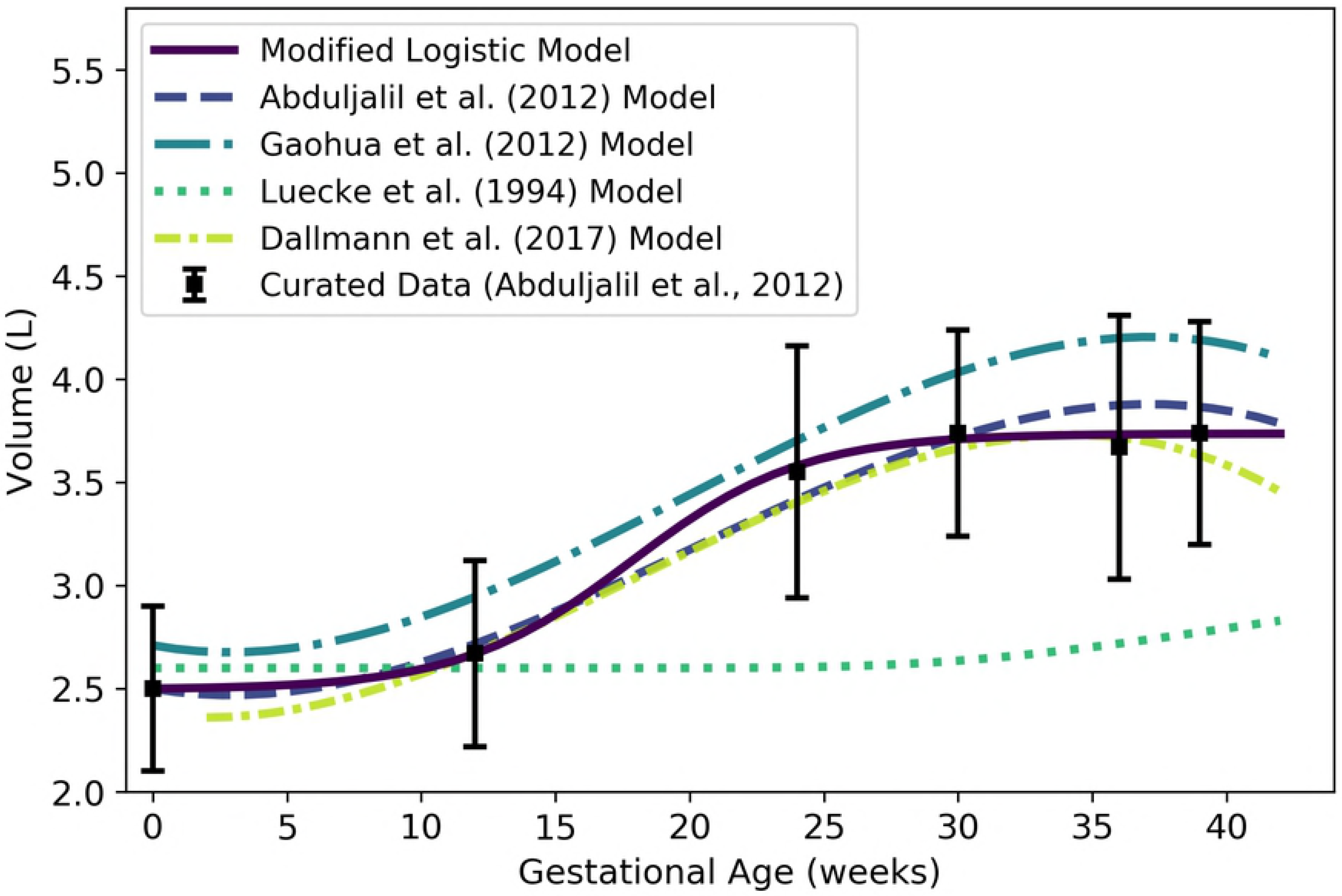
Maternal plasma volume vs. gestational age. The modified logistic model (solid line) given by Equation 5 was selected as the most parsimonious model in our analysis. The models of Abduljalil et al. [28] and Gaohua et al. [29], both cubic polynomials, were calibrated using the same curated data set [28] used by us. The model of Luecke et al. [15] was calibrated using different data. It assumes an initial maternal plasma volume of 2.6 L (or an initial plasma mass of 2.6 kg) and predicts an increase in maternal plasma volume (or mass) as a function of total fetal mass. Dallmann et al. [3] calibrated their cubic model with different data.

#### Red Blood Cells

Red blood cells (RBCs) make up the other major component of human blood. We used curated data from Abduljalil et al. [28] to calibrate various models for RBC volume in a human mother during pregnancy. We selected the modified logistic model given by

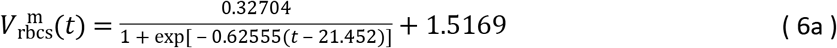

as the most parsimonious of the candidate models. Here, 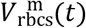 denotes the volume (L) of the RBCs at gestation age *t* (weeks). Table 9 shows the maximum likelihood estimates of the parameter values for all models considered along with the associated log-likelihood and AIC values.

**Table 9.** Maternal RBC volume models (volume in L vs. fetal mass in kg for power law models, volume in L vs. gestational age in weeks for all other models). For each model considered, the maximum likelihood parameter estimates 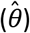, log-likelihood 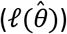, and AIC are provided. The row describing the selected model is shown in boldface.

| Model | $\hat{\theta}_0$ | $\hat{\theta}_1$ | $\hat{\theta}_2$ | $\hat{\theta}_3$ | $\ell(\hat{\theta})$ | AIC |
| --- | --- | --- | --- | --- | --- | --- |
| Linear | 1.4555 | 0.01101 | — | — | 3597.70 | -7191.41 |
| Quadratic | 1.4741 | 0.007927 | $7.7997 \times 10^{-5}$ | — | 3612.98 | -7219.95 |
| Cubic | 1.4918 | -0.0070658 | 0.0011160 | $-1.7512 \times 10^{-5}$ | 3660.39 | -7312.78 |
| Modified Gompertz | $1.9369 \times 10^{-5}$ | 1.1786 | 0.11792 | 1.4925 | 3679.15 | -7350.31 |
| **Modified Logistic** | **0.32704** | **0.62555** | **21.452** | **1.5169** | **3753.96** | **-7499.93** |
| Modified Power Law | 0.27569 | 0.33366 | 1.4802 | — | 3662.1 | 7318.19 |
| Modified Luecke Power Law | 0.28532 | 0.37478 | -0.18321 | 1.5109 | 3693.36 | -7378.72 |

Through analysis of data sets describing maternal plasma volume, RBC volume, and hematocrit [28], we independently obtained models for each of these quantities (cf. Equations 5, 6a, and 21a). However, as one might expect when independently constructing models of interrelated quantities, the models that arose are not perfectly consistent. Because hematocrit represents the volume percentage of RBCs in whole blood, and because whole blood is mostly made up of plasma and RBCs (with only a small fraction made up of white blood cells and platelets), we can estimate the volume (L) of RBCs in maternal blood as

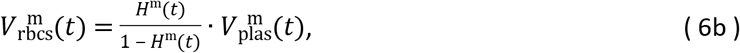

where *H*^m^(*t*) and 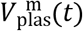 represent the maternal hematocrit and plasma volume (L) at gestation age *t* (weeks). Thus, if one uses the models for maternal plasma volume, RBC volume, and hematocrit given by Equations 5, 6b, and 21a, respectively, the model predictions will be consistent with one another. The modified logistic model of Equation 6a, the alternate hematocrit-based model of Equation 6b, and three published models for maternal RBC volume [3,28,29] are shown in Figure 5.

**Figure 5.**
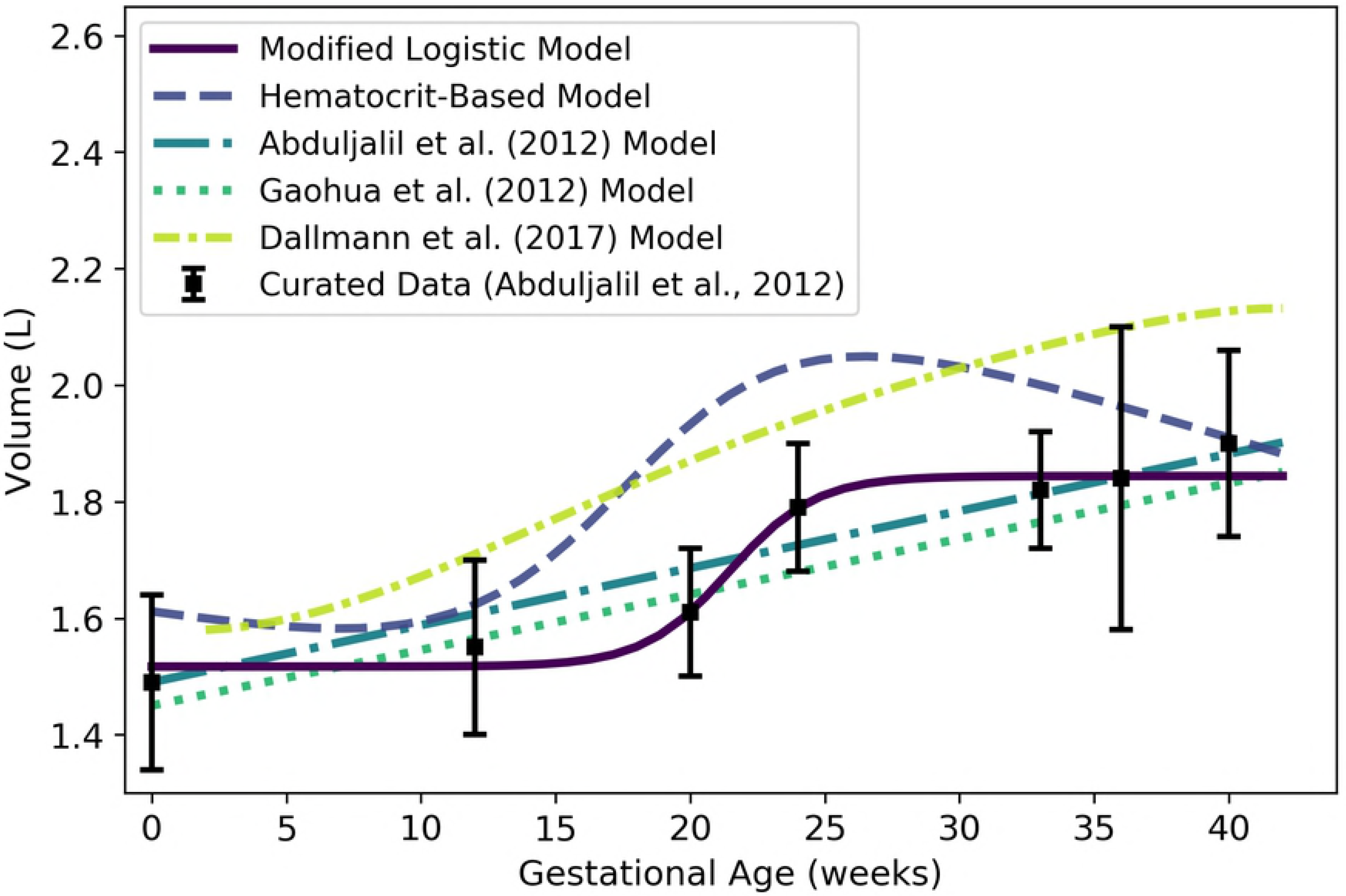
Maternal RBC volume vs. gestational age. The modified logistic model (solid line) given by Equation 6a was selected as the most parsimonious model in our analysis, but the hematocrit-based model (second in legend) of Equation 6b ensures consistency with models for plasma volume (Equation 5) and hematocrit (Equation 21). (See also Figure 4 and Figure 13.) The models of Abduljalil et al. [28] and Gaohua et al. [29], both of which are linear models, were calibrated using the same curated data set [28] used by us. Dallmann et al. [3] did not create a model for maternal RBC volume, so the model attributed to them here is algebraically derived from their models for plasma volume and hematocrit.

#### Placenta

We used the curated data of Abduljalil et al. [28] to calibrate various models for human placenta volume. The cubic growth model given by

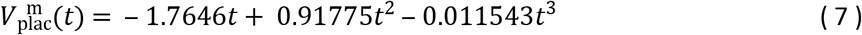

was selected as the most parsimonious model for placenta volume (mL) at gestational age *t* (weeks). Table 10 shows the maximum likelihood estimates of the parameter values for all models considered along with the associated log-likelihood and AIC values. Figure 6 shows the power law model of Equation 7 and three published models for placenta volume [3,15,28].

**Table 10.** Placenta volume models (volume in mL vs. fetal mass in kg for power law models, volume in mL vs. gestational age in weeks for all other models). For each model considered, the maximum likelihood parameter estimates 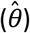, log-likelihood 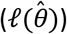, and AIC are provided. The row describing the selected model is shown in boldface.

| Model | $\hat{\theta}_0$ | $\hat{\theta}_1$ | $\hat{\theta}_2$ | $\hat{\theta}_3$ | $\ell(\hat{\theta})$ | AIC |
| --- | --- | --- | --- | --- | --- | --- |
| Linear Growth | 16.395 | — | — | — | -78521.5 | 157045 |
| Quadratic Growth | 8.3802 | 0.20441 | — | — | -78222.9 | 156450 |
| **Cubic Growth** | **-1.7646** | **0.91775** | **-0.011543** | — | **-78114.3** | **156235** |
| Gompertz | 5.4388 | 0.33757 | 0.065182 | — | -78119.3 | 156245 |
| Logistic | 802.77 | 0.12491 | 27.831 | — | -78146.5 | 156299 |
| Power Law | 398.05 | 0.40647 | — | — | -78131.6 | 156267 |
| Luecke Power Law | 389.55 | 0.4129 | 0.0096187 | — | -78125.4 | 156257 |

**Figure 6.**
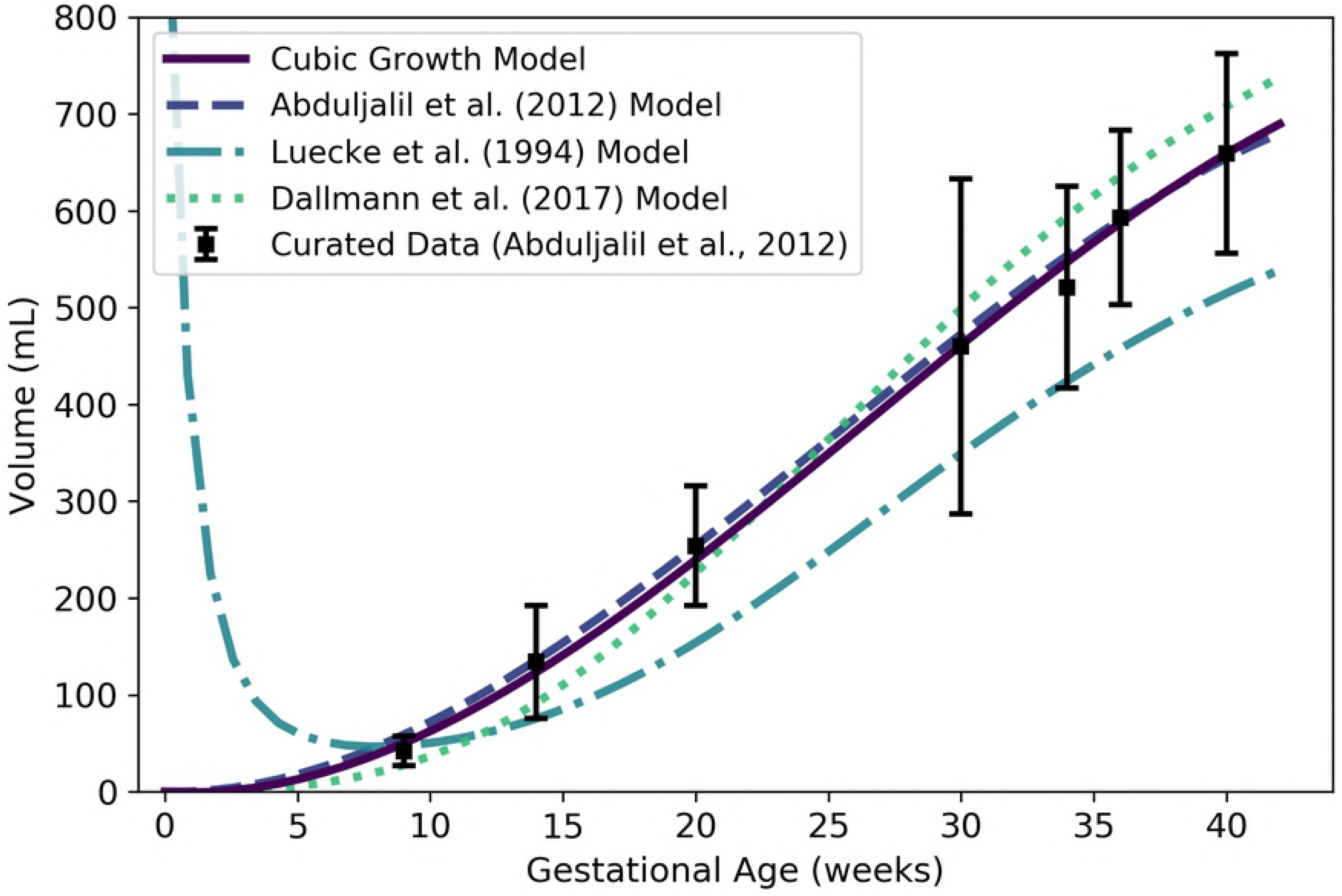
Placenta volume vs. gestational age. The cubic growth model (solid line) given by Equation 7 was selected as the most parsimonious model in our analysis. The model of Abduljalil et al. [28], also a cubic polynomial model, was calibrated using the same curated data set [28] used by us. The model of Luecke et al. [15] was calibrated using different data and assumes a relationship between placenta volume and fetal mass. Dallmann et al. [3] calibrated their cubic model with different data.

It is worth noting that Equation 7 yields negative volumes for the placenta for gestational ages less than about 1.97 weeks; thus, this equation should not be used to estimate placental volume during that time frame. In any case, conception does not occur until a gestational age of about 2 weeks, so neither the placenta nor the embryo, itself, come into existence until after that time. At a fetal age of about 4 days, or a gestational age of about 18 days, the cells in the periphery of the blastocyst (i.e., the early embryo) become distinguishable as a trophoblast; this trophoblast is a precursor to the fetal component of the placenta [40]. We propose that Equation 7 be used to estimate the volume of the placenta after 2 weeks, but it should not be considered accurate until 9 weeks (the time of the first data point of Abduljalil et al. [28]).

#### Amniotic Fluid

We used the curated data of Abduljalil et al. [28] to calibrate various models for human amniotic fluid volume. The logistic model given by

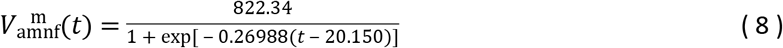

was selected as the most parsimonious model for amniotic fluid volume (mL) at gestational age *t* (weeks). Table 11 shows the maximum likelihood estimates of the parameter values for all models considered along with the associated log-likelihood and AIC values. The logistic model of Equation 8 and three published models for amniotic fluid volume [3,15,28] are shown in Figure 7.

**Table 11.** Amniotic fluid volume models (volume in mL vs. fetal mass in kg for power law models, volume in mL vs. gestational age in weeks for all other models). For each model considered, the maximum likelihood parameter estimates 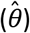, log-likelihood 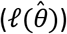, and AIC are provided. The row describing the selected model is shown in boldface.

| Model | $\hat{\theta}_0$ | $\hat{\theta}_1$ | $\hat{\theta}_2$ | $\hat{\theta}_3$ | $\ell(\hat{\theta})$ | AIC |
| --- | --- | --- | --- | --- | --- | --- |
| Linear Growth | 9.3837 | — | — | — | -1122.31 | 2246.62 |
| Quadratic Growth | -0.045477 | 0.61316 | — | — | -824.80 | 1653.6 |
| Cubic Growth | -15.702 | 2.4783 | -0.039285 | — | -759.30 | 1524.61 |
| Gompertz | 0.041998 | 1.2118 | 0.12121 | — | -750.239 | 1506.48 |
| **Logistic** | **822.34** | **0.26988** | **20.150** | — | **-739.00** | **1484.00** |
| Power Law | 535.45 | 0.43942 | — | — | -774.08 | 1552.16 |
| Luecke Power Law | 611.83 | 0.31801 | -0.038286 | — | -753.05 | 1512.10 |

**Figure 7.**
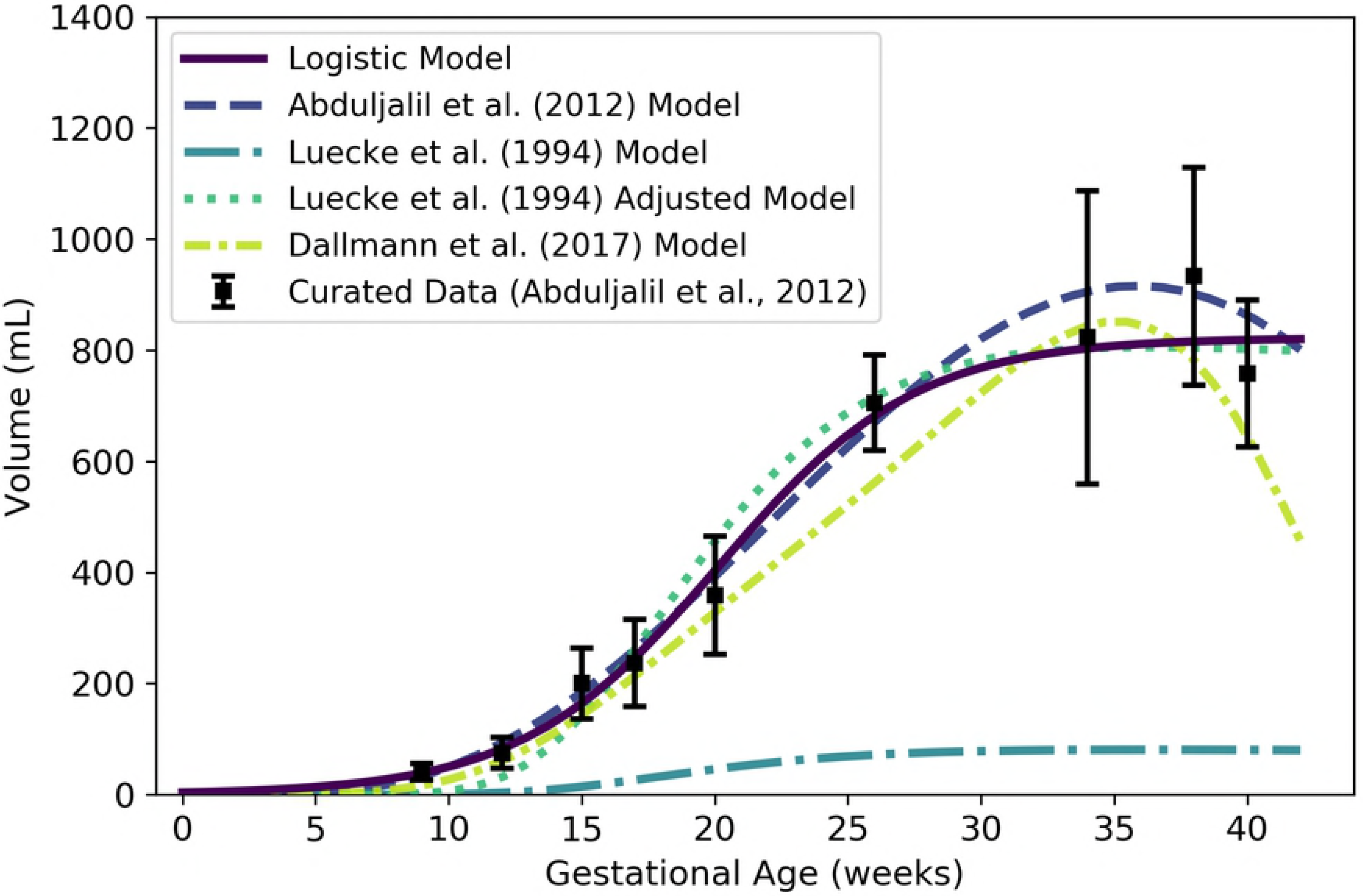
Amniotic fluid volume vs. gestational age. The logistic model (solid line) given by Equation 8 was selected as the most parsimonious model in our analysis. The model of Abduljalil et al. [28], which is a fifth degree polynomial model, was calibrated using the same curated data set [28] used by us. The model of Luecke et al. [15] was calibrated using different data and assumes a relationship between amniotic fluid volume and fetal mass. That model is shown here both as originally stated (in the publication) and after correcting a presumed error (to obtain the “Adjusted Model”) as described in the text. Dallmann et al. [3] calibrated their fourth degree polynomial model with different data.

Equation 8 yields nonnegative values for amniotic fluid volume on the time domain of interest (*t* > 0); however, as with the placenta volume model, it is worth noting that conception does not occur until a gestational age of about 2 weeks and so none of the products of conception (including amniotic fluid) exist until after that time. To be clear, Equation 8 should not be used to estimate the volume of the amniotic fluid prior to 2 weeks, and should not be considered accurate until 9 weeks (the time of the first data point of Abduljalil et al. [28]).

In Figure 7, we present two versions of the amniotic fluid volume model of Luecke et al. [15]. The first version shown (“Luecke et al. (1994) Model”) represents a Luecke power law model (cf. Table 2) with coefficient values as shown in Table 3 of Luecke et al. [15]. Since that model seems to under-predict amniotic fluid volumes by an order of magnitude, we hypothesized that the first coefficient printed in their table (θ_0_ = 0.002941) may have been off by a factor of 10. We therefore increased that coefficient by a factor of 10 (θ_0_ = 0.02941) to obtain the second model version (“Luecke et al. (1994) Adjusted Model”) shown in Figure 7.

#### Other Specific Compartments

Any given PBPK model may include compartments that correspond to specific organs and tissues in the mother that do not vary significantly during a normal pregnancy. To obtain volumes for such compartments that coincide with the time-varying volumes already described, we examined reference values for total body masses of “typical” women and the masses of various organs and tissues in such women as reported by the ICRP [33]. We used these to compute the percent of total body mass accounted for by various tissues. These percentages, along with densities associated with various organs and tissues, are displayed in Table 7. We calculated the static compartment volumes for a typical pregnant woman by multiplying the non-pregnant body mass (*W*^m^(0) = 61.103 kg) by the female mass percentage of the relevant tissue or organ and dividing by the relevant density. The volumes obtained for maternal brain, thyroid, kidneys, gut, liver, and lungs are shown in Table 12.

**Table 12.** Volumes of (some) maternal compartments that do not change during pregnancy.

| Compartment | Symbol | Value (L) |
| --- | --- | --- |
| Brain | $V_{\text{bran}}^{\text{m}}$ | 1.2749 |
| Thyroid | $V_{\text{thyr}}^{\text{m}}$ | 0.016469 |
| Kidneys | $V_{\text{kidn}}^{\text{m}}$ | 0.26653 |
| Gut | $V_{\text{gutx}}^{\text{m}}$ | 1.1110 |
| Liver | $V_{\text{livr}}^{\text{m}}$ | 1.3559 |
| Lungs | $V_{\text{lung}}^{\text{m}}$ | 0.91945 |

#### Rest of Body

We used the principle of mass balance to obtain a formula for the volume of a “rest of body” compartment comprising all mass in the pregnant female body that has not been accounted for in one of the specific compartments already described. Assuming that the fat-free mass of the mother has an average density of 1.1 g/mL throughout pregnancy [53], the total volume (L) of the maternal fat-free mass is

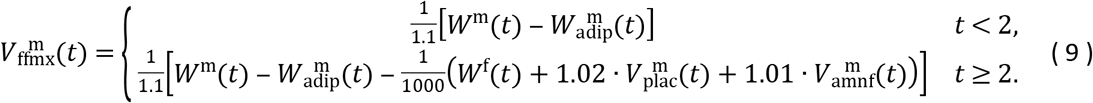

Here, we have assumed that the products of conception appear at gestational age 2 weeks, and that the densities of the placenta and amniotic fluid are 1.02 g/mL [51] and 1.01 g/mL [52], respectively (cf. Table 7). (We further assumed that *W*^m^, *W*^f^, 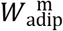, 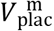, and 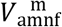 are described by Equations 1, 2, 3, 7, and 8, respectively.) Also, the total volume (L) of all the specific maternal compartments, excluding adipose tissue, is

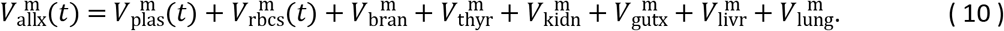

(We assumed that the quantities on the right-hand side of this equation are described by Equations 5 and 6b and the values listed in Table 12.) Thus, the volume of the maternal rest of body compartment (L) is

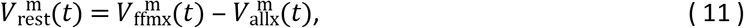

where *t* is the gestational age (weeks). As shown in Figure 8, Equation 11 results in volumes for the maternal rest of body compartment that fluctuate between approximately 31 and 34 L during pregnancy.

**Figure 8.**
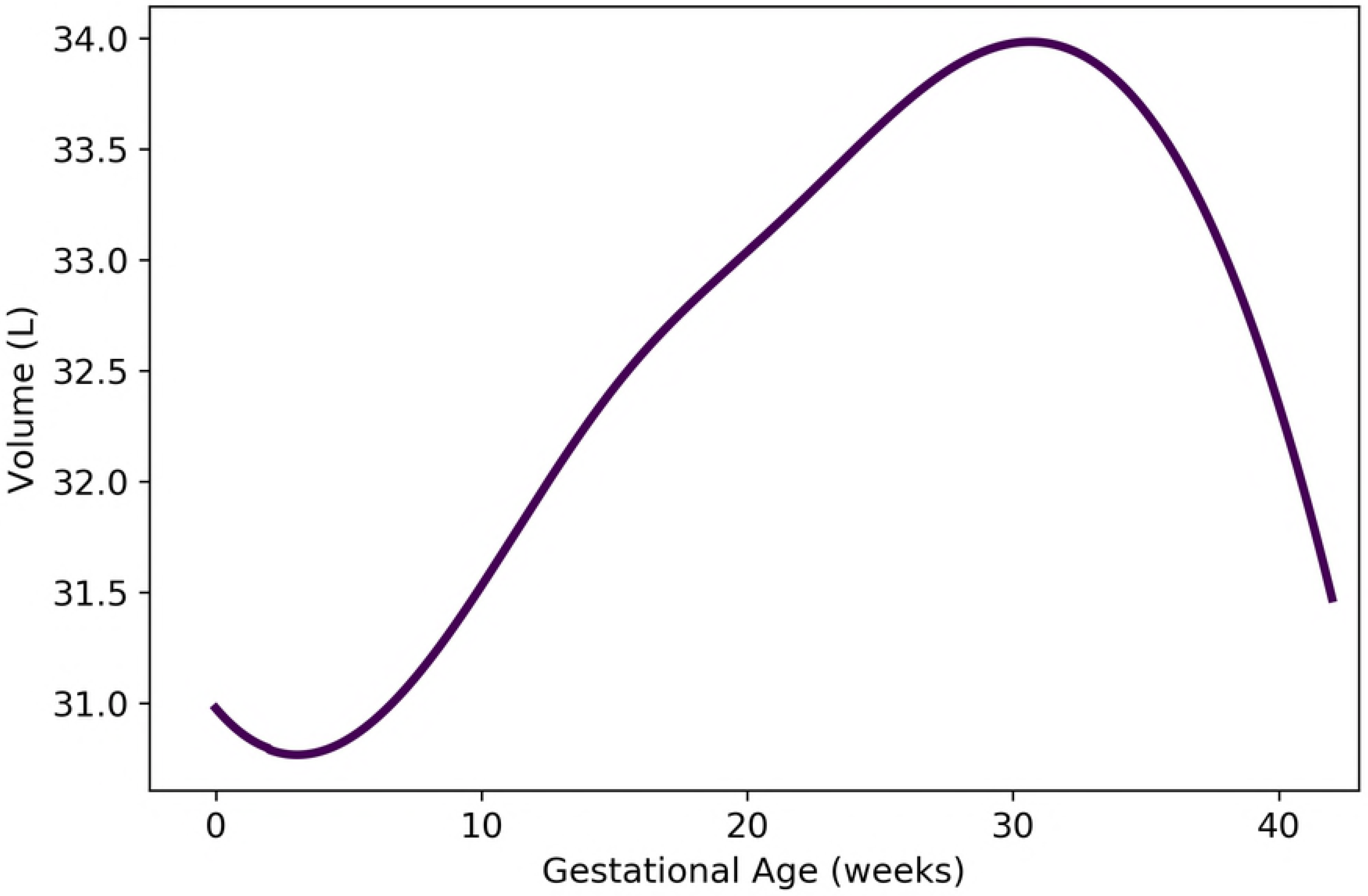
Volume of maternal “rest of body” compartment vs. gestational age. The formula for the depicted model, which is based on mass balance, is provided as Equation 11.

### Maternal Blood Flow Rates

During pregnancy, a woman experiences an increase in cardiac output as well as significant changes in the relative blood flow rates to various organs and tissues [33]. We first provide an expression for the maternal cardiac output, which corresponds to the quantity we denote 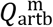 (the blood flow rate into the mother’s arterial blood compartment). Then, we describe flow rates to the other maternal compartments as proportions of the cardiac output. As shown in Table 2.44 from Section 12.2.7 of the ICRP [33] report, the percentage of cardiac output directed to any particular organ or tissue can change as pregnancy progresses. For blood flow rates for which time course data were not available, we assumed that the change from the “non-pregnant” percentage to the near-term “pregnant” percentage (as reported in that table) occurs in a linear fashion between 0 and 40 weeks of gestational age; i.e., the percentage is a linear function of the gestational age. Hereafter, we refer to such models as “linear transition models”.

#### Cardiac Output

We used the curated data of Abduljalil et al. [28] to calibrate various models for maternal cardiac output. We assume this to be both the total flow rate into the maternal arterial blood compartment (hence the use here of the symbol 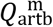) and the total flow rate into the maternal venous blood compartment. We selected the cubic model given by

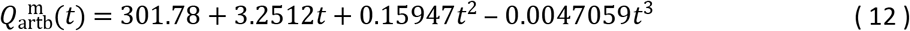

as the most parsimonious for maternal cardiac output (L/h) at gestational age *t* (weeks). Table 13 shows the maximum likelihood estimates of the parameter values for all models considered along with the associated log-likelihood and AIC values. Figure 9 shows the cubic model of Equation 12 along with several other models for maternal cardiac output.

**Table 13.** Maternal cardiac output models (flow rate in L/h vs. maternal mass in kg for power law models, flow rate in L/h vs. gestational age in weeks for all other models). For each model considered, the maximum likelihood parameter estimates 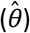, log-likelihood 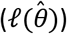, and AIC are provided. The row describing the selected model is shown in boldface.

| Model | $\hat{\theta}_0$ | $\hat{\theta}_1$ | $\hat{\theta}_2$ | $\hat{\theta}_3$ | $\ell(\hat{\theta})$ | AIC |
| --- | --- | --- | --- | --- | --- | --- |
| Linear | 309.15 | 3.1963 | — | — | -5195.06 | 10394.1 |
| Quadratic | 300.24 | 6.4348 | -0.094905 | — | -5178.48 | 10363.0 |
| **Cubic** | **301.78** | **3.2512** | **0.15947** | **-0.0047059** | **-5175.96** | **10359.9** |
| Modified Gompertz | 18.903 | 0.21568 | 0.11022 | 282.19 | -5179.72 | 10367.4 |
| Modified Logistic | 167.03 | 0.12506 | 6.0723 | 247.93 | -5179.64 | 10367.3 |
| Power Law | 0.81619 | 1.4473 | — | — | -5200.06 | 10404.1 |
| Luecke Power Law | 0.081119 | 2.5485 | -0.13127 | — | -5199.52 | 10405.0 |

**Figure 9.**
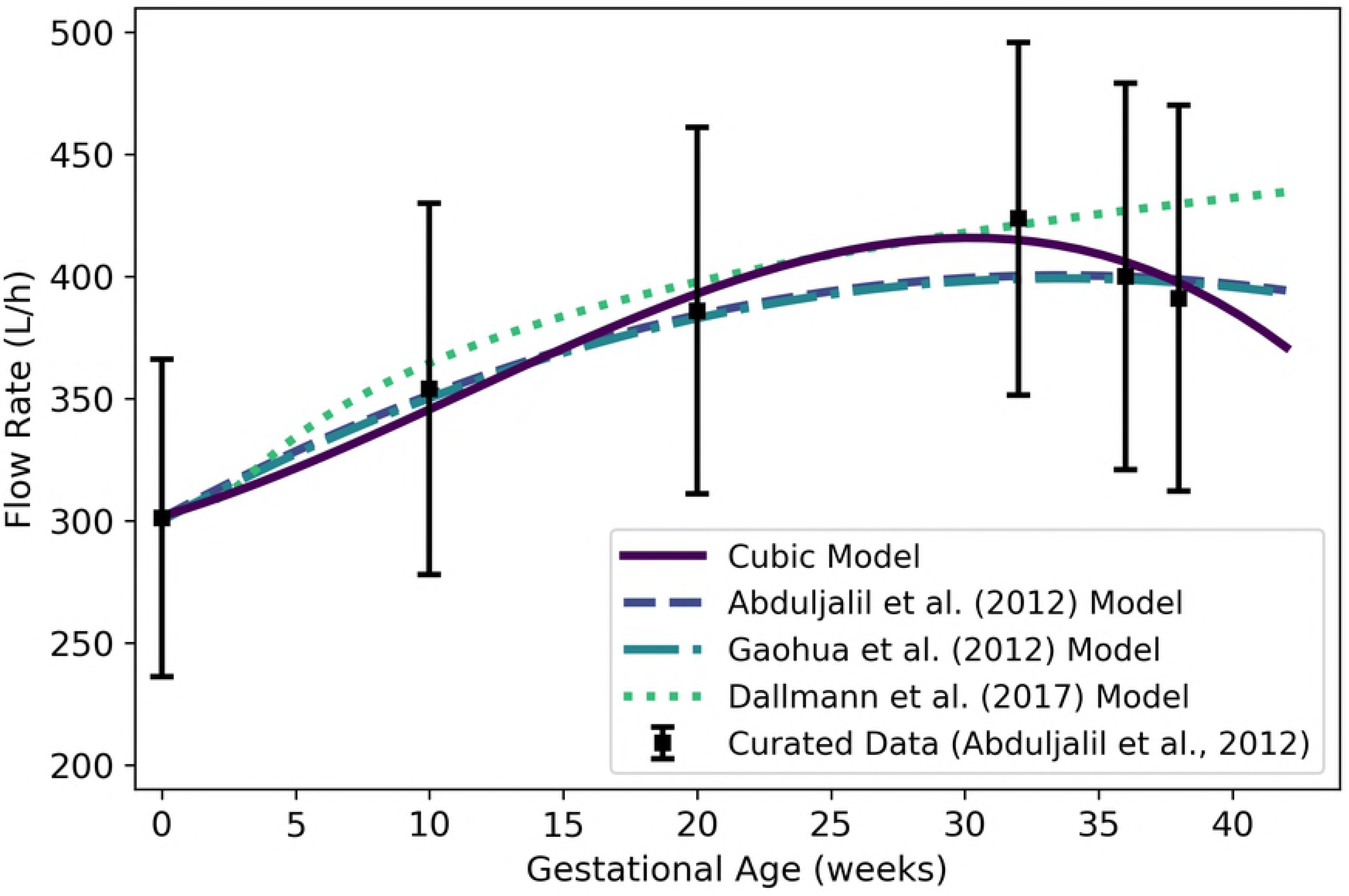
Maternal cardiac output vs. gestational age. The cubic model (solid line) given by Equation 12 was selected as the most parsimonious model in our analysis. The models of Abduljalil et al. [28] and Gaohua et al. [29], both of which are quadratic models, were calibrated using the same curated data set [28] used by us. Dallmann et al. [3] calibrated their model with different data.

#### Adipose Tissue

Assuming the blood flow rate into the maternal adipose tissue transitions linearly from 8.5% of cardiac output at 0 weeks to 7.8% near term [33], this flow rate (L/h) is given by

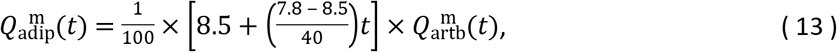

where *t* is the gestational age (weeks), where we have assumed “near term” corresponds to 40 weeks of gestational age.

#### Brain

Assuming the blood flow rate into the maternal brain transitions linearly from 12.0% of cardiac output at 0 weeks to 8.8% near term [33], this flow rate (L/h) is given by

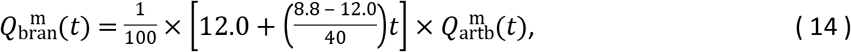

where *t* is the gestational age (weeks).

#### Kidneys

Assuming the blood flow rate into the kidneys transitions linearly from 17.0% of cardiac output at 0 weeks to 16.6% near term [33], this flow rate (L/h) is given by

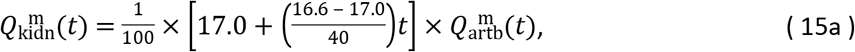

where *t* is the gestational age (weeks).

Since Abduljalil et al. [28] curated data for blood flow to the maternal kidneys during gestation, we also calibrated various models from Table 2 to describe kidney blood flow using this data set. We selected the cubic model given by

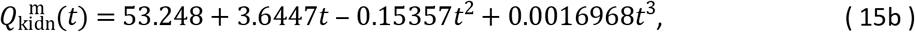

where 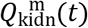 is the flow rate (L/h) at gestational age *t* (weeks), as the most parsimonious of the candidate models.

Table 14 shows the maximum likelihood estimates of the parameter values for all models considered along with the associated log-likelihood and AIC values. Both the cubic model of Equation 15b and the “linear transition” model of Equation 15a are depicted in Figure 10 along with two other models [3,28] for blood flow to the maternal kidneys.

**Table 14.** Maternal kidney blood flow models (flow rate in L/h vs. maternal mass in kg for power law models, flow rate in L/h vs. gestational age in weeks for all other models). For each model considered, the maximum likelihood parameter estimates 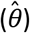, log-likelihood 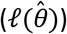, and AIC are provided. The row describing the selected model is shown in boldface.

| Model | $\hat{\theta}_0$ | $\hat{\theta}_1$ | $\hat{\theta}_2$ | $\hat{\theta}_3$ | $\ell(\hat{\theta})$ | AIC |
| --- | --- | --- | --- | --- | --- | --- |
| Linear | 59.979 | 0.41969 | — | — | -944.39 | 1892.78 |
| Quadratic | 54.042 | 2.6204 | -0.065712 | — | -894.31 | 1794.63 |
| **Cubic** | **53.248** | **3.6447** | **-0.15357** | **0.0016968** | **-891.88** | **1791.76** |
| Modified Gompertz | 9.6979 | 2.0563 | 1.8668 | 43.402 | -901.80 | 1811.59 |
| Modified Logistic | 38.466 | 1.6729 | 0.015416 | 34.115 | -901.80 | 1811.59 |
| Power Law | 1.8508 | 0.85156 | — | — | -950.55 | 1905.09 |
| Luecke Power Law | $4.9856 \times 10^{-5}$ | 5.8656 | -0.59716 | — | -948.88 | 1903.76 |

**Figure 10.**
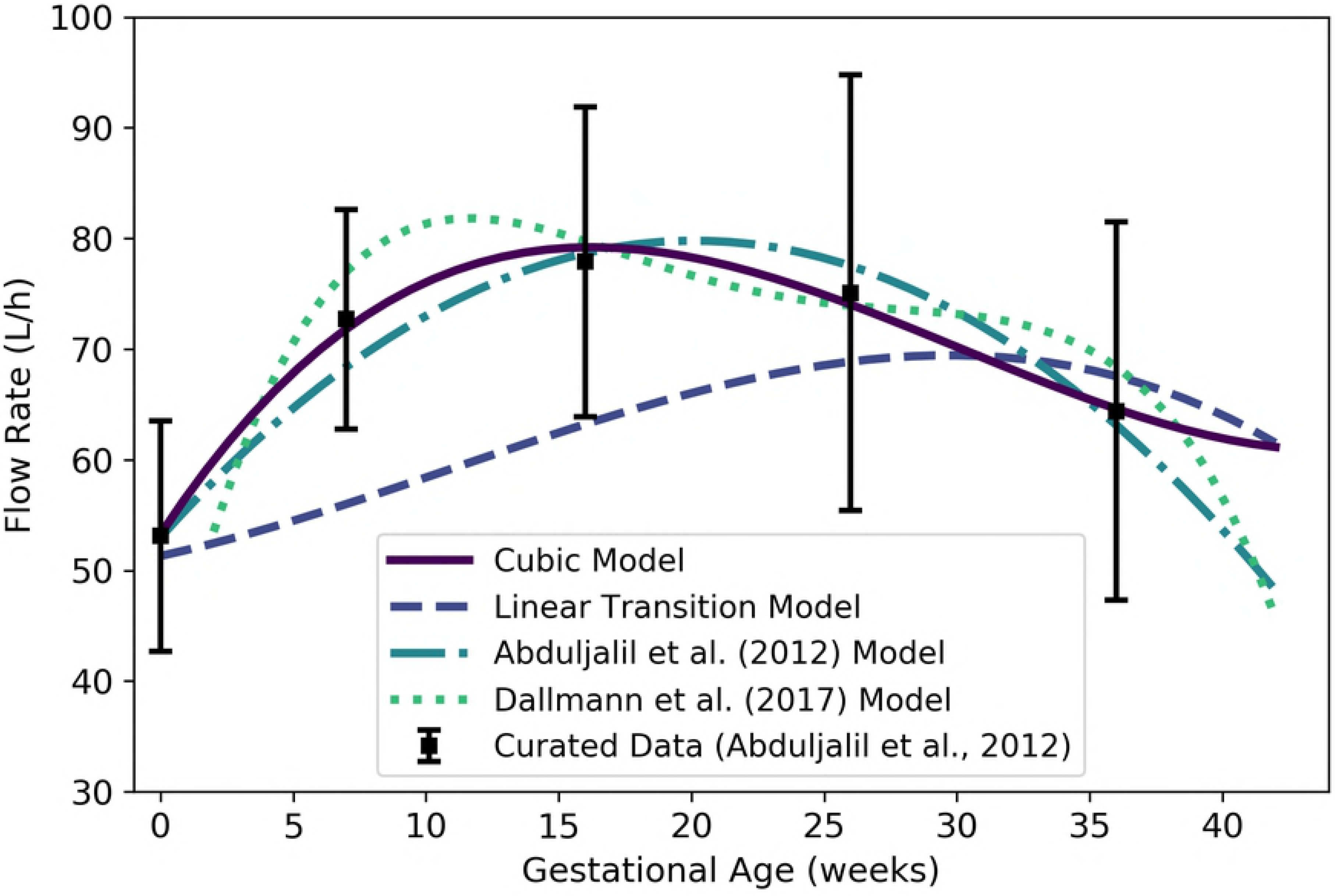
Maternal kidney blood flow vs. gestational age. The cubic model (solid line) given by Equation 15b was selected as the most parsimonious model in our analysis. The linear transition model given by Equation 15a is also shown. The model of Abduljalil et al. [28], which is a quadratic model, was calibrated using the same curated data set [28] used by us. Dallmann et al. [3] calibrated their model with different data.

#### Gut

Assuming the blood flow rate into the gut compartment transitions linearly from 17.0% of cardiac output at 0 weeks to 12.5% at 40 weeks [33], this flow rate (L/h) is given by

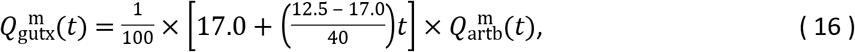

where *t* is the gestational age (weeks).

#### Liver

Assuming the blood flow rate into the liver transitions linearly from 27.0% of cardiac output at 0 weeks to 20.0% at 40 weeks [33], this flow rate (L/h) is given by

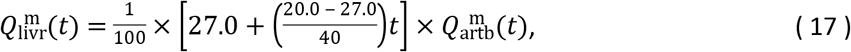

where *t* is the gestational age (weeks).

#### Thyroid

Assuming the blood flow rate into the thyroid transitions linearly from 1.5% of cardiac output at 0 weeks to 1.1% at 40 weeks [33], this flow rate (L/h) is given by

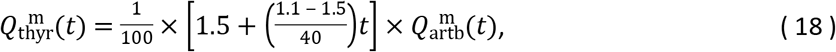

where *t* is the gestational age (weeks).

#### Placenta

Because the placenta is not listed in Table 2.44 of the ICRP [33] report, we constructed a model for maternal blood flow to the placenta by assuming that the percentage of maternal cardiac output reported for the uterus is actually a percentage for “uteroplacental” blood flow. According to Wang and Zhao [34], maternal blood flows to the placenta at about 600 to 700 mL/min (or 36 to 42 L/h) at term (end-of-pregnancy), accounting for 80% of the uteroplacental blood flow. Also, uteroplacental circulation is established at 11 to 12 days post-fertilization, corresponding to a gestational age of about 3.6 weeks [40]. If we assume that the blood flow rate to the uteroplacental region transitions linearly from 0.4% of cardiac output at 0 weeks to 12.0% at term [33], and further assume the proportion of this blood that flows directly to the placenta transitions from 0% at 3.6 weeks to 80% at 40 weeks, then the flow rate (L/h) to the placenta can be estimated using the linear transition model

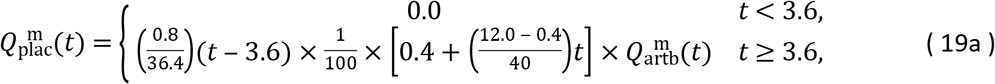

where *t* is the gestational age (weeks). This model is flawed in that it predicts unrealistically low values (considerably less than 36 L/h) close to term. Thus, as an alternative, we propose a model that sets maternal blood flow to the placenta proportional to the placenta volume with a proportionality constant (0.059176) selected to ensure that the maternal blood flow rate equal to the midpoint of the range given by Wang and Zhao [34] (39 L/h) at term (40 weeks). This model is given by

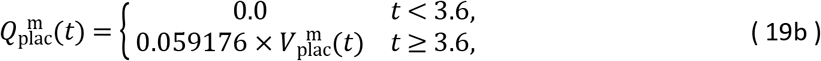

where 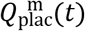 is the flow rate (L/h) to the placenta and 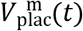 is the volume (mL) of the placenta (cf. Equation 7) at gestational age *t* (weeks). Figure 11 shows the linear transition model of Equation 19a, the proportional-to-volume model of Equation 19b, and two published models [3,29] for maternal blood flow to the placenta.

**Figure 11.**
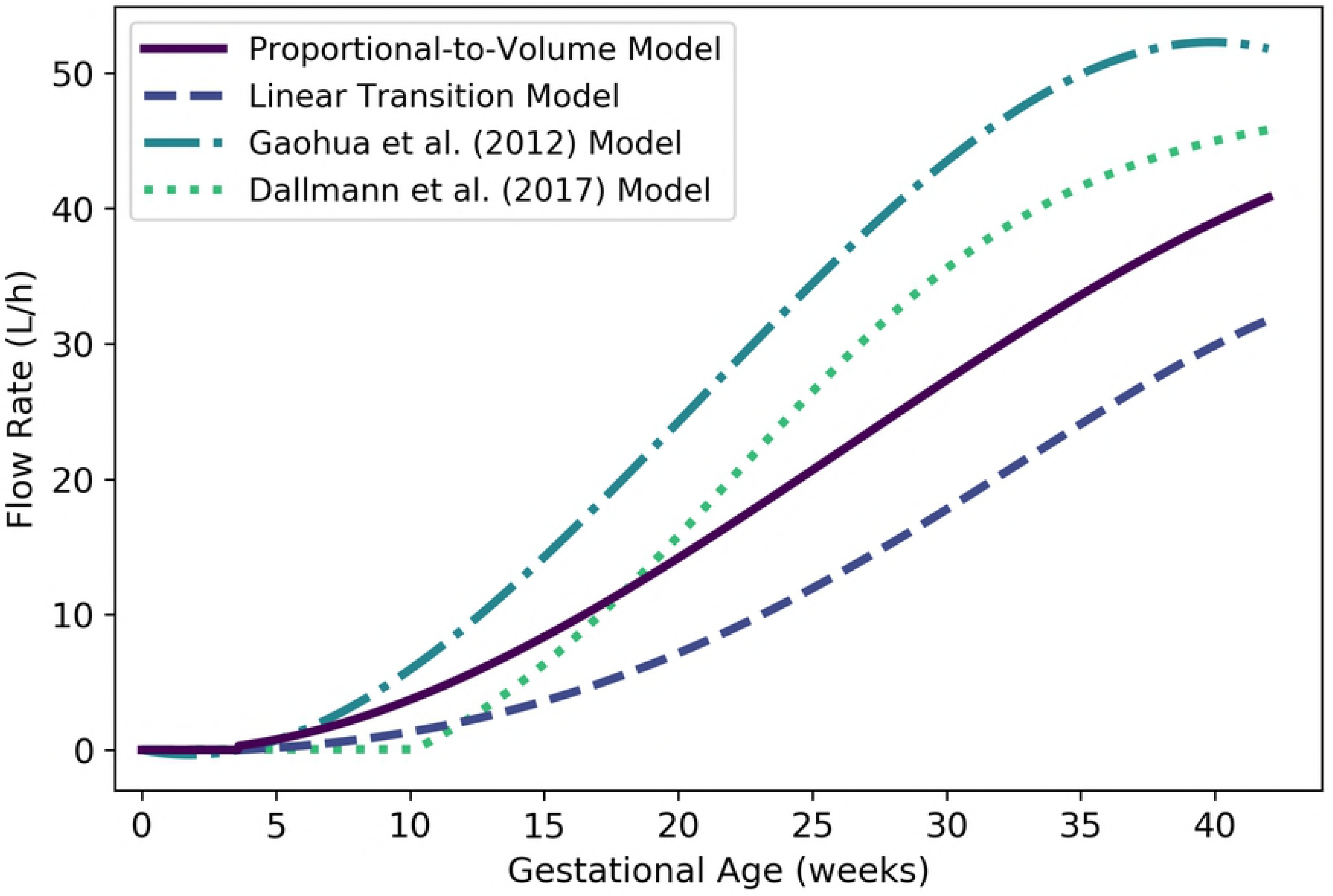
Maternal blood flow to the placenta vs. gestational age. The proportional-to-volume model (solid line) given by Equation 19b, the linear transition model given by Equation 19a, and two published models [3,29] are shown.

#### Rest of Body

We used the principle of conservation of flow to obtain a formula for the blood flow rate to the maternal rest of body compartment. Thus, the flow rate to the rest of body compartment (in L/h) is given by

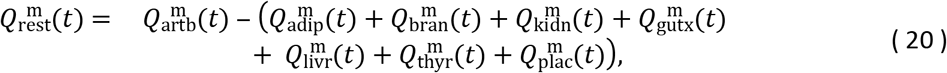

where *t* is the gestational age (in weeks) and 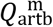, 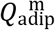, 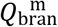, 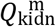, 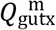, 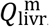, 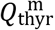 and 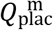 are given by Equations 12, 13, 14, 15b, 16, 17, 18, and 19b, respectively. As shown in Figure 12, Equation 20 results in flow rates to the maternal rest of body compartment that fluctuate between approximately 47 and and 97 L/h during pregnancy.

**Figure 12.**
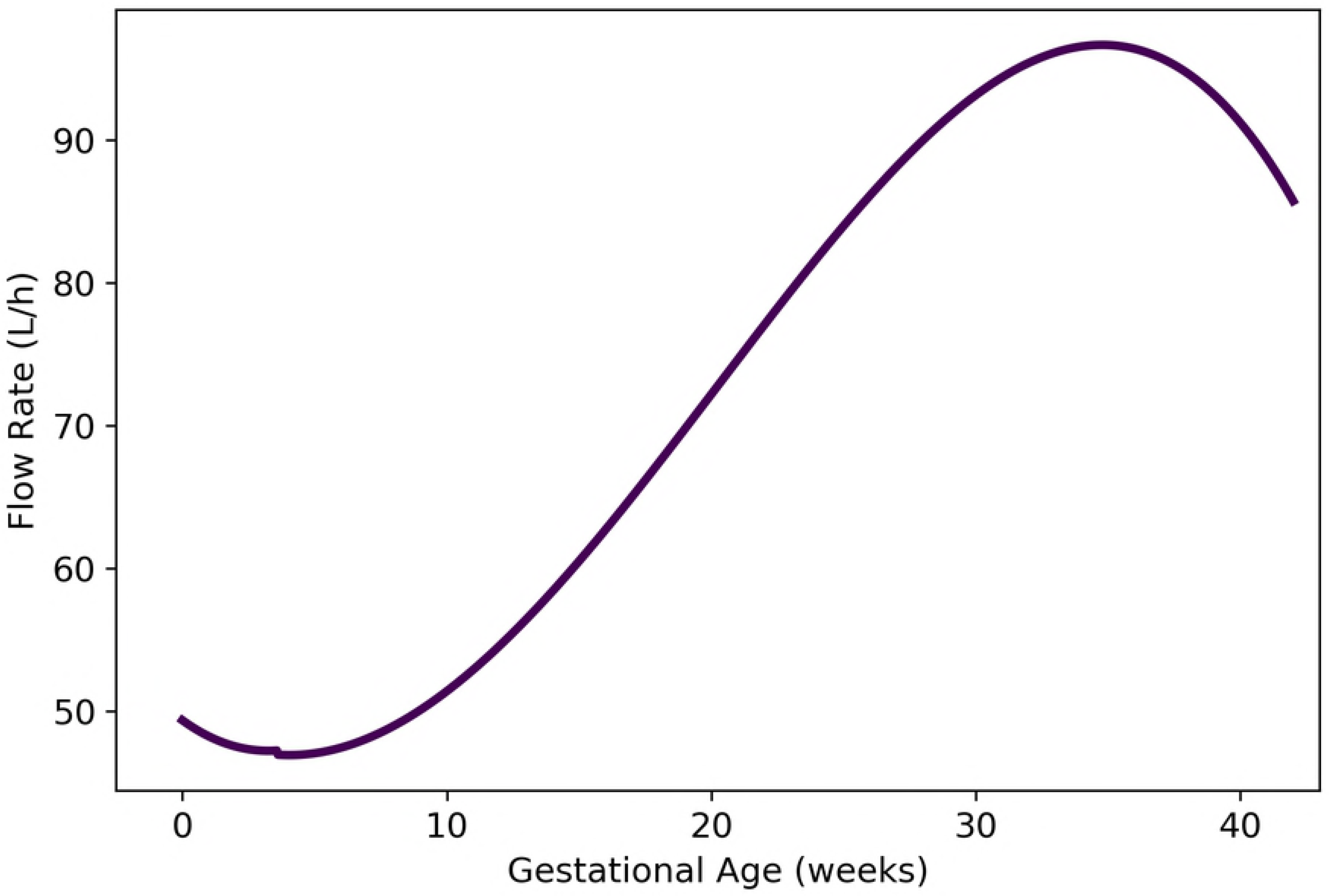
Maternal blood flow to the “rest of body” compartment vs. gestational age (cf. Equation 20).

### Other Maternal Physiological Parameters

#### Hematocrit

We used the curated data of Abduljalil et al. [28] to calibrate various models for maternal hematocrit. The quadratic model given by

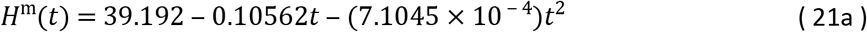

was selected as the most parsimonious model for hematocrit (as a percentage) at gestational age *t* (weeks). Table 15 shows the maximum likelihood estimates of the parameter values for all models considered along with the associated log-likelihood and AIC values. The quadratic model of Equation 21a and several other models [3,28,29] for maternal hematocrit are shown in Figure 13.

**Table 15.**
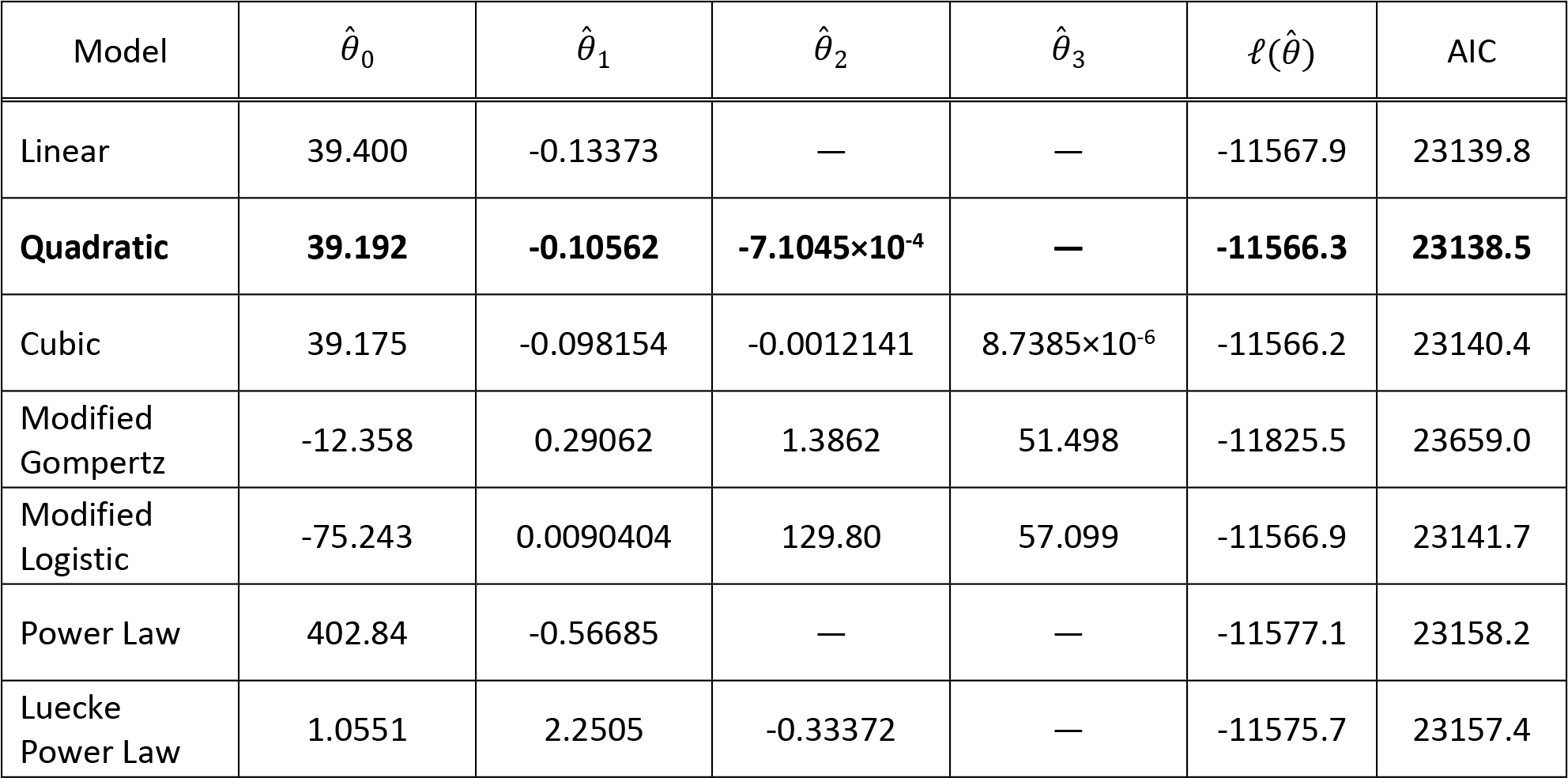
Maternal hematocrit models (percentage vs. maternal mass in kg for power law models, percentage vs. gestational age in weeks for all other models). For each model considered, the maximum likelihood parameter estimates 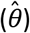, log-likelihood 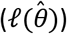, and AIC are provided. The row describing the selected model is shown in boldface.

| Model | $\hat{\theta}_0$ | $\hat{\theta}_1$ | $\hat{\theta}_2$ | $\hat{\theta}_3$ | $\ell(\hat{\theta})$ | AIC |
| --- | --- | --- | --- | --- | --- | --- |
| Linear | 39.400 | -0.13373 | — | — | -11567.9 | 23139.8 |
| **Quadratic** | **39.192** | **-0.10562** | **-7.1045×10<sup>-4</sup>** | — | **-11566.3** | **23138.5** |
| Cubic | 39.175 | -0.098154 | -0.0012141 | 8.7385×10<sup>-6</sup> | -11566.2 | 23140.4 |
| Modified Gompertz | -12.358 | 0.29062 | 1.3862 | 51.498 | -11825.5 | 23659.0 |
| Modified Logistic | -75.243 | 0.0090404 | 129.80 | 57.099 | -11566.9 | 23141.7 |
| Power Law | 402.84 | -0.56685 | — | — | -11577.1 | 23158.2 |
| Luecke Power Law | 1.0551 | 2.2505 | -0.33372 | — | -11575.7 | 23157.4 |

**Figure 13.**
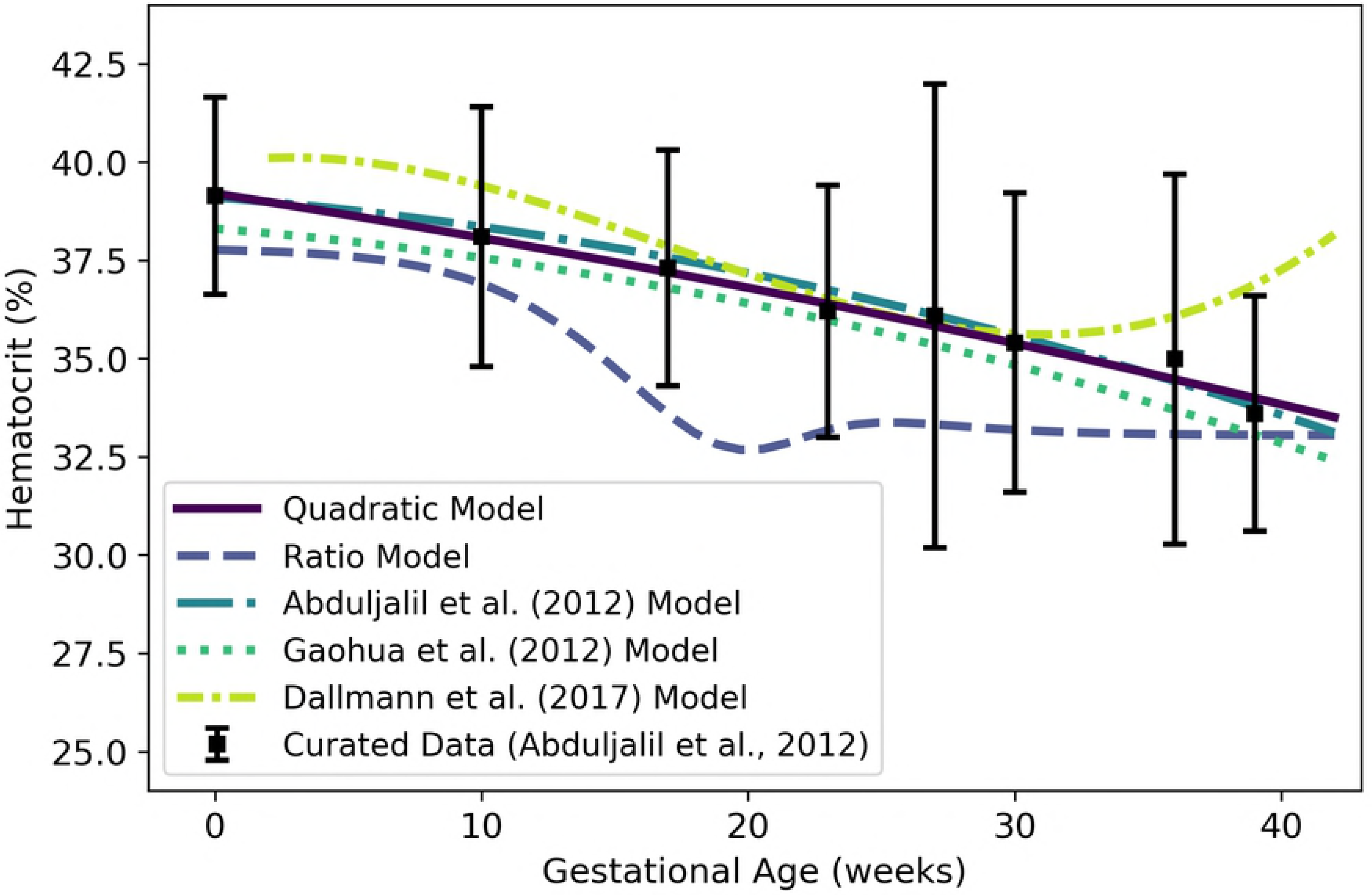
Maternal hematocrit vs. gestational age. The quadratic model (solid line) given by Equation 21a was selected as the most parsimonious model in our analysis. The “ratio model” given by Equation 21b is also shown. The models of Abduljalil et al. [28] and Gaohua et al. [29], both of which are quadratic models, were calibrated using the same curated data set [28] used by us. Dallmann et al. [3] calibrated their model with different data.

Because hematocrit represents the volume percentage of red blood cells in whole blood, and because whole blood is mostly made up of plasma and red blood cells (with only a small fraction made up of white blood cells and platelets), we can estimate the hematocrit using our models for maternal plasma volume, 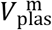, and maternal red blood cell volume, 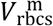. That is, another model for hematocrit is given by

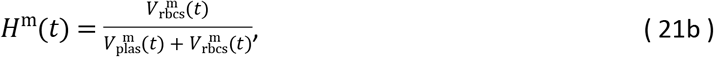

where *t* is the gestational age (weeks) and 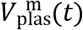 and 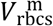 represent the maternal plasma and red blood cells volumes given by Equations 5 and 6, respectively, at gestational week *t*. We refer the hematocrit model of Equation 21b as the “ratio model” in Figure 13. There, we see that the ratio model yields a lower hematocrit than any of the other models (including the quadratic model of Equation 21a). Note that this ratio model considers the volume of blood components other than plasma and RBCS to be negligible; if such components have a non-negligible volume, the hematocrit values predicted by this model should *over*estimate hematocrit.

#### Glomerular Filtration Rate

We used the curated data of Abduljalil et al. [28] to calibrate various models for glomerular filtration rate (GFR). The quadratic model given by

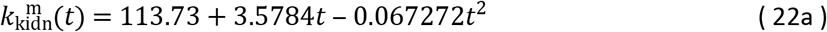

was selected as the most parsimonious model for GFR (mL/min) at gestational age *t* (weeks). Table 16 shows the maximum likelihood estimates of the parameter values for all models considered along with the associated log-likelihood and AIC values. The linear model of Equation 22a and two other models for maternal GFR [3,28] are shown in Figure 14.

**Table 16.** Maternal GFR models (rate in mL/min vs. maternal mass in kg for power law models, rate in mL/min vs. gestational age in weeks for all other models). For each model considered, the maximum likelihood parameter estimates 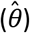, log-likelihood 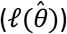, and AIC are provided. The row describing the selected model is shown in boldface.

| Model | $\hat{\theta}_0$ | $\hat{\theta}_1$ | $\hat{\theta}_2$ | $\hat{\theta}_3$ | $\ell(\hat{\theta})$ | AIC |
| --- | --- | --- | --- | --- | --- | --- |
| Linear | 122.82 | 1.3341 | — | — | -1379.34 | 2762.68 |
| **Quadratic** | **113.73** | **3.5784** | **-0.067272** | — | **-1363.91** | **2733.81** |
| Cubic | 113.66 | 3.6732 | -0.074784 | $1.4036 \times 10^{-4}$ | -1363.90 | 2735.80 |
| Modified Gompertz | $2.6041 \times 10^{-5}$ | 4.4691 | 0.31100 | 113.87 | -1363.38 | 2734.77 |
| Modified Logistic | 45.062 | 0.46667 | 10.026 | 113.59 | -1363.25 | 2734.50 |
| Power Law | 0.41867 | 1.3891 | — | — | -1384.28 | 2772.56 |
| Luecke Power Law | $8.9811 \times 10^{-5}$ | 5.4117 | -0.47878 | — | -1383.17 | 2772.34 |

**Figure 14.**
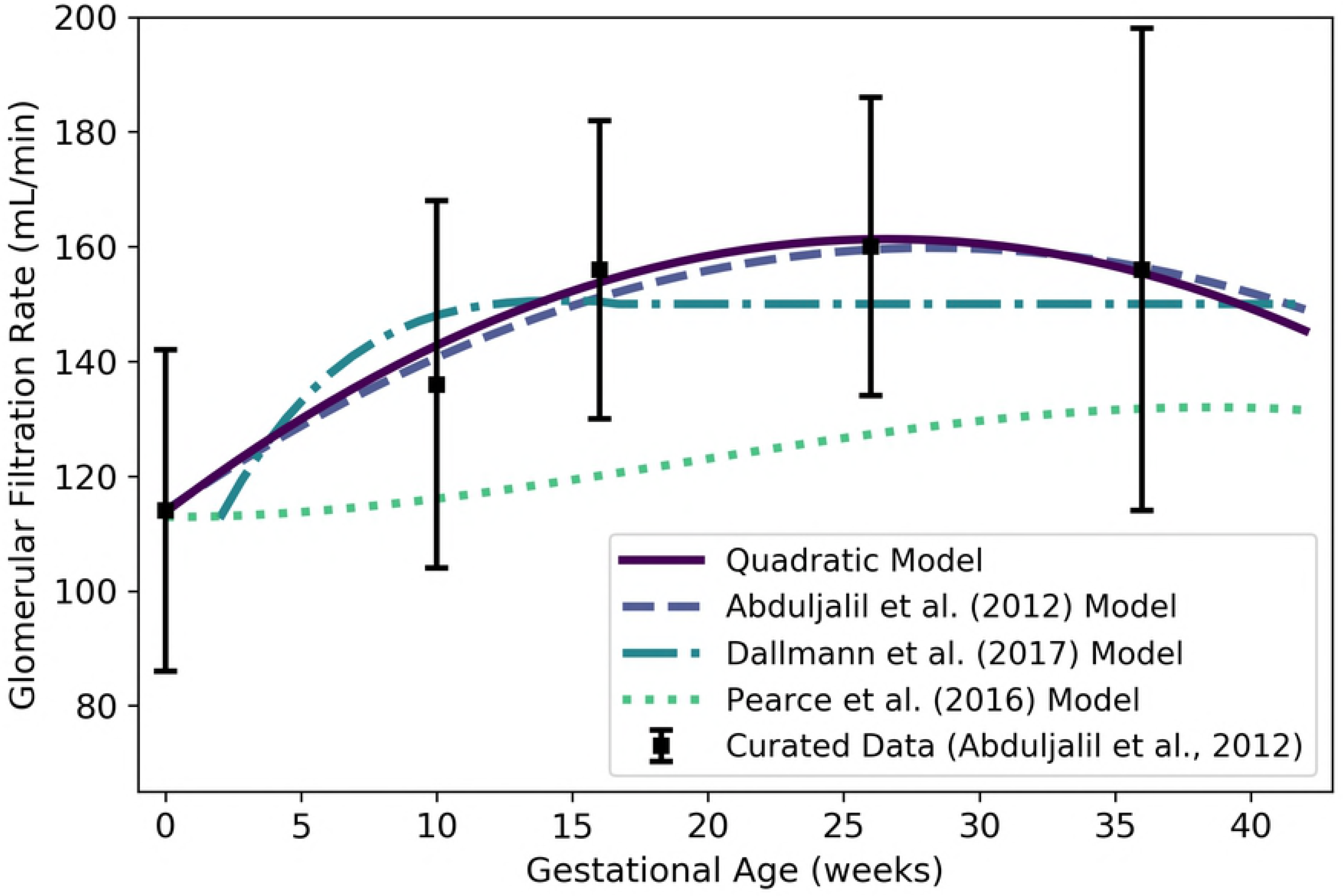
Maternal glomerular filtration rate vs. gestational age. The quadratic model (solid line) given by Equation 22 was selected as the most parsimonious model in our analysis. The model of Abduljalil et al. [28], also a quadratic model, was calibrated using the same curated data set [28] used by us. The Dallmann et al. [3] model depicted here has been modified from the published version, which contained typographical errors, based on personal correspondence with the lead author. The model attributed to Pearce et al. [54] is evaluated as described in the text.

Pearce et al. [54] have used an allometric model for GFR. These authors assume that for a 70 kg human, GFR is 125 mL/min [55] and glomerular filtration can be computed as a multiple of body mass to the ¾ power [56]. Thus, one can compute the glomerular filtration as

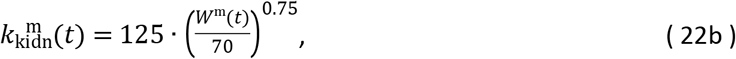

where *t* is the gestational age (in weeks) and *W*^m^(t) represents the maternal mass given by Equation 1 at gestational week *t*. For purposes of comparison, we show the plot of Equation 22b in Figure 14. It is labeled there as the “Pearce et al. (2016) Model”.

### Fetal Compartment Volumes

#### Brain

We used data curated by Abduljalil et al. [31] to calibrate various models for the brain mass of a human fetus during gestation. The cubic growth model given by

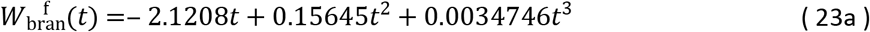

was selected as the most parsimonious model for fetal brain *mass* (g) at gestational age *t* (weeks); however, this model gives negative values for fetal brain mass during early gestation. The Gompertz model given by

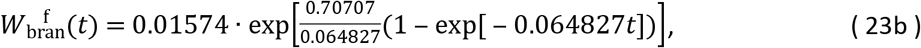

has the next lowest AIC (21893.5 vs. 21861.4), but it yields strictly positive values for gestational ages greater than or equal to zero. Table 17 shows the maximum likelihood estimates of the parameter values for all models considered along with the associated log-likelihood and AIC values. The cubic growth model of Equation 23a, the Gompertz model of Equation 23b, two published models [15,31], and the curated summary data [31] are shown in Figure 15.

**Table 17.** Fetal brain mass models (g vs. fetal mass in g for power law models, g vs. gestational age in weeks for all other models). For each model considered, the maximum likelihood parameter estimates 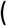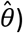, log-likelihood 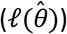, and AIC are provided. The row describing the selected model is shown in boldface.

| Model | $\hat{\theta}_0$ | $\hat{\theta}_1$ | $\hat{\theta}_2$ | $\hat{\theta}_3$ | $\ell(\hat{\theta})$ | AIC |
| --- | --- | --- | --- | --- | --- | --- |
| Linear Growth | 2.9256 | — | — | — | -21528.2 | 43058.5 |
| Quadratic Growth | -3.9135 | 0.32870 | — | — | -11008.6 | 22021.2 |
| **Cubic Growth** | **-2.1208** | **0.15645** | **0.0034746** | — | **-10927.7** | **21861.4** |
| Gompertz | 0.015740 | 0.70707 | 0.064827 | — | -10943.7 | 21893.5 |
| Logistic | 462.90 | 0.19240 | 32.321 | — | -11074.2 | 22154.4 |
| Power Law | 0.35659 | 0.85318 | — | — | -10970.6 | 21945.2 |
| Luecke Power Law | 0.93210 | 0.55821 | 0.021936 | — | -10956.8 | 21919.6 |

**Figure 15.**
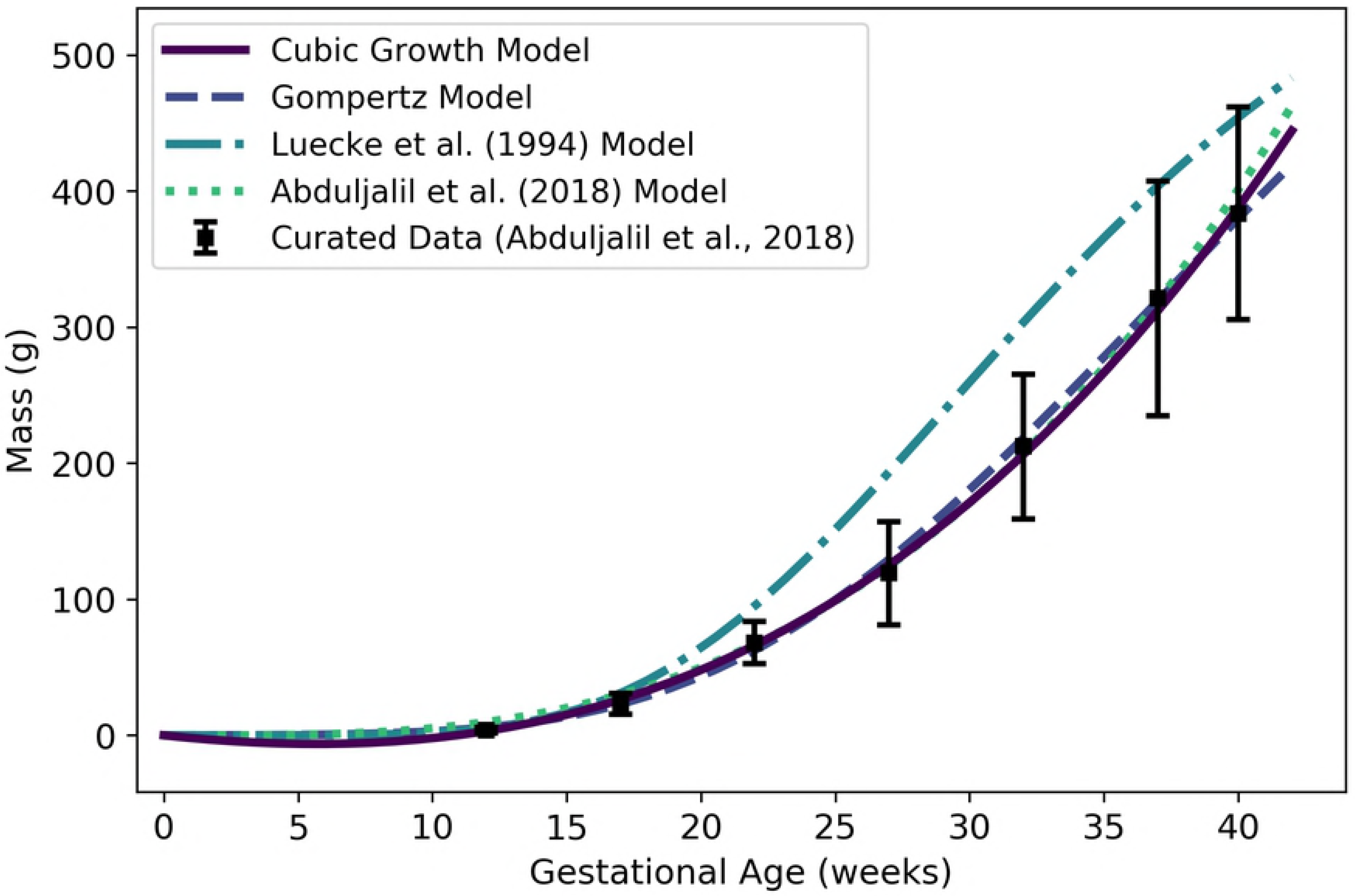
Fetal brain mass vs. gestational age. The quadratic growth model (solid line) given by Equation 23a was selected as the most parsimonious model in our analysis; however, that model gives negative brain mass values during early gestation. The Gompertz model (dashed line) given by Equation 23b is strictly positive on the time domain of interest. The model of Luecke et al. [15], which is a power law model based on fetal mass, was calibrated using a different data set [25].

Since the mean density of human brain tissue is 1.04 g/mL [33], the *volume* (mL) of the fetal brain tissue can be computed as

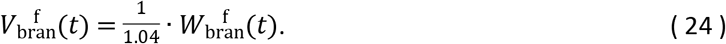

#### Liver

We used data curated by Abduljalil et al. [31] to calibrate various models for the liver mass of a human fetus during gestation. The cubic growth model given by

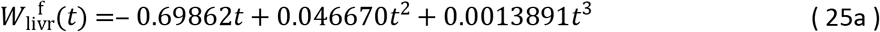

was selected as the most parsimonious model for fetal liver *mass* (g) at gestational week age *t* (weeks); however, this model gives negative values for fetal liver mass during early gestation. The Gompertz model given by

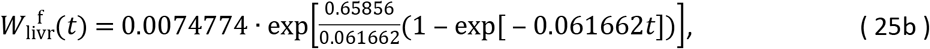

yields strictly positive values for gestational ages greater than or equal to zero and has the next lowest AIC (16430.6 vs. 16402.1). Table 18 shows the maximum likelihood estimates of the parameter values for all models considered along with the associated log-likelihood and AIC values. The cubic growth model of Equation 25a, the Gompertz model of Equation 25b, two published models [15,31], and the curated summary data [31] are shown in Figure 16.

**Table 18.** Fetal liver mass models (g vs. fetal mass in g for power law models, g vs. gestational age in weeks for all other models). For each model considered, the maximum likelihood parameter estimates 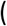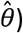, log-likelihood 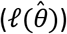, and AIC are provided. The row describing the selected model is shown in boldface.

| Model | $\hat{\theta}_0$ | $\hat{\theta}_1$ | $\hat{\theta}_2$ | $\hat{\theta}_3$ | $\ell(\hat{\theta})$ | AIC |
| --- | --- | --- | --- | --- | --- | --- |
| Linear Growth | 1.0599 | — | — | — | -12411.80 | 24825.6 |
| Quadratic Growth | -1.4130 | 0.11397 | — | — | -8241.18 | 16486.4 |
| **Cubic Growth** | **-0.69862** | **0.046670** | **0.0013891** | — | **-8198.03** | **16402.1** |
| Gompertz | 0.0074774 | 0.65856 | 0.061662 | — | -8212.32 | 16430.6 |
| Logistic | 161.85 | 0.18771 | 32.712 | — | -8275.32 | 16556.6 |
| Power Law | 0.10885 | 0.86562 | — | — | -8231.05 | 16466.1 |
| Luecke Power Law | 0.38023 | 0.48526 | 0.028240 | — | -8221.42 | 16448.8 |

**Figure 16.**
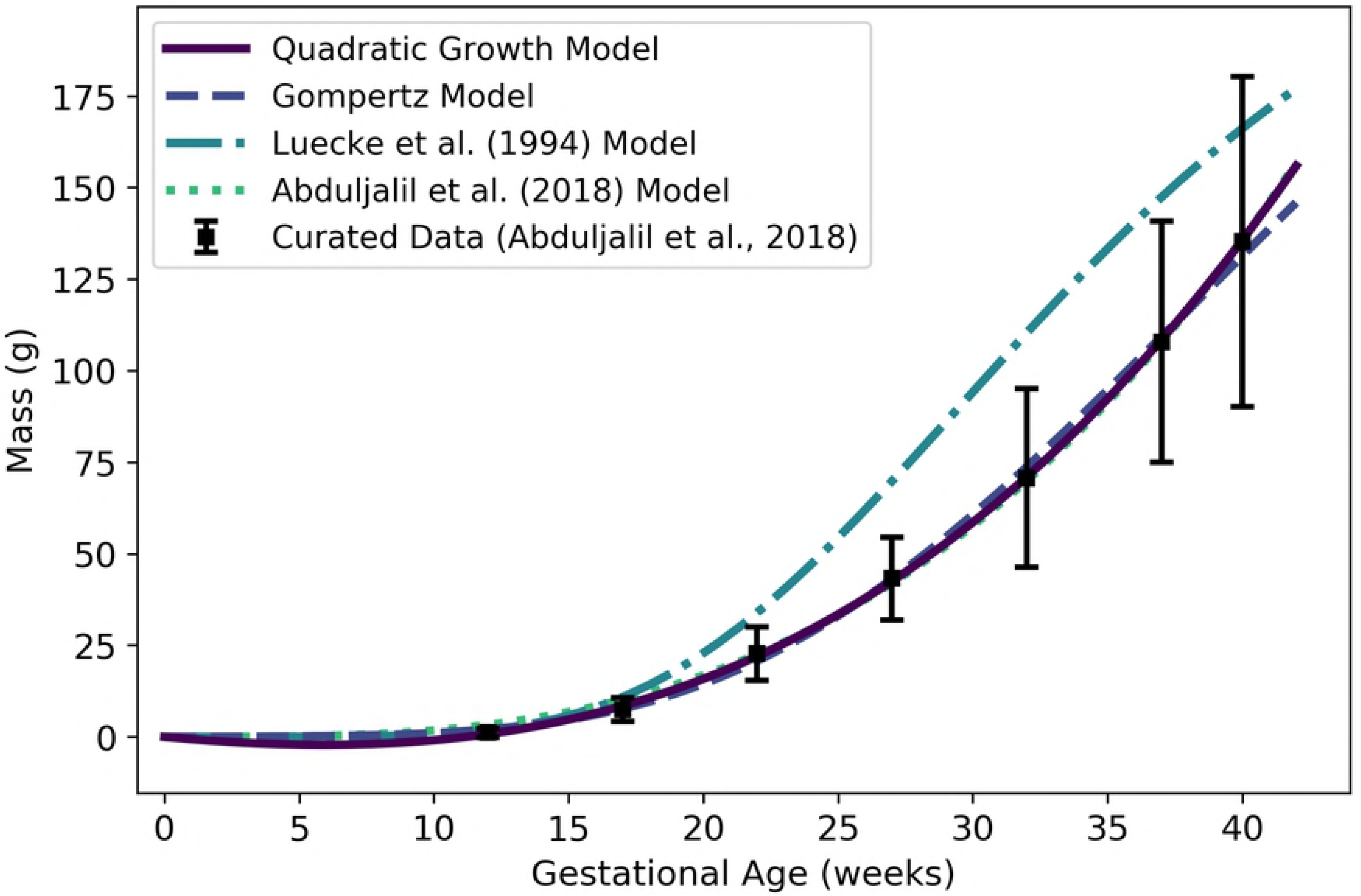
Fetal liver mass vs. gestational age. The quadratic growth model (solid line) given by Equation 25a was selected as the most parsimonious model in our analysis; however, that model gives negative liver mass values during early gestation. The Gompertz model (dashed line) given by Equation 25b is strictly positive on the time domain of interest. The model of Luecke et al. [15], which is a power law model based on fetal mass, was calibrated using a different data set [25].

Since the mean density of human liver tissue is 1.05 g/mL [35], the *volume* (mL) of the fetal liver tissue can be computed as

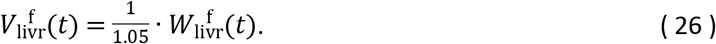

#### Kidneys

We used data curated by Abduljalil et al. [31] to calibrate various models for the kidney mass of a human fetus during gestation. The power law model given by

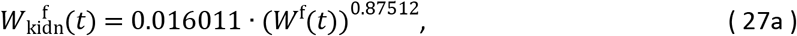

where *W*^f^(*t*) denotes the fetal mass (g) given by Equation 2, was selected as the most parsimonious model for fetal kidney *mass* (g) at gestational age *t* (weeks). The power law relates fetal kidney mass to total fetal mass and thus requires an (intermediate) estimate of total fetal mass at each time point of interest. The Gompertz model given by

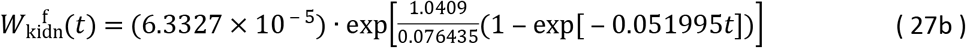

has an AIC that is only slightly larger (11838.3 vs. 11836.6) and does not require an intermediate calculation for total fetal mass. Table 19 shows the maximum likelihood estimates of the parameter values for all models considered along with the associated log-likelihood and AIC values. The power law model of Equation 27a, the Gompertz model of Equation 27b, two published models [15,31], and the curated summary data [31] are shown in Figure 17.

**Table 19.** Fetal kidney mass models (g vs. fetal mass in g for power law models, g vs. gestational age in weeks for all other models). For each model considered, the maximum likelihood parameter estimates 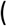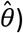, log-likelihood 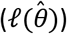, and AIC are provided. The row describing the selected model is shown in boldface.

| Model | $\hat{\theta}_0$ | $\hat{\theta}_1$ | $\hat{\theta}_2$ | $\hat{\theta}_3$ | $\ell(\hat{\theta})$ | AIC |
| --- | --- | --- | --- | --- | --- | --- |
| Linear Growth | 0.10014 | — | — | — | -12495.00 | 24992.0 |
| Quadratic Growth | -0.26646 | 0.022757 | — | — | -6097.13 | 12198.3 |
| Cubic Growth | -0.093661 | 0.0043005 | 0.00040110 | — | -5967.55 | 11941.1 |
| Gompertz | $6.33270 \times 10^{-5}$ | 1.0409 | 0.076435 | — | -5916.16 | 11838.3 |
| Logistic | 29.583 | 0.23193 | 30.697 | — | -5984.42 | 11974.8 |
| **Power Law** | **0.016011** | **0.91410** | — | — | **-5916.29** | **11836.6** |
| Luecke Power Law | 0.018129 | 0.87512 | 0.0029479 | — | -5916.15 | 11838.3 |

**Figure 17.**
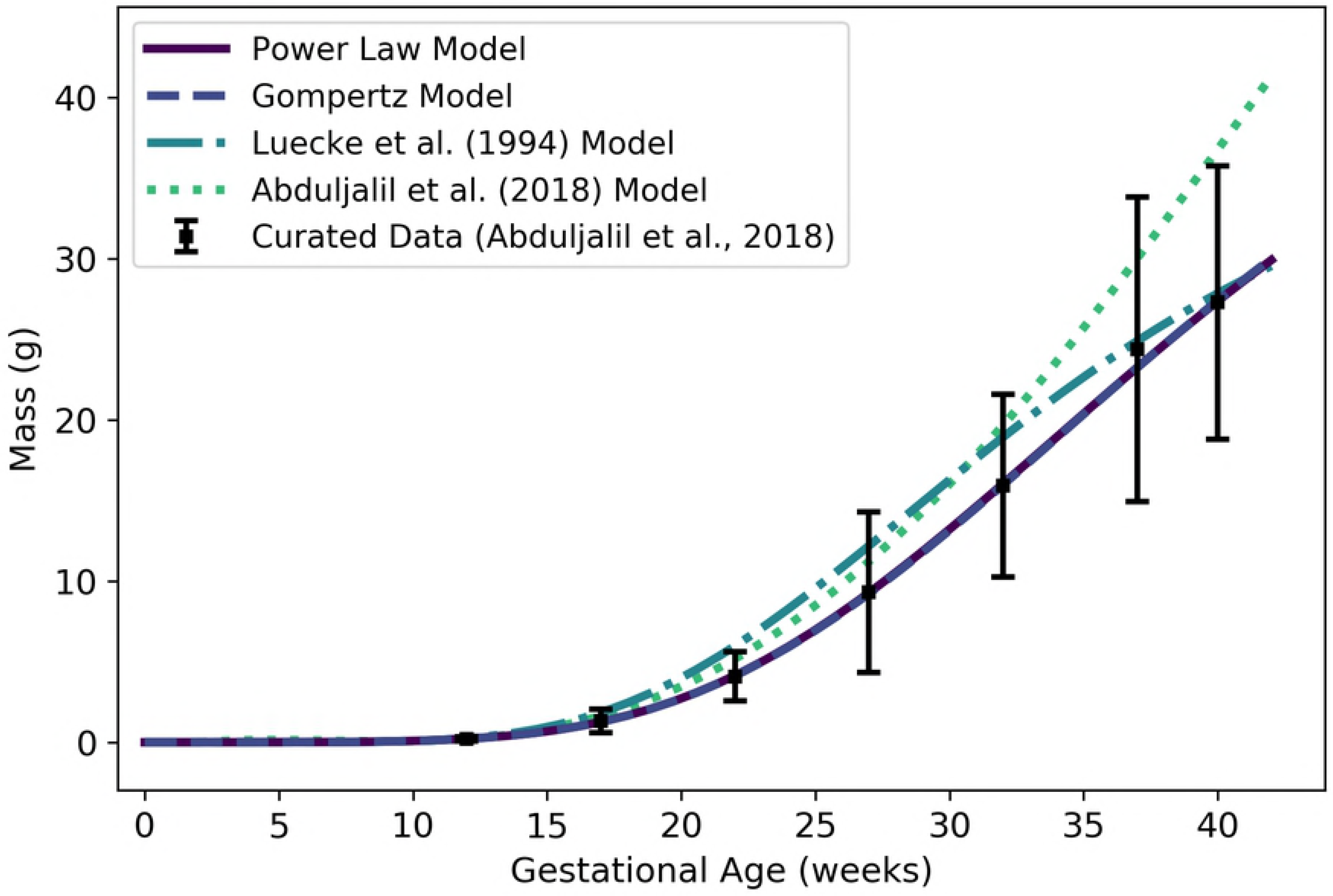
Fetal kidney mass vs. gestational age. The power law model (solid line) given by Equation 27a was selected as the most parsimonious model in our analysis; however, that model is a function of fetal mass. The Gompertz model (dashed line) given by Equation 27b may be preferred since it is a function of gestational age and does not require an intermediate model for fetal mass. Note that the Gompertz model and the power law model are virtually indistinguishable in this plot. The model of Luecke et al. [15], which is also a power law model based on fetal mass, was calibrated using a different data set [25]. The apparently poor fit of the model of Abduljalil et al. [31] to their own curated data set is probably due to precision-related errors – they only reported the leading coefficient of their polynomial model to one significant figure.

Since the mean density of human kidney tissue is 1.05 g/mL [33], the *volume* (mL) of the fetal kidneys can be computed as

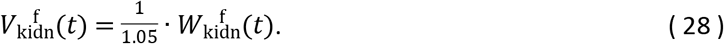

#### Lungs

We used data curated by Abduljalil et al. [31] to calibrate various models for the lung mass of a human fetus during gestation. The Gompertz model given by

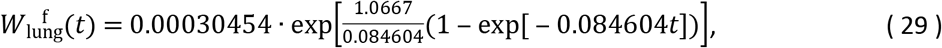

was selected as the most parsimonious model for fetal lung *mass* (g) at gestational age *t* (weeks). Table 20 shows the maximum likelihood estimates of the parameter values for all models considered along with the associated log-likelihood and AIC values. The Gompertz model of Equation 29, two published models [15,31], and the curated summary data [31] are shown in Figure 18.

**Table 20.** Fetal lung mass models (g vs. fetal mass in g for power law models, g vs. gestational age in weeks for all other models). For each model considered, the maximum likelihood parameter estimates 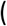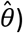, log-likelihood 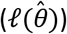, and AIC are provided. The row describing the selected model is shown in boldface.

| Model | $\hat{\theta}_0$ | $\hat{\theta}_1$ | $\hat{\theta}_2$ | $\hat{\theta}_3$ | $\ell(\hat{\theta})$ | AIC |
| --- | --- | --- | --- | --- | --- | --- |
| Linear Growth | 0.49354 | — | — | — | -13910.20 | 27822.4 |
| Quadratic Growth | -0.58156 | 0.053565 | — | — | -8893.10 | 17790.2 |
| Cubic Growth | -0.57275 | 0.052706 | $1.8119 \times 10^{-5}$ | — | -8893.05 | 17792.1 |
| **Gompertz** | **0.00030454** | **1.0667** | **0.084604** | — | **-8823.24** | **17652.5** |
| Logistic | 59.996 | 0.22339 | 28.200 | — | -8866.84 | 17739.7 |
| Power Law | 0.12888 | 0.75526 | — | — | -8827.70 | 17659.4 |
| Luecke Power Law | 0.074365 | 0.92497 | -0.012730 | — | -8824.40 | 17654.8 |

**Figure 18.**
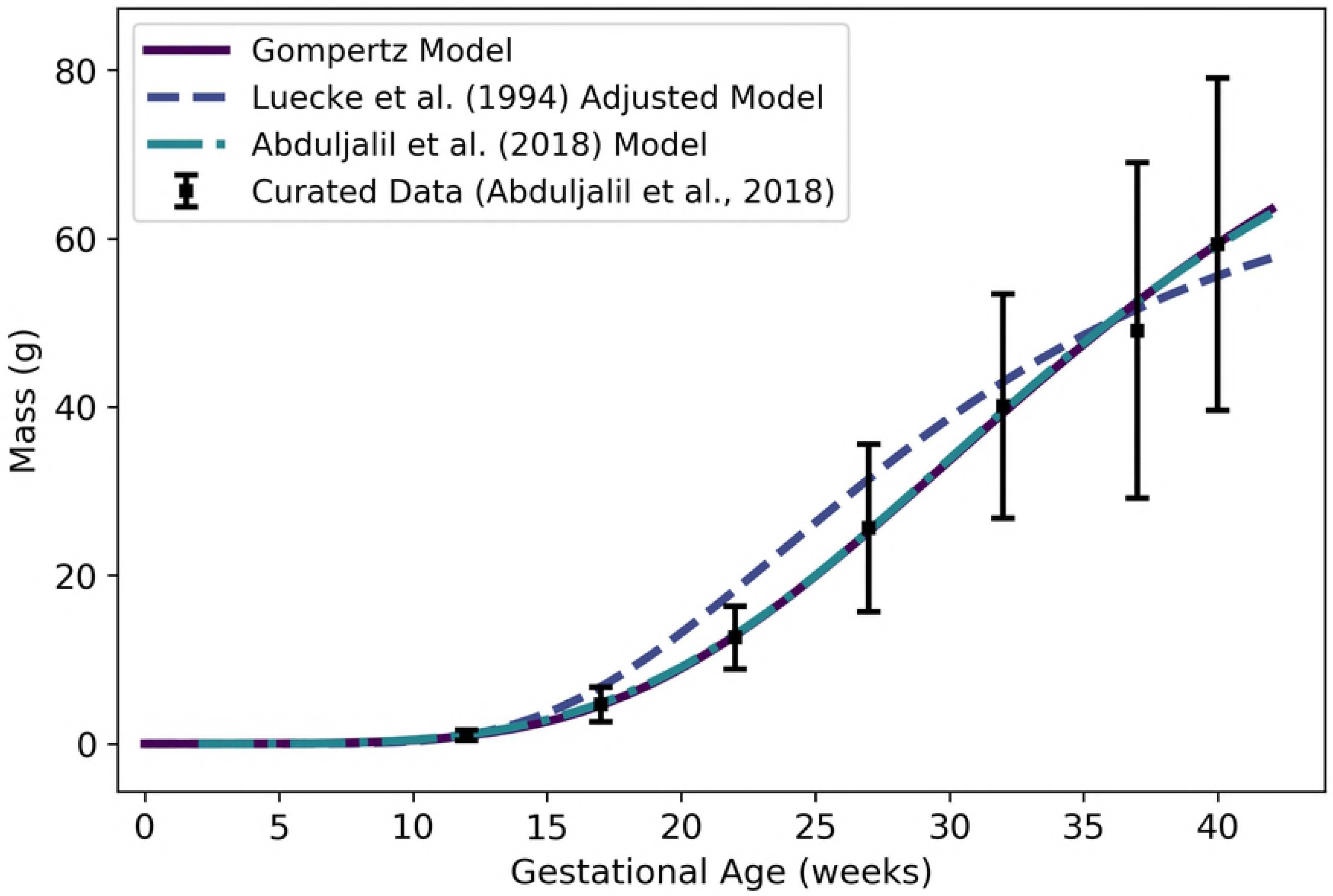
Fetal lung mass vs. gestational age. The Gompertz model (solid line) given by Equation 29 was selected as the most parsimonious model in our analysis. The model of Luecke et al. [15], which is a power law model based on fetal mass, was calibrated using a different data set [25]. The version of the Luecke et al. [15] model depicted here has been adjusted as described in the text. Note that our Gompertz model and the model of Abduljalil et al. [31] are virtually indistinguishable in this plot.

The model of Luecke et al. [15] depicted in Figure 18 is a Luecke power law model (cf. Table 2) with coefficient values that have been modified from those that the authors show in their Table 3. When applied exactly as specified, that model over-predicts fetal lung masses by an order of magnitude, so we hypothesized that the first coefficient printed in their table (*θ*_0_ = 0.09351) may have been off by a factor of 10. We therefore decreased that coefficient by a factor of 10 (*θ*_0_ = 0.009351) to obtain the model version (“Luecke et al. (1994) Adjusted Model”) shown in Figure 18.

Since the mean density of human lung tissue is 1.05 g/mL [33], the *volume* (mL) of the fetal lung tissue an be computed as

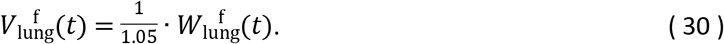

#### Thyroid

We used data curated by Abduljalil et al. [31] to calibrate various models for the thyroid mass of a human fetus during gestation. The Gompertz model given by

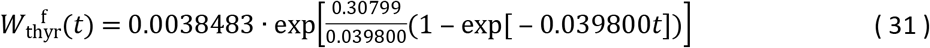

was selected as the most parsimonious model for fetal thyroid *mass* (g) at gestational age *t* (weeks). We remark that the cubic growth model yielded a slightly lower AIC, but the AIC difference (−334.07 vs. - 334.16) is too small to recommend one model over the other. Because the Gompertz model yields strictly positive values for gestational ages greater than or equal to zero, we selected that model over the cubic growth model.

Table 21 shows the maximum likelihood estimates of the parameter values for all models considered along with the associated log-likelihood and AIC values. The Gompertz model of Equation 31, two published models [15,31], and the curated summary data [31] are shown in Figure 19.

**Table 21.** Fetal thyroid mass models (g vs. fetal mass in g for power law models, g vs. gestational age in weeks for all other models). For each model considered, the maximum likelihood parameter estimates 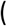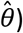, log-likelihood 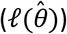, and AIC are provided. The row describing the selected model is shown in boldface.

| Model | $\hat{\theta}_0$ | $\hat{\theta}_1$ | $\hat{\theta}_2$ | $\hat{\theta}_3$ | $\ell(\hat{\theta})$ | AIC |
| --- | --- | --- | --- | --- | --- | --- |
| Linear Growth | 0.014960 | — | — | — | 28.2108 | -54.42 |
| Quadratic Growth | -0.015229 | 0.0014665 | — | — | 168.39 | -332.78 |
| Cubic Growth | -0.0029982 | 0.00043792 | $1.9641 \times 10^{-5}$ | — | 170.08 | -334.16 |
| **Gompertz** | **0.0038483** | **0.30799** | **0.039800** | — | **170.035** | **-334.07** |
| Logistic | 2.6022 | 0.14914 | 34.613 | — | 169.644 | -333.29 |
| Power Law | 0.0050755 | 0.71057 | — | — | 164.447 | -324.89 |
| Luecke Power Law | 0.10683 | -0.25045 | 0.073247 | — | 169.579 | -333.16 |

**Figure 19.**
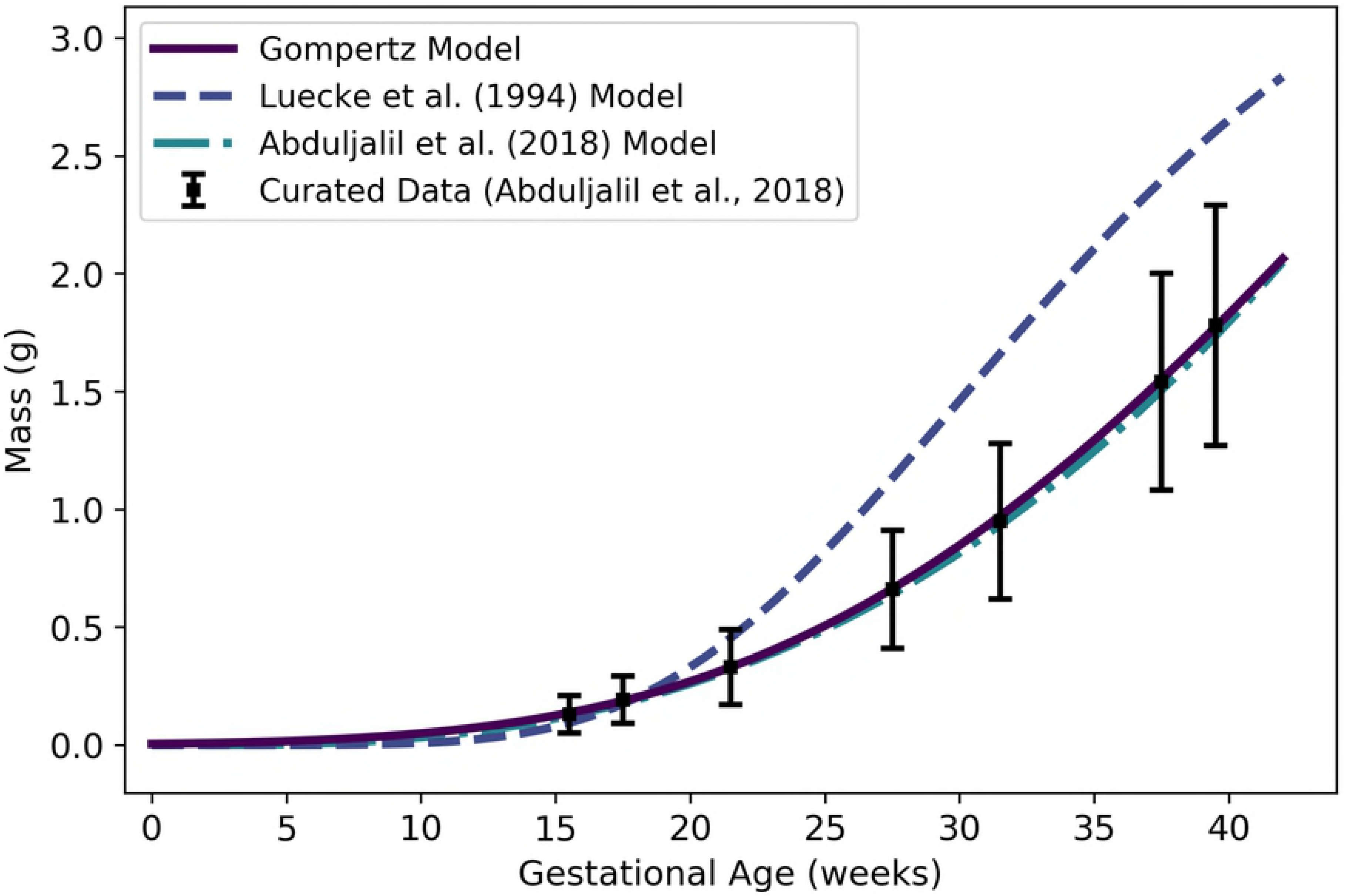
Fetal thyroid mass vs. gestational age. The Gompertz model (solid line) given by Equation 31 was selected as the most parsimonious model in our analysis. The model of Luecke et al. [15], which is a power law model based on fetal mass, was calibrated using a different data set [25].

Since the mean density of human thyroid tissue is 1.05 g/mL [33], the *volume* (mL) of the fetal thyroid can be computed as

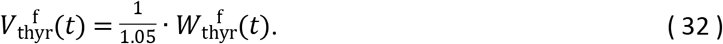

#### Gut

We used data curated by Abduljalil et al. [31] to calibrate various models for mass of the gastrointestinal tract (or “gut”) of a human fetus during gestation. The Gompertz model given by

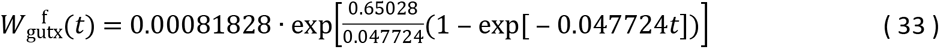

was selected as the most parsimonious model for fetal thyroid *mass* (g) at gestational age *t* (weeks). Table 22 shows the maximum likelihood estimates of the parameter values for all models considered along with the associated log-likelihood and AIC values. The Gompertz model of Equation 33, two published models [15,31], and the curated summary data [31] are shown in Figure 20.

**Table 22.** Fetal gut mass models (g vs. fetal mass in g for power law models, g vs. gestational age in weeks for all other models). For each model considered, the maximum likelihood parameter estimates 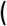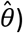, log-likelihood 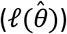, and AIC are provided. The row describing the selected model is shown in boldface.

| Model | $\hat{\theta}_0$ | $\hat{\theta}_1$ | $\hat{\theta}_2$ | $\hat{\theta}_3$ | $\ell(\hat{\theta})$ | AIC |
| --- | --- | --- | --- | --- | --- | --- |
| Linear Growth | 0.0059272 | — | — | — | -125.99 | 253.98 |
| Quadratic Growth | -0.10234 | 0.015358 | — | — | -58.46 | 120.93 |
| Cubic Growth | 0.14024 | -0.032080 | 0.0018327 | — | -11.68 | 29.37 |
| **Gompertz** | **0.00081828** | **0.65028** | **0.047724** | — | **-7.3947** | **20.79** |
| Logistic | 81.648 | 0.30673 | 31.545 | — | -15.16 | 36.32 |
| Power Law | 0.026604 | 0.93314 | — | — | -20.31 | 44.62 |
| Luecke Power Law | 0.038437 | 0.48413 | 0.056261 | — | -9.00 | 23.99 |

**Figure 20.**
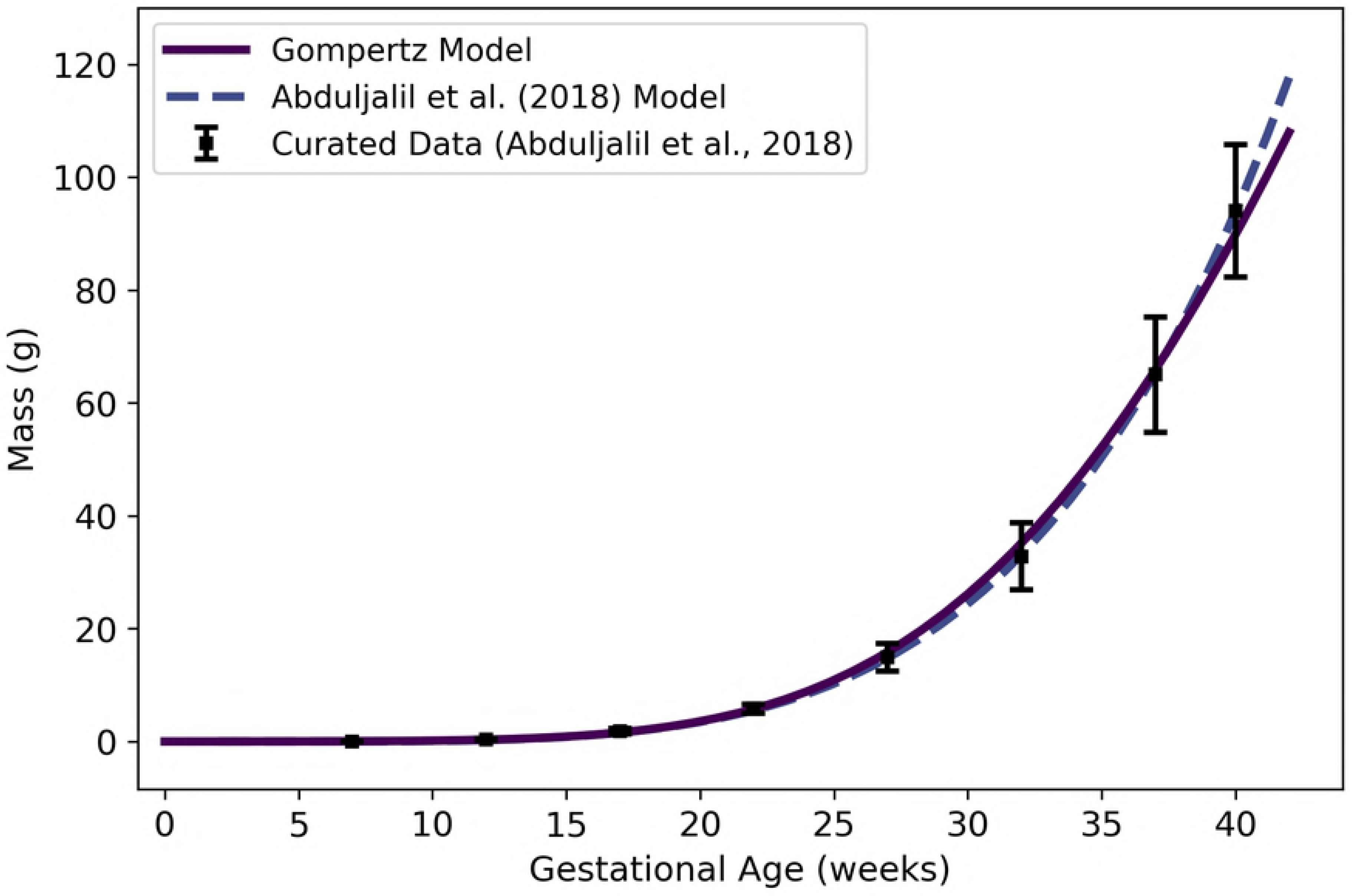
Fetal gut mass vs. gestational age. The Gompertz model (solid line) given by Equation 33 was selected as the most parsimonious model in our analysis.

Since the mean density of human gut tissue is 1.045 g/mL [33], the *volume* of the fetal gut (in mL) can be computed as

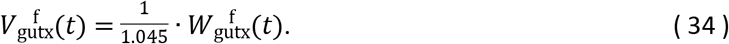

#### Rest of Body

We used the principle of mass balance to obtain a formula for the volume of a “rest of body” compartment comprising all mass in the fetal body that has not been accounted for in one of the specific compartments already described. The volume (mL) of the rest of body can be calculated as

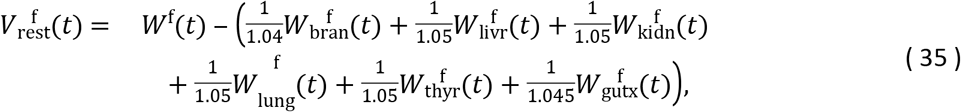

where *W*^f^, 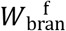, 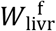, 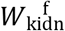, 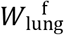, 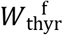, and 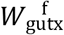 are given by Equations 2, 23b, 25b, 27b, 29b, 31b, and 33 respectively, and the scalars represent appropriate densities (cf. Table 7). Note that we assume the average density of the fetus is 1 g/mL throughout gestation, so the total mass (g) of the fetus equals the total volume (mL) of the fetus. Equation 35 yields negative values until about 8 weeks of gestational age, so it should not be used for predictions before that time. In fact, because several of the models upon which Equation 35 depends were derived from data sets that contain no observations from the first trimester, we recommend that this model only be applied for gestational ages greater than 13 weeks. As shown in Figure 21, Equation 35 results in volumes for the fetal rest of body compartment that increase from about 15 mL at 13 weeks to about 2800 mL at 40 weeks.

**Figure 21.**
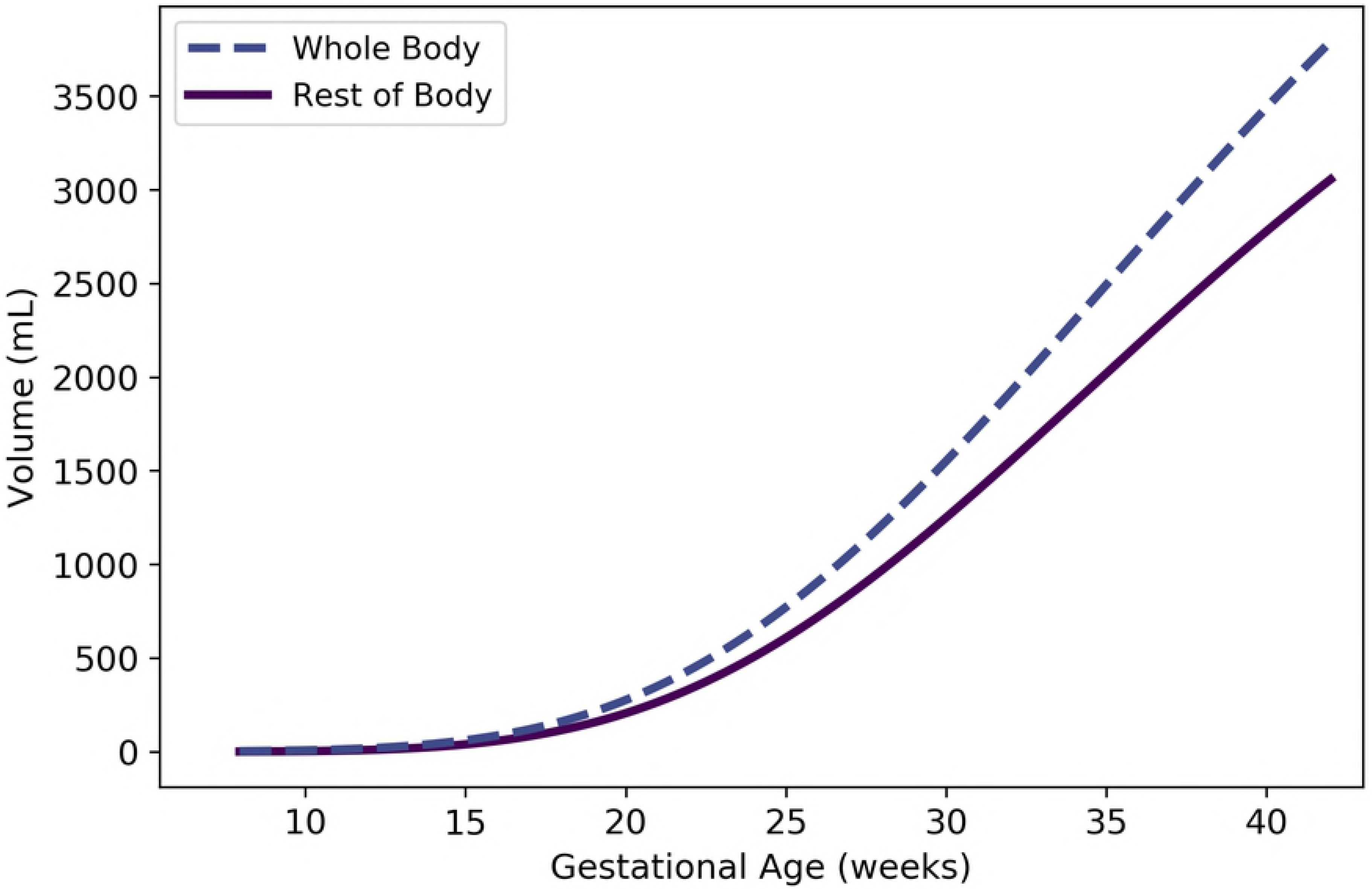
Fetal rest of body vs. gestation age (cf. Equation 35). The volume of the whole fetal body (cf. Equation 2) is shown for comparison.

### Fetal Blood Flow Rates

Doppler ultrasound studies provide a non-invasive means for determining cardiac output and distribution of blood flow in an organism at various stages of development. We examined several studies that rely on such measurements to obtain formulas for blood flow rates to fetal tissues and through various components of the fetal circulatory system.

#### Right Ventricle

We extracted data from Figure 5A of Kiserud et al. [36] and used it to calibrate various models for blood flow rate through the right ventricle of the fetal heart. The logistic model given by

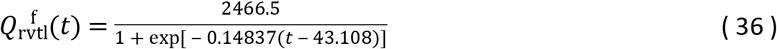

was selected as the most parsimonious model for fetal right ventricle flow (mL/min) at gestational age *t* (weeks). Table 23 shows the maximum likelihood estimates of the parameter values for all models considered along with the associated log-likelihood and AIC values. Figure 22 shows the logistic model of Equation 36 along with the published model of Kiserud et al. [36].

**Table 23.** Fetal right ventricle blood flow models (mL/min vs. fetal mass in g for power law models, mL/min vs. gestational age in weeks for all other models). For each model considered, the maximum likelihood parameter estimates 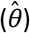, log-likelihood 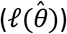, and AIC are provided. The row describing the selected model is shown in boldface.

| Model | $\hat{\theta}_0$ | $\hat{\theta}_1$ | $\hat{\theta}_2$ | $\hat{\theta}_3$ | $\ell(\hat{\theta})$ | AIC |
| --- | --- | --- | --- | --- | --- | --- |
| Linear Growth | 3.265 | — | — | — | -1563.53 | 3129.07 |
| Quadratic Growth | -19.697 | 1.0498 | — | — | -964.06 | 1932.13 |
| Cubic Growth | 11.755 | -1.0908 | 0.034829 | — | -940.55 | 1887.09 |
| Gompertz | 1.0138 | 0.27387 | 0.025665 | — | -940.51 | 1887.02 |
| **Logistic** | **2466.5** | **0.14837** | **43.108** | — | **-940.14** | **1886.29** |
| Power Law | 0.024935 | 1.2908 | — | — | -956.00 | 1915.99 |
| Luecke Power Law | 16619 | -2.2943 | 0.23852 | — | -941.29 | 1888.57 |

**Figure 22.**
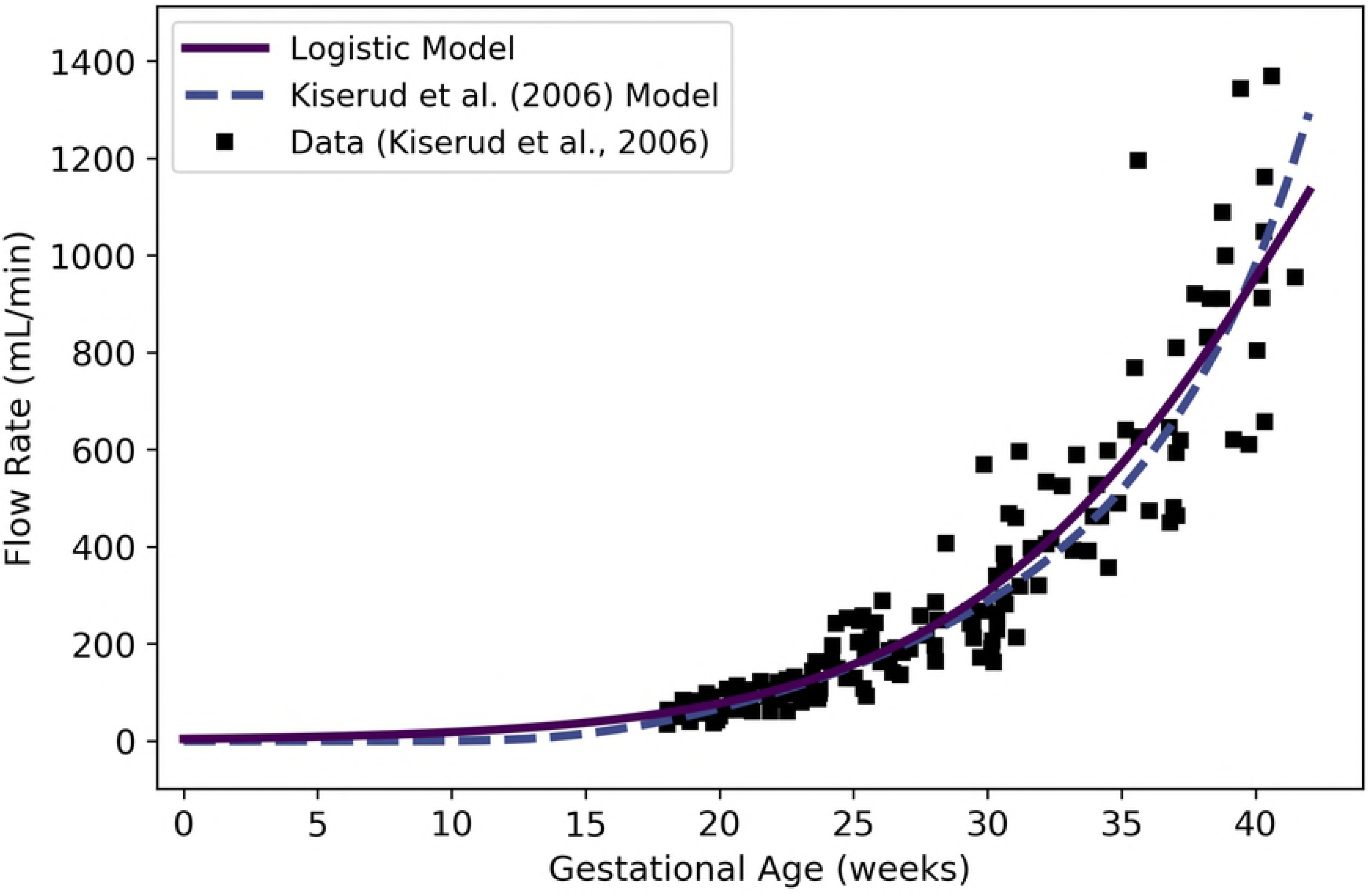
Fetal blood flow through the right ventricle vs. gestational age. The logistic model (solid line) given by Equation 36 was selected as the most parsimonious model in our analysis. The model of Kiserud et al. [36] was calibrated using the same data set [36] used by us.

#### Left Ventricle

We extracted data from Figure 4A of Kiserud et al. [36] and used it to calibrate various models for blood flow rate through the left ventricle of the fetal heart. The logistic model given by

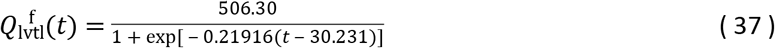

was selected as the most parsimonious model for fetal left ventricle flow (mL/min) at gestational age *t* (weeks). Table 24 shows the maximum likelihood estimates of the parameter values for all models considered along with the associated log-likelihood and AIC values. Figure 23 shows the logistic model of Equation 37 along with the published model of Kiserud et al. [36].

**Table 24.** Fetal left ventricle blood flow models (mL/min vs. fetal mass in g for power law models, mL/min vs. gestational age in weeks for all other models). For each model considered, the maximum likelihood parameter estimates 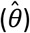, log-likelihood 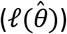, and AIC are provided. The row describing the selected model is shown in boldface.

| Model | $\hat{\theta}_0$ | $\hat{\theta}_1$ | $\hat{\theta}_2$ | $\hat{\theta}_3$ | $\ell(\hat{\theta})$ | AIC |
| --- | --- | --- | --- | --- | --- | --- |
| Linear Growth | 8.2127 | — | — | — | -1328.70 | 2659.41 |
| Quadratic Growth | -6.4372 | 0.46894 | — | — | -1002.69 | 2009.38 |
| Cubic Growth | -18.700 | 1.3115 | -0.013824 | — | -992.93 | 1991.86 |
| Gompertz | $2.4461 \times 10^{-5}$ | 1.5891 | 0.092460 | — | -989.10 | 1984.20 |
| **Logistic** | **506.30** | **0.21916** | **30.231** | — | **-985.42** | **1976.84** |
| Power Law | 0.42909 | 0.86053 | — | — | -991.64 | 1987.28 |
| Luecke Power Law | 0.0024511 | 2.2736 | -0.095998 | — | -989.07 | 1984.14 |

**Figure 23.**
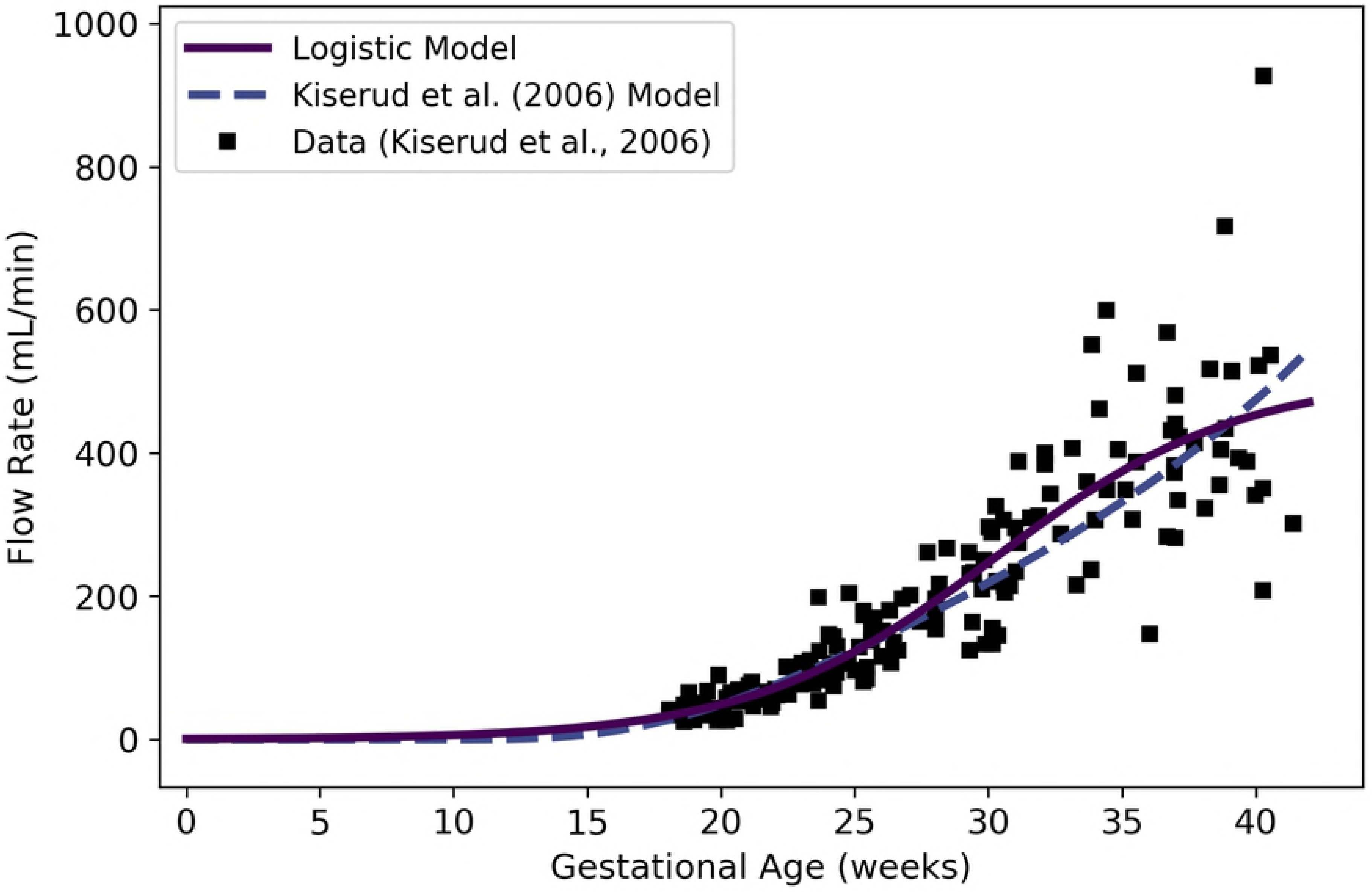
Fetal blood flow through the left ventricle vs. gestational age. The logistic model (solid line) given by Equation 37 was selected as the most parsimonious model in our analysis. The model of Kiserud et al. [36] was calibrated using the same data set [36] used by us.

#### Ductus Arteriosus

We extracted data from Figure 8 of Mielke and Benda [37] and used it to calibrate various models for blood flow rate through ductus arteriosus of the human fetus. The logistic model given by

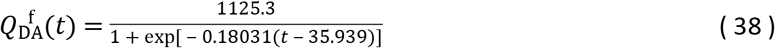

was selected as the most parsimonious model for fetal ductus arteriosus flow (mL/min) at gestational age *t* (weeks). Table 25 shows the maximum likelihood estimates of the parameter values for all models considered along with the associated log-likelihood and AIC values. Figure 24 shows the logistic model of Equation 38 and the data of Mielke and Benda [37].

**Table 25.** Fetal ductus arteriosus blood flow models (mL/min vs. fetal mass in g for power law models, mL/min vs. gestational age in weeks for all other models). For each model considered, the maximum likelihood parameter estimates 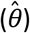, log-likelihood 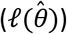, and AIC are provided. The row describing the selected model is shown in boldface.

| Model | $\hat{\theta}_0$ | $\hat{\theta}_1$ | $\hat{\theta}_2$ | $\hat{\theta}_3$ | $\ell(\hat{\theta})$ | AIC |
| --- | --- | --- | --- | --- | --- | --- |
| Linear Growth | 13.401 | — | — | — | -1714.79 | 3431.58 |
| Quadratic Growth | -13.001 | 0.79207 | — | — | -1258.10 | 2520.21 |
| Cubic Growth | -3.2054 | 0.079383 | 0.012051 | — | -1252.88 | 2511.76 |
| Gompertz | 0.011836 | 0.71885 | 0.058724 | — | -1249.89 | 2505.78 |
| **Logistic** | **1125.3** | **0.18031** | **35.939** | — | **-1247.00** | **2499.99** |
| Power Law | 0.068851 | 1.1419 | — | — | -1251.60 | 2507.2 |
| Luecke Power Law | 23.030 | -0.41319 | 0.10352 | — | -1249.07 | 2504.15 |

**Figure 24.**
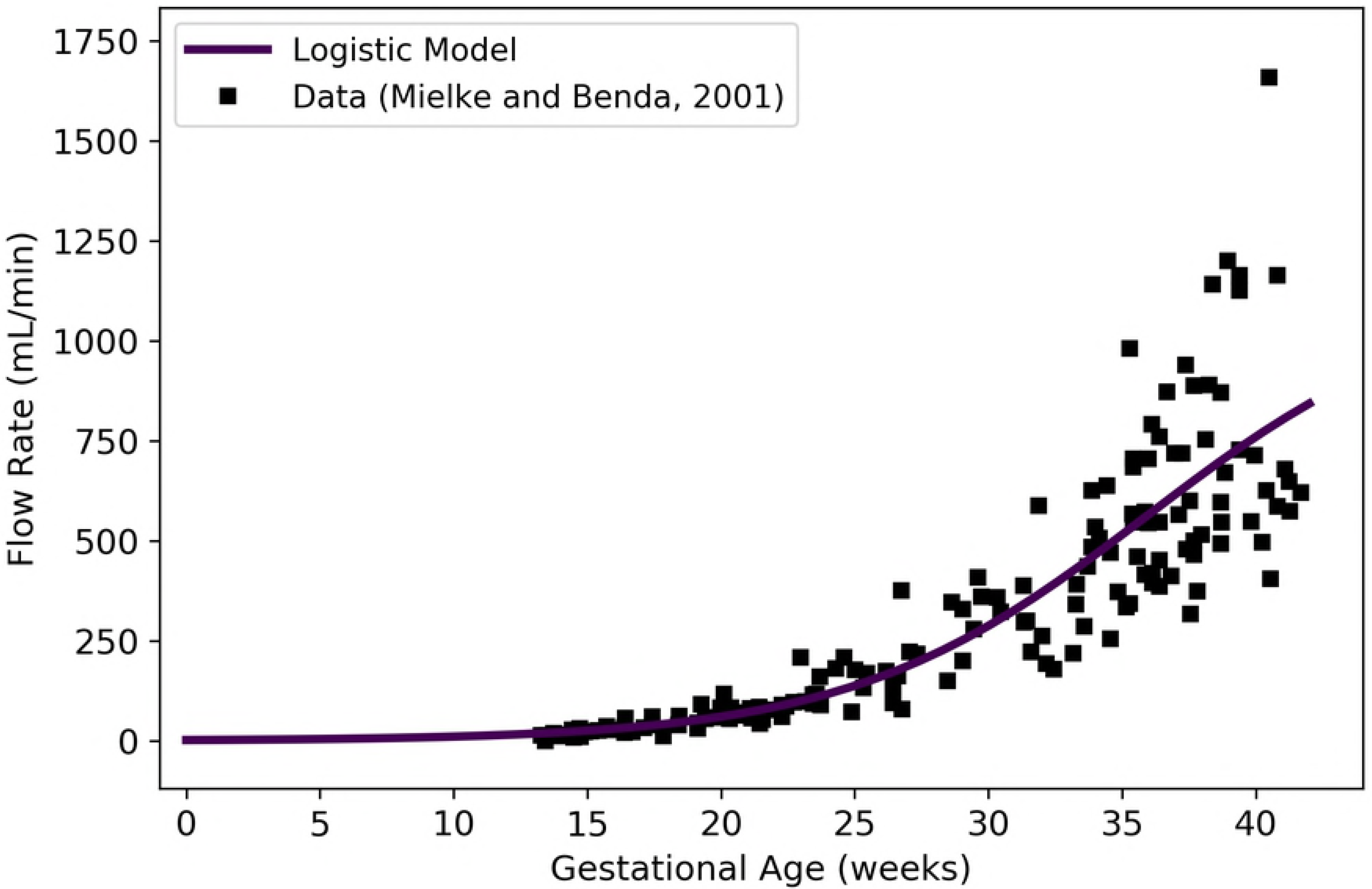
Fetal blood flow through the ductus arteriosus vs. gestational age. The logistic model (solid line) given by Equation 38 was selected as the most parsimonious model in our analysis, which utilized data of Mielke and Benda [37].

#### Arterial Blood

We define the flow rate into the fetal “arterial blood compartment” as the flow through the aorta just beyond its junction with the ductus arteriosus [57]. This flow (mL/min) can be computed as

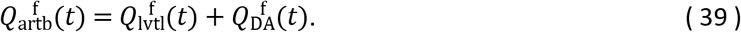

Note that this formula represents a composite model, and it is not based directly upon measured time-course data for flow into the fetal arterial blood compartment.

#### Lung

The flow rate into the lungs equals the flow rate out of the left ventricle minus the flow that is shunted away from the lungs by the ductus arteriosus [57]. This flow (mL/min) is given by

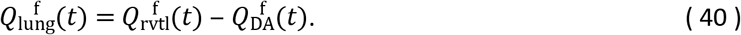

Note that this formula represents a composite model, and it is not based directly upon measured time-course data for flow into the fetal lung.

#### Foramen Ovale

Assuming a conservation of flow, the total flow rate returning to the heart from the body via the “venous blood compartment” should be equal to the total flow rate leaving the heart and entering the “arterial blood compartment”. This flow enters the right atrium of the heart, and part of the flow is directed to the right ventricle, while the remaining part is directed through the foramen ovale [57]. Thus, the flow (mL/min) through the foramen ovale can be computed as

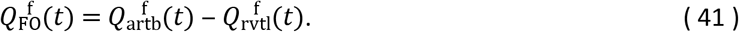

Note that this formula represents a composite model, and it is not based directly upon measured time-course data for flow through the foramen ovale.

#### Placenta

We extracted data from Figure 5A of Kiserud et al. [38] and used it to calibrate various models for the fetal blood flow rate through the placenta. The data depicted in the aforementioned figure are for blood flow in the “intra-abdominal umbilical vein”, and so they measure rates of blood flowing from the placenta toward the liver and the ductus venosus [38]. The logistic model given by

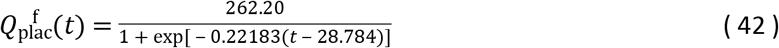

was selected as the most parsimonious model for fetal flow (mL/min) through the placenta at gestational age *t* (weeks).

Table 26 shows the maximum likelihood estimates of the parameter values for all models considered along with the associated log-likelihood and AIC values. Figure 25 shows the logistic model of Equation 42 along with the published model of Kiserud et al. [38].

**Table 26.**
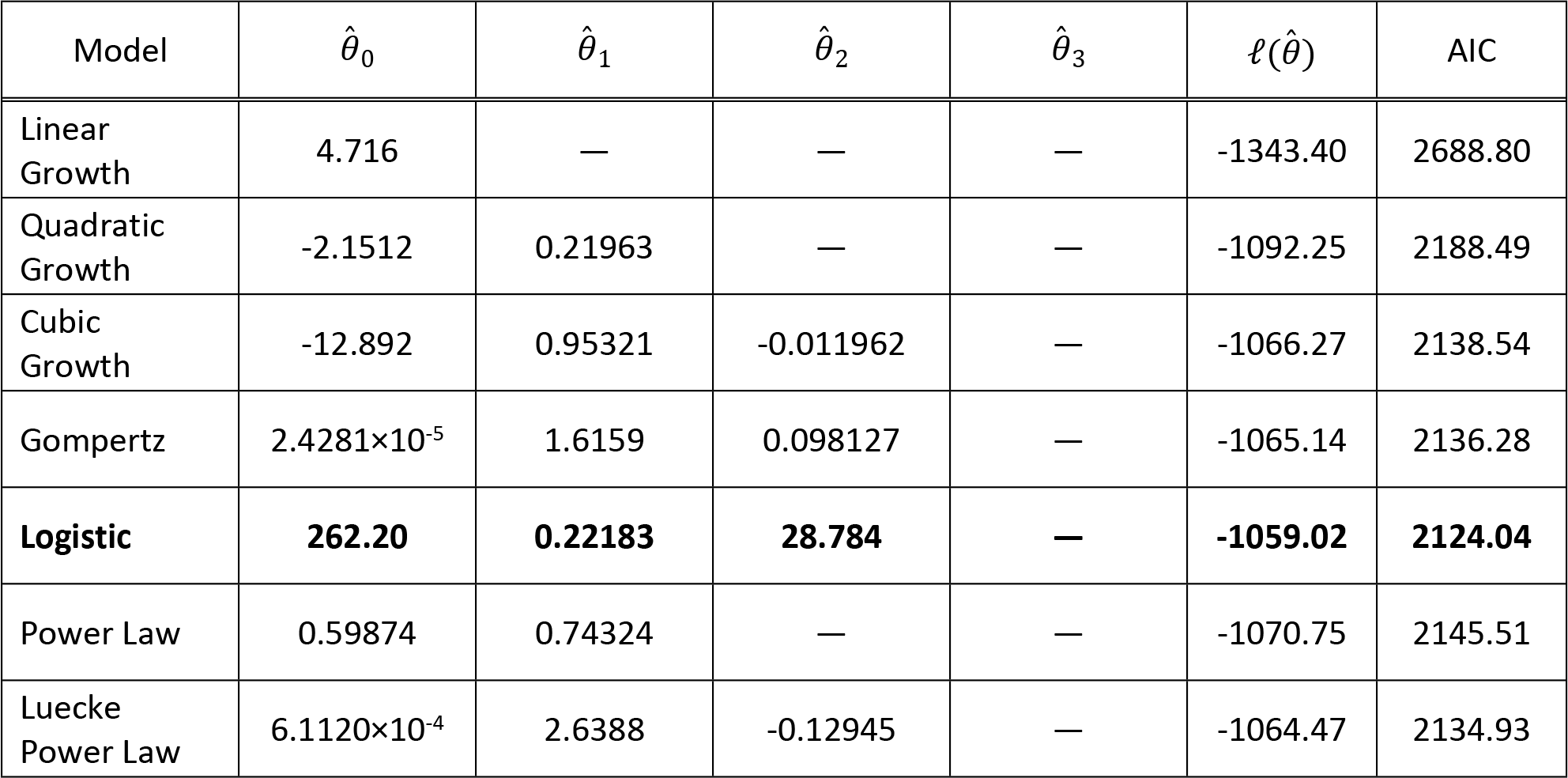
Models for fetal blood flow to the placenta (mL/min vs. fetal mass in g for power law models, mL/min vs. gestational age in weeks for all other models). For each model considered, the maximum likelihood parameter estimates 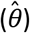, log-likelihood 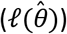, and AIC are provided. The row describing the selected model is shown in boldface.

**Figure 25.**
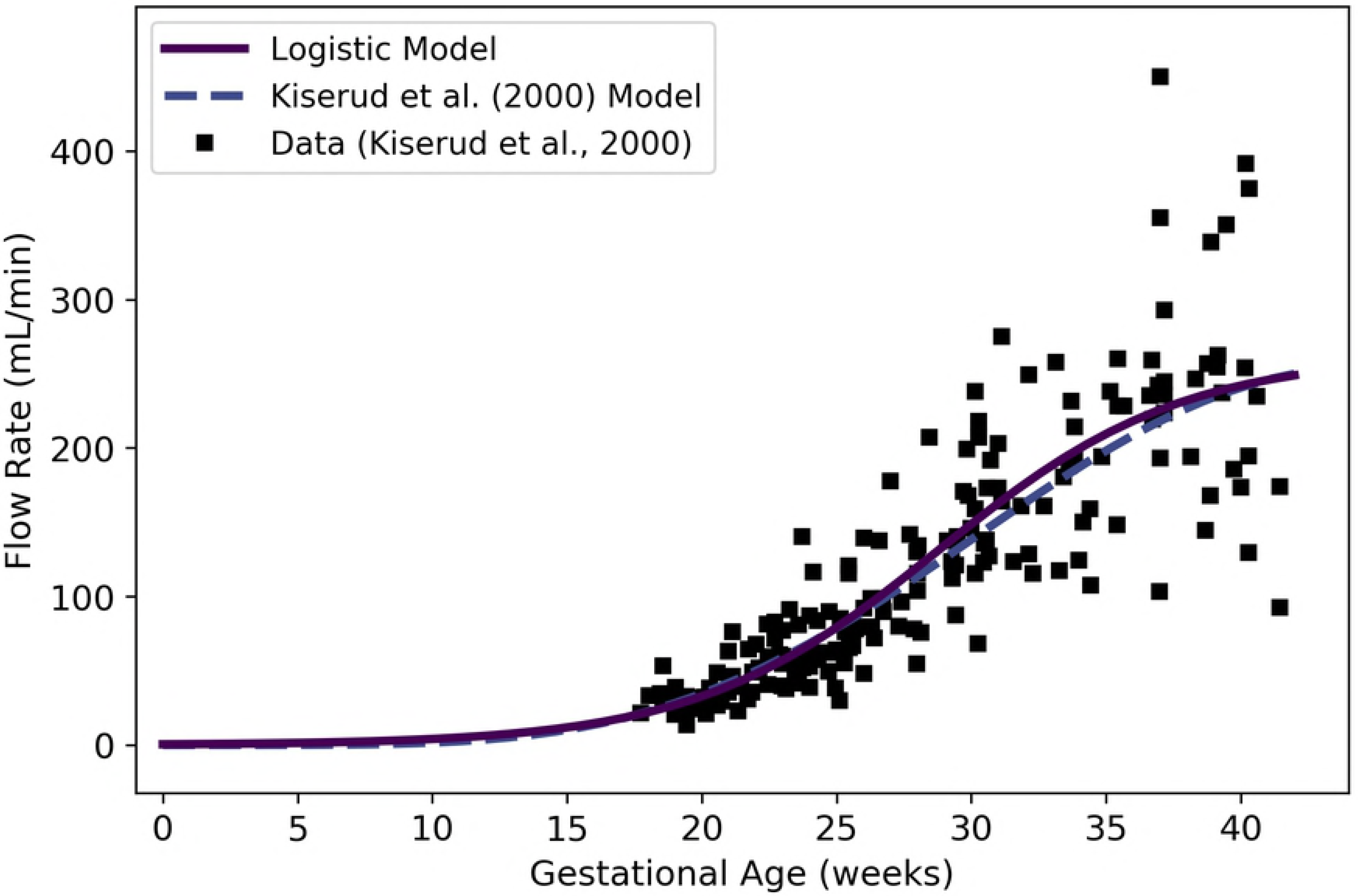
Fetal blood flow through the placenta vs. gestational age. The logistic model (solid line) given by Equation 42 was selected as the most parsimonious model in our analysis. The model of Kiserud et al. [38] was calibrated using the same data set [38] used by us.

#### Ductus Venosus

We extracted data from Figure 5B of Kiserud et al. [38] and used it to calibrate various models for the fetal blood flow rate through the ductus venosus. The Gompertz model given by

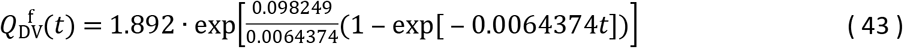

was selected as the most parsimonious model for fetal blood flow (mL/min) through the ductus venosus at gestational age *t* (weeks). Table 27 shows the maximum likelihood estimates of the parameter values for all models considered along with the associated log-likelihood and AIC values. Figure 26 shows the Gompertz model of Equation 43 along with the published model of Kiserud et al. [38].

**Table 27.** Models for fetal blood flow through the ductus venosus (mL/min vs. fetal mass in g for power law models, mL/min vs. gestational age in weeks for all other models). For each model considered, the maximum likelihood parameter estimates 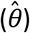, log-likelihood 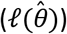, and AIC are provided. The row describing the selected model is shown in boldface.

| Model | $\hat{\theta}_0$ | $\hat{\theta}_1$ | $\hat{\theta}_2$ | $\hat{\theta}_3$ | $\ell(\hat{\theta})$ | AIC |
| --- | --- | --- | --- | --- | --- | --- |
| Linear Growth | 1.0214 | — | — | — | -1578.25 | 3158.5 |
| Quadratic Growth | -0.41055 | 0.046294 | — | — | -1330.9 | 2665.8 |
| Cubic Growth | 0.39477 | -0.0095095 | $9.2087 \times 10^{-4}$ | — | -1327.59 | 2661.18 |
| **Gompertz** | **1.8920** | **0.098249** | **0.0064374** | — | **-1326.87** | **2659.75** |
| Logistic | 2072.9 | 0.081565 | 82.890 | — | -1327.09 | 2660.18 |
| Power Law | 0.14986 | 0.72500 | — | — | -1347.34 | 2698.68 |
| Luecke Power Law | 180.87 | -1.2763 | 0.13957 | — | -1335.16 | 2676.32 |

**Figure 26.**
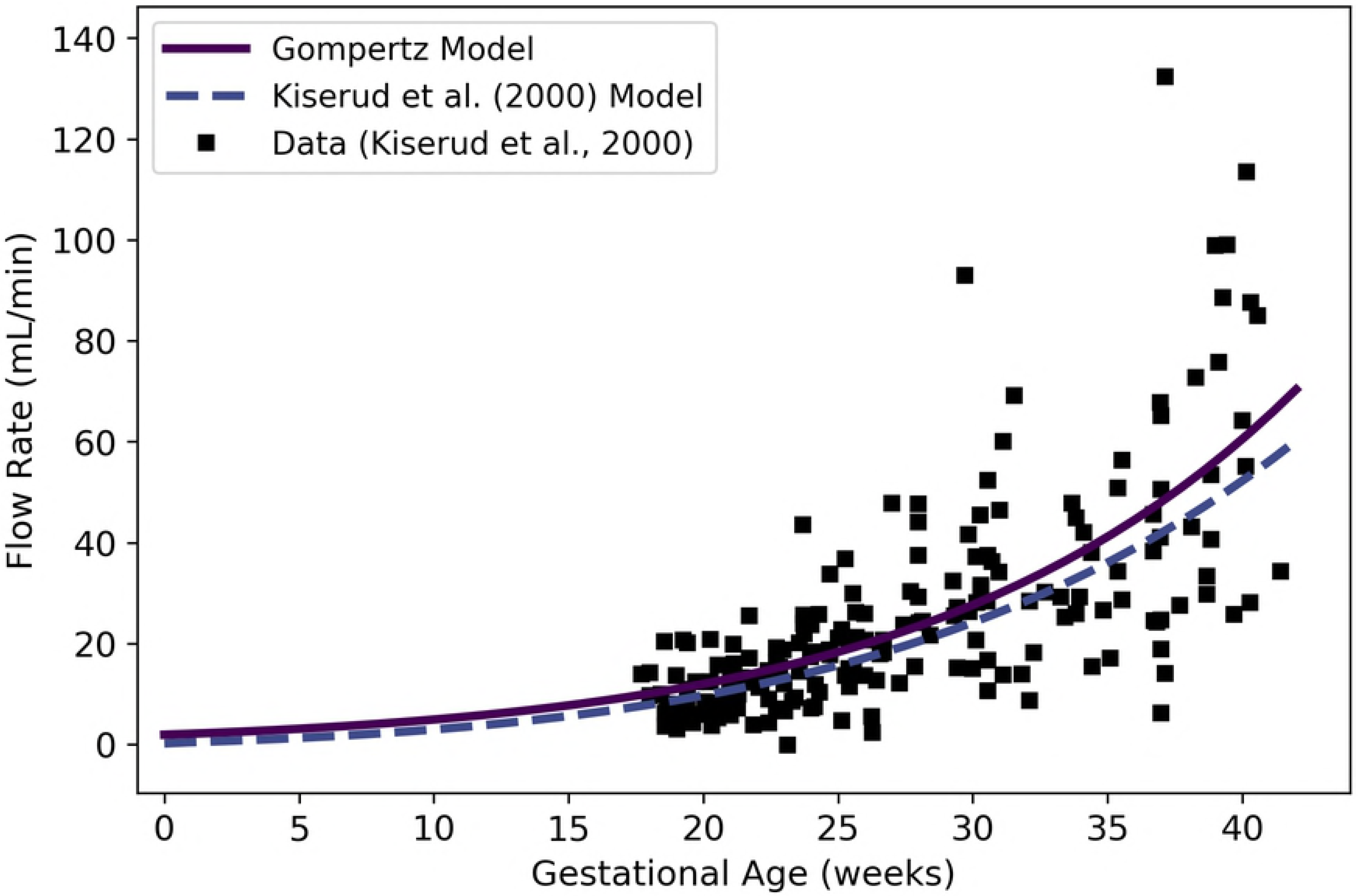
Fetal blood flow through the ductus venosus vs. gestational age. The Gompertz model (solid line) given by Equation 43 was selected as the most parsimonious model in our analysis. The model of Kiserud et al. [38] was calibrated using the same data set [38] used by us.

When using the models for flow to the placenta (Equation 42) and through the ductus venosus (Equation 43) together, one should exercise caution. The ductus venosus effectively diverts a portion of the blood traveling through the umbilical vein directly to the venous blood compartment, whereas the rest of that blood is carried to the liver [38]. Thus, the flow through the ductus venosus should always be less than the flow to (and from) the placenta. As shown in Figure 27, our models ensure that 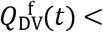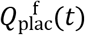 for gestational ages *t* greater than 12 weeks, but for some earlier gestational ages this is not the case.

**Figure 27.**
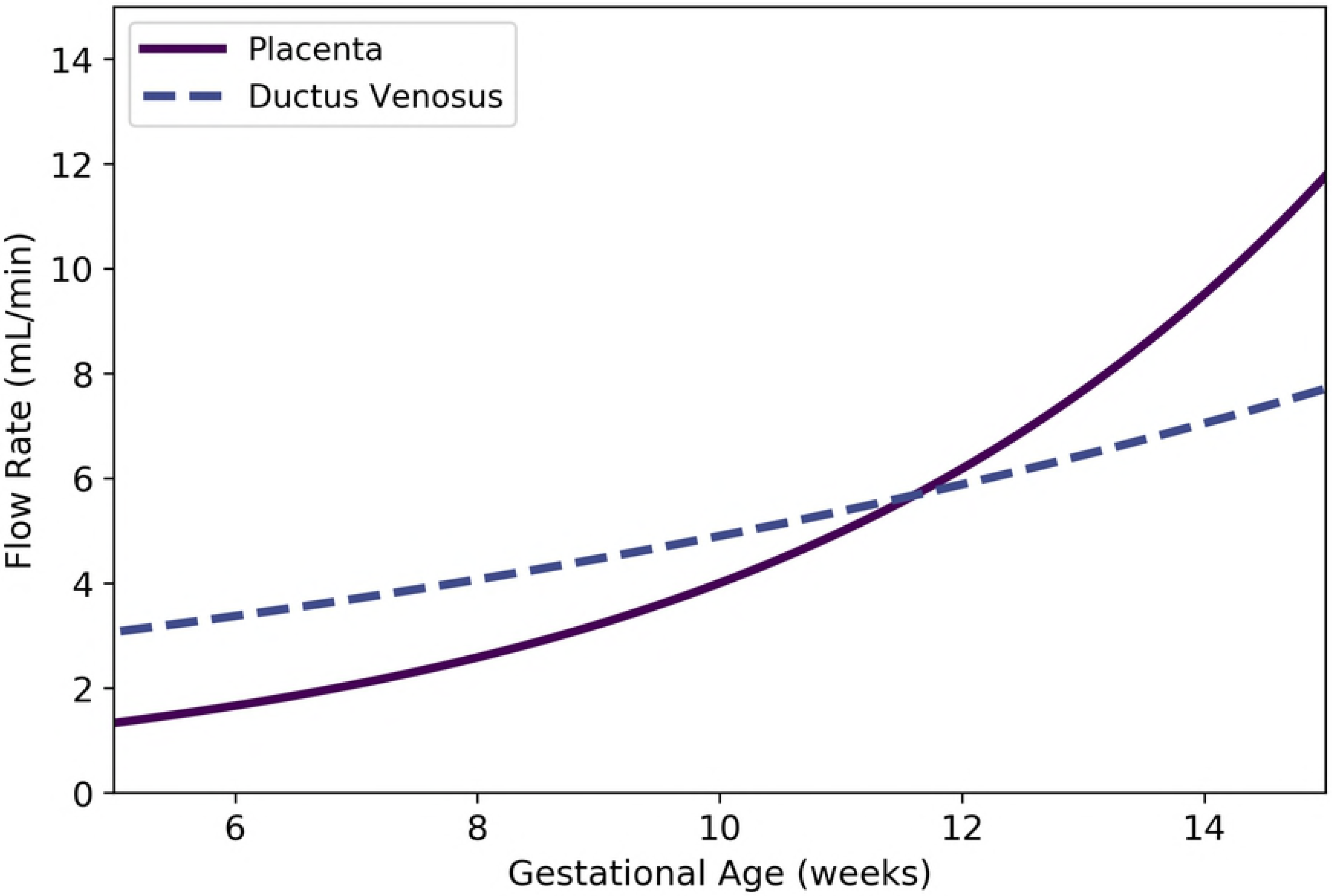
Comparison of flow rates through the placenta and through the ductus venosus vs. gestational age (cf. Equations 42 and 43). Because blood flow through the ductus venosus should always be less than blood flow through the placenta, these models should not be used prior to gestational age 12 weeks.

#### Other Specific Compartments

We compute flow rates to other specific compartments in the fetus using proportions of the total flow to the arterial blood compartment, 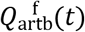, which is given in Equation 39. Rudolph et al. [58] measured blood flow rates to various organs in 33 previable human fetuses with masses ranging from 12 to 272 grams (and estimated gestational ages between 10 and 20 weeks). Using this data, we generated the mean flow rate data shown in Table 28. To compute the fetal blood flows to the various sites other than the placenta (for which we already have Equation 42), we made the following assumptions:

- The proportion of blood flow to the placenta is not a constant, but depends on gestational age. This proportion can be computed as 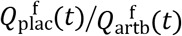 using Equations 39 and 42.
- All arterial blood that does not flow to the placenta flows to the other tissues and tissue groups listed in Table 28. The total proportion of arterial blood flowing to these other tissues can be computed as 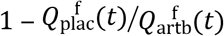
- The blood flow rates to compartments other than the placenta maintain constant proportions according to the values listed in Table 28. Since the total percentage of cardiac output flowing to tissues other than the placenta equals 75 (according to Table 28), all proportions from the table are scaled by this value.

Thus, blood flows (mL/min) to the gut, kidneys, and brain can be computed as

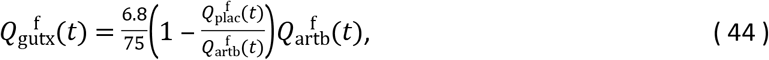

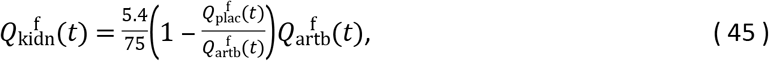

and

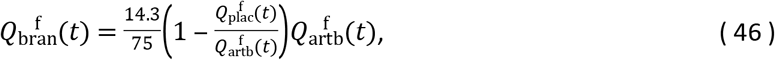

respectively.

**Table 28.** Average blood flow rates to various fetal tissues. The values shown indicate mean percentages of total cardiac output (% CO), and are based upon data of Rudolph et al. [58]. In some cases, these authors made multiple measurements for a single fetus; for the purposes of computing the means in this table we only used the first measurement for each fetus. For many of the individual fetuses measured by Rudolph et al. [58], the percentages summed to 99 or 101; this can be explained by the fact that percentage values were rounded to the nearest unit. In some cases, however, the percentages for an individual fetus summed to a total less than 99 (and, in one case, as low as 88). The reason for these larger discrepancies is unclear, but they were rare and we decided to use the values from all 33 fetuses to compute the averages reported here.

| Tissue | Relevant Symbol | % CO |
| --- | --- | --- |
| Placenta | $Q_{\text{plac}}^f$ | 23.9 |
| Gut | $Q_{\text{gutx}}^f$ | 6.8 |
| Kidneys | $Q_{\text{kidn}}^f$ | 5.4 |
| Adrenals | — | 4.8 |
| Lower Body | — | 27.1 |
| Heart | — | 3.0 |
| Brain | $Q_{\text{bran}}^f$ | 14.3 |
| Upper Body | — | 13.6 |
| Total |  | 98.9 |

Several fetal tissue compartments were not examined by Rudolph et al. [58], and are therefore not listed in Table 28. To determine blood flow rates to these fetal compartments, we relied on published *adult* blood flow proportions [33]. In a typical adult (averaging male and female values), the total proportion of cardiac output *not* flowing to the gut, kidneys, or brain is 54% [33]. (We assume that the “gut” includes the stomach and esophagus, the small intestine, and the large intestine.) Also, in the fetus, the proportion of the arterial blood flow 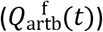 that goes to the gut, kidneys, and brain is 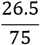 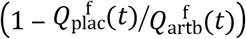 (where 26.5 is the sum of the percentages flowing to gut, kidneys, and brain in Table 28). Thus, the proportion of arterial blood flow not going to the gut, kidneys, brain, or placenta is 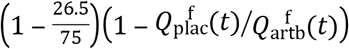. Since the adult proportions of cardiac output flowing to the liver and thyroid are 6.5% and 1.5%, respectively [33], we can estimate blood flows (mL/min) to the *fetal* liver and thyroid as

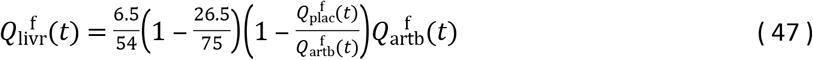

and

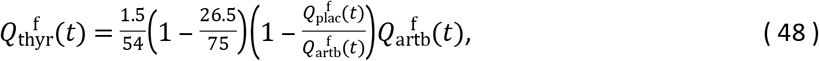

respectively.

#### Rest of Body

We used the principle of conservation of flow to obtain a formula for the blood flow rate to the fetal rest of body compartment. Thus, the flow rate to the rest of body compartment (in mL/min) is given by

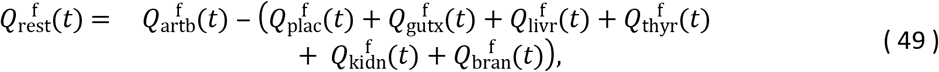

where *t* is the gestational age (weeks) and 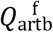, 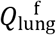, 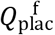, 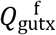, 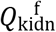, 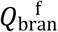, 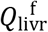, and 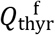 are given by Equations 39, 40, 42, 44, 45, 46, 47, and 48, respectively. As shown in Figure 28, Equation 49 results in flow rates to the fetal rest of body compartment that increase from approximately 12 mL/min at 13 weeks to approximately and 535 mL/min at 40 weeks. The model produces strictly positive values for all gestational ages greater than zero; however, because the models upon which Equation 49 depends were derived from data sets that contain no observations prior to the second trimester, we recommend that this model only be applied for gestational ages greater than 13 weeks.

**Figure 28.**
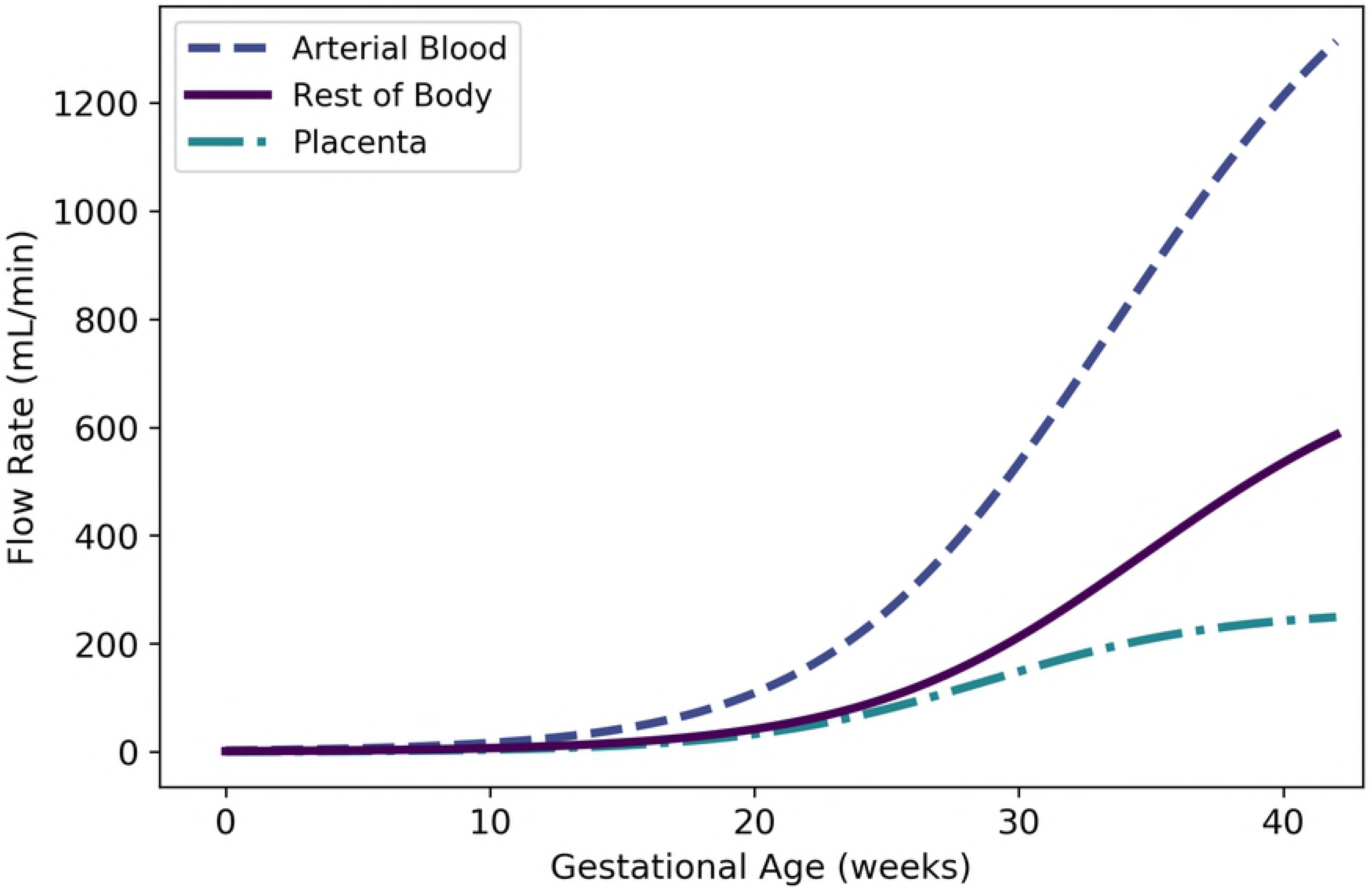
Fetal blood flow to the “rest of body” compartment vs. gestational age (cf. Equation 49). Fetal blood flow to the arterial blood compartment (cf. Equation 39) and the placenta (cf. Equation 42) are shown for comparison.

### Other Fetal Physiological Parameters

#### Hematocrit

Ohls (2011) reported means and standard deviations for hematocrits of fetuses at gestational ages 18-21, 22-25, 26-29, and “>30” weeks, as well as at “term”. To calibrate our models using these data, we assumed these periods correspond to 20, 24, 28, and 35 weeks of gestational age. Ohls also reported that “term hematocrit values range from 50% to 63%,” so we included an additional data point of 56.5% +/− 6.5% (the midpoint +/− half the difference of these two values) at 40 weeks. To incorporate an assumption that the embryo has no red blood cells at the time of conception (0 weeks gestational age), we included a data point for 0.0% +/− 4.0% at 0 weeks. As shown in Table 29, the quadratic growth model given by

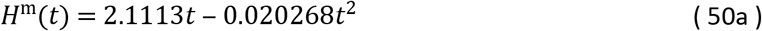

is the most parsimonious model based on the AIC; however, the cubic growth model

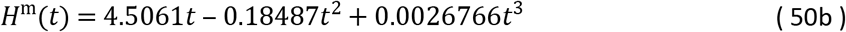

gives a better visual fit to the data and its AIC is only 0.8% larger. Both models allow for prediction of the fetal hematocrit (as a percentage) at gestational *t* (weeks) and are shown in Figure 29. The cubic model of Dallmann et al. [3] is also shown in Figure 29.

**Table 29.**
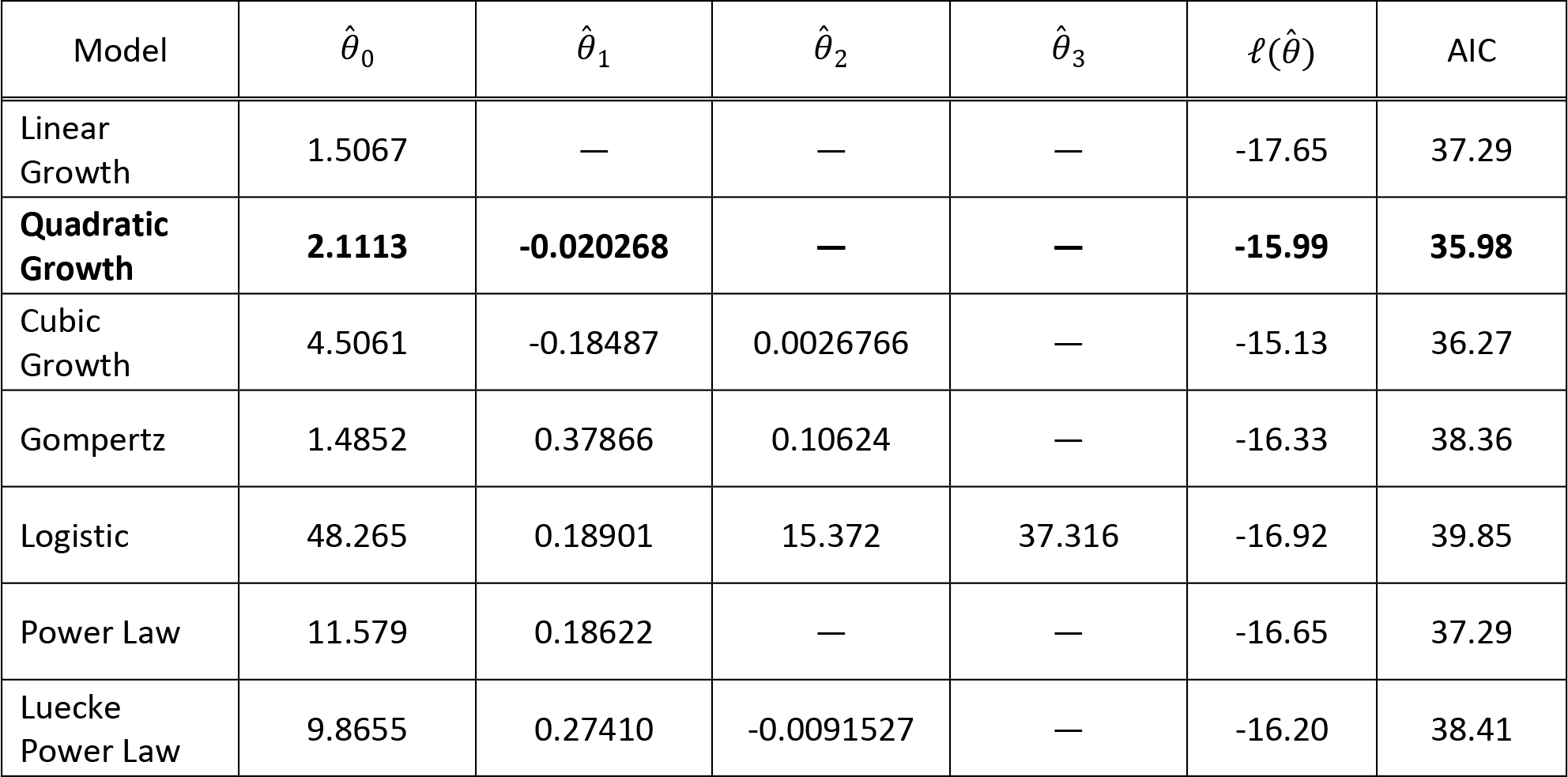
Fetal hematocrit models (percentage vs. gestational age in weeks). For each model considered, the maximum likelihood parameter estimates 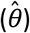, log-likelihood 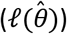, and AIC are provided. The row describing the selected model is shown in boldface.

| Model | $\hat{\theta}_0$ | $\hat{\theta}_1$ | $\hat{\theta}_2$ | $\hat{\theta}_3$ | $\ell(\hat{\theta})$ | AIC |
| --- | --- | --- | --- | --- | --- | --- |
| Linear Growth | 1.5067 | — | — | — | -17.65 | 37.29 |
| **Quadratic Growth** | **2.1113** | **-0.020268** | — | — | **-15.99** | **35.98** |
| Cubic Growth | 4.5061 | -0.18487 | 0.0026766 | — | -15.13 | 36.27 |
| Gompertz | 1.4852 | 0.37866 | 0.10624 | — | -16.33 | 38.36 |
| Logistic | 48.265 | 0.18901 | 15.372 | 37.316 | -16.92 | 39.85 |
| Power Law | 11.579 | 0.18622 | — | — | -16.65 | 37.29 |
| Luecke Power Law | 9.8655 | 0.27410 | -0.0091527 | — | -16.20 | 38.41 |

**Figure 29.**
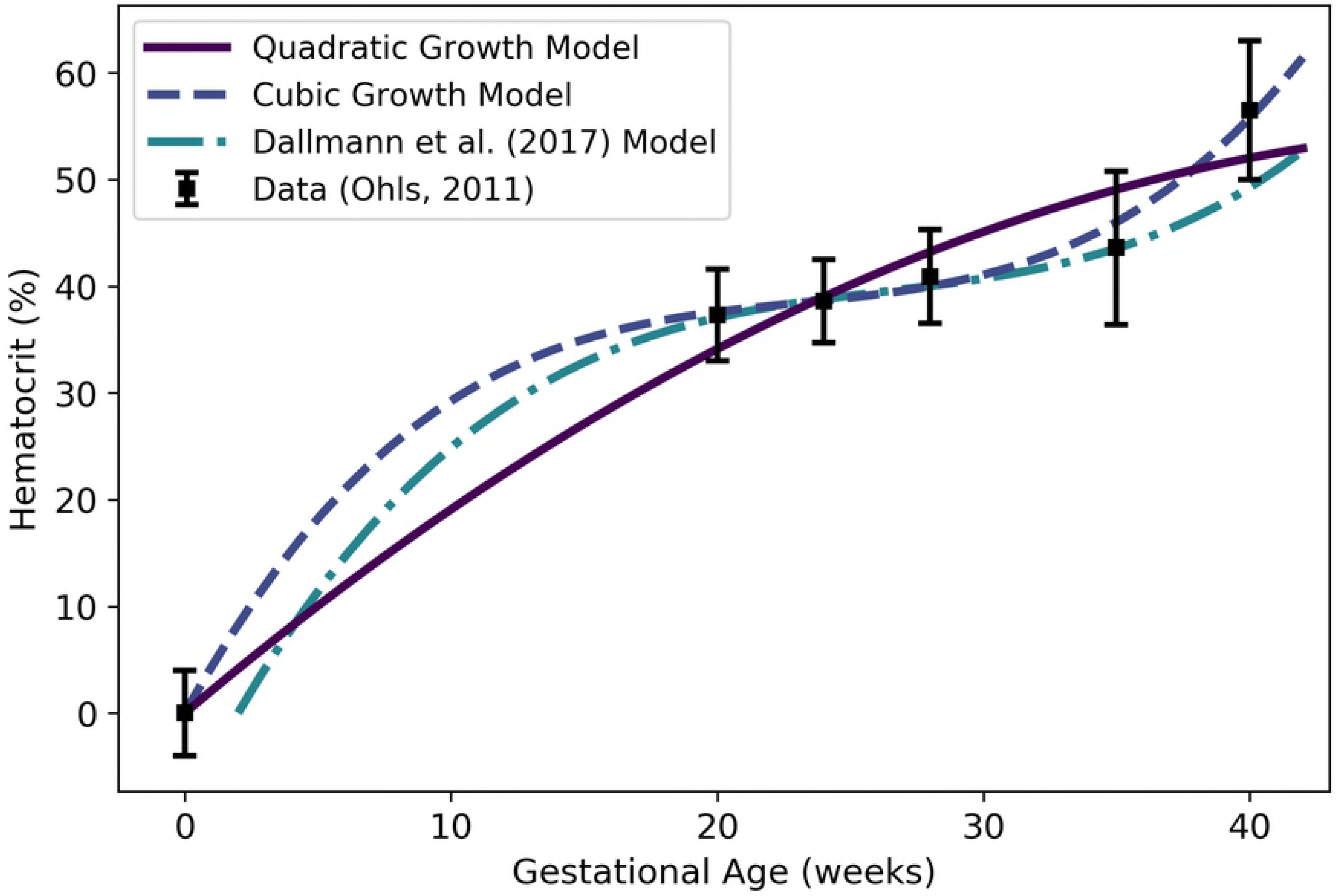
Fetal hematocrit vs. gestational age. The quadratic growth model (solid line) given by Equation 50a was selected as the most parsimonious model in our analysis, but the cubic growth model given by Equation 50b gives a better visual fit to the data while yielding only a slightly (0.8%) larger AIC score. We calibrated our models using the data curated by Ohls [39]. Dallmann et al. [3] calibrated their model with different data.

## DISCUSSION

Here we have presented a comprehensive collection of empirical models for anatomical and physiological changes related to human pregnancy and gestation. (See Table 1 for our list of preferred models.) Several similar repositories have been published [3,15,28,31], but in evaluating each of these we found shortcomings that we sought to address in our own set of empirical models. One distinguishing feature of the current collection of models derives from our recognition that the fetal circulatory system differs fundamentally from the adult circulatory system. Thus, in addition to describing models for those pregnancy- and gestation-related changes that have been included in other collections, we have provided models for blood flow rates in the fetus that do not have analogs in the adult; in particular, we have included models for blood flow rates through the ductus venosus, the ductus arteriosus, and the foramen ovale of the human fetus. To establish a single best model for each quantity of interest, we applied a rigorous set of statistical and mathematical procedures for calibrating a variety of models and selecting an optimal model. Finally, to better inform potential users of the models described herein, we have compared our models to previously-published alternative models wherever possible.

Some of the published models for fetal and maternal changes in anatomical and physiological quantities [15,28,29,36–38] are based on gestational age, or time since the last menstruation, while others [3,31] are based on fetal age, or time since fertilization of the ovum. We have used gestational age as the basis for all the models reported herein since it seems to represent the more standard time scale in descriptions of pregnancy. It is generally acknowledged that a fetal age of zero weeks, or fertilization, coincides with a gestational age of 2 weeks [3,31,40]; thus, none of our models describing fetal quantities or products of conception (e.g., the placenta or amniotic fluid) should be used for predictions prior to gestational age 2 weeks! Whenever necessary (i.e., for data analysis or model comparison), we assumed that gestational age equals fetal age plus two weeks to convert between the two time scales.

One of our primary concerns with previously published repositories of human pregnancy and gestation models is that they tend to neglect models for changes in the developing fetus, and none of them included models for those fetal blood flow rates that do not have analogs in the adult. In fact, with the exception of the collection of models described by Luecke et al. [15], none of these repositories for “pregnancy PBPK” model parameters included any models for fetal blood flow rates (other than flow through the umbilical vein). Furthermore, while Luecke et al. [15] provided models for changes in maternal and fetal blood flow rates that were based upon allometric scaling of corresponding tissue masses, they did not describe the methods and data that were used to calibrate these models.

In examining the models for human pregnancy- and gestation-related changes that are available in the literature, we discovered that some of the models give results that are clearly incorrect and/or substantially different from available data. For example, the maternal plasma volume model of Luecke et al. [15] predicts only small increases (^~^9%) in plasma volume during pregnancy. Our model and the curated data set we used for calibrating it predict considerably larger increases (^~^50%) in plasma volume (cf. Figure 4). Also, the amniotic fluid volume model of Luecke et al. [15] predicts volumes approximately an order of magnitude lower that those seen in the curated data set we used for calibrating our models. We hypothesize that one of the model parameters provided by those authors may have been in error by a factor of 10 (cf. Figure 7 and its caption). In the model collection published by Dallmann et al. [3], we found that the maternal glomerular filtration model (see their Equation 27) contains typographical and scaling errors. Through personal correspondence with the lead author of this work, we confirmed this finding and obtained a corrected version of the formula (cf. Figure 14 and its caption). Finally, the Luecke et al. (1994) models for fetal brain mass, liver mass, and thyroid mass tend to greatly exceed compiled data values for these quantities [31] during later gestational weeks (cf. Figure 15, Figure 16, and Figure 19).

Besides various situations in which previously published models seem to contain errors, we discovered a few instances in which potential users of those models might need to exercise caution. For example, although the careful reader will note that the placenta volume model of Luecke et al. (1994) is only intended for use from days 25 to 300 of gestation, extrapolating to earlier times leads to unreasonably large values for placenta volume (cf. Figure 6). Thus, researchers desiring to estimate placenta volume prior to the 3rd week of gestation should avoid using that model. In an apparent discrepancy with the curated data set [28] we used for calibrating our models for maternal hematocrit, the maternal hematocrit model of Dallmann et al. [3] predicts an increase in hematocrit in late pregnancy. Dallmann et al. [3] used a different curated data set to calibrate their model, and this data does seem to suggest an increase in maternal hematocrit starting at about 31 weeks. Both our preferred model (Equation 21a) and the Dallmann et al. [3] model do, however, agree in their predictions that maternal hematocrit decreases during the first 30 weeks of pregnancy (cf. Figure 13).

None of the previously-published collections of human pregnancy and gestation models explicitly considered mass balance, flow balance, or “rest of body” compartments. In the present work, we have applied the principle of mass balance to provide models for volumes of and flow rates to rest of body compartments for both the mother and the fetus. With the exception of the fetal rest of body volume model, all of these models yield positive values on the entire time domain of interest (i.e., for gestational ages between 0 and 42 weeks), and the fetal rest of body volume model predicts positive volumes for gestational ages greater than 8 weeks. However, as discussed in the Results section, we recommend that the fetal flow rate and volume models not be used for gestational ages prior to 13 weeks due to the general sparsity of fetal data for the first trimester. We have also endeavored evaluate consistency of the flow rate models. For example, based on the basic anatomy of the fetal circulatory system, the flow rate through the ductus venosus should always be less than the flow rate through the placenta. As shown in Figure 27, our models (Equations 42 and 43) satisfy this requirement for gestational ages greater than 13 weeks.

While our collection of models offers many advantages and improvements over similar collections that have been published previously, it does still have some shortcomings. For one, we have provided no description of active transport across the placenta and how this might change throughout pregnancy. Understanding movement of a chemical across the placenta that is not based on passive diffusion may be critically important in assessing the levels of that chemical in both mother and fetus during pregnancy and gestation. Another limitation of our treatment is that most data sets we have used (and therefore most of the models we have presented) describe “healthy” women from predominantly Caucasian populations that were experiencing singleton, low-risk, uncomplicated pregnancies [28,31]. Such women (and their fetuses) may not be representative of the most vulnerable pregnant (and gestating) humans. Furthermore, we have offered no account of population variability in the time-dependent anatomical and physiological quantities we have analyzed. Dallmann et al. [3] attempted to do this by providing empirical models of both a “mean” and “standard deviation” for each quantity of interest they considered; however, using such models of component variability to account for whole-system variability (e.g., in a PBPK model for a mother and her fetus) may lead to unrealistic or unreasonable results, as the individual models for various components (e.g., maternal mass and maternal cardiac output) do not take into account the interdependencies inherent in the corresponding anatomical and physiological parameters [59,60]. One subtle weakness of our analysis is that many of our fetal models tacitly assume the existence of functional organ systems in the embryo and fetus at early gestational weeks. In constructing a PBPK model of human pregnancy and gestation, such assumptions should be carefully evaluated; in fact, in any such modeling effort, perhaps the best strategy would be to model the embryo/fetus as a single “compartment” up until the end of the first trimester, at which time distinct organs and tissues certainly exist in the fetus [40] and can be more reliably characterized. Finally, our repository of models is only valid for *human* mothers and their developing fetuses. For PBPK modeling applications involving intraspecies extrapolation, it would be helpful to have a repository of models comparable to the one contained herein but specific to various non-human animals.

## CONCLUSIONS

All PBPK models involve parameters that describe the anatomy and physiology of an organism, but defining values for these parameters requires special care when designing a PBPK model to represent a mother and fetus during the period from conception to birth. Because pregnancy and gestation represent a time of dramatic changes, many of the PBPK model parameters that are traditionally treated as constants for adult organisms or shorter-duration simulations must be described by time-varying functions in any PBPK model intended for studying this period, either in its entirety or for shorter time periods during which pregnancy- and gestation-related changes might be substantial. To facilitate the development of human pregnancy and gestation PBPK models, we have provided a repository of empirical models describing tissue volumes, blood flow rates, hematocrits, and glomerular filtration rates in an “average” mother and fetus. This collection of models offers advantages over similar repositories in that it includes descriptions for many fetal parameters that have previously been neglected and it uses rigorous and well-defined methods for model calibration and selection. We anticipate that the formulae provided in this collection will provide an improved basis for PBPK models of human pregnancy and gestation, and ultimately support decision-making with respect to optimal pharmacological dosing and risk assessment for pregnant women and their developing fetuses. In fact, we are currently preparing a manuscript that describes a high-throughput toxicokinetic (HTTK) model for pregnancy and gestation that is based upon the formulae provided herein.

## ACKNOWLEDGEMENTS

The authors are grateful to Elaina Kenyon and Paul Schlosser for their careful review of any early draft of this manuscript.

## SUPPORTING INFORMATION CAPTIONS

S1 Appendix. Model Calibration Details.

S2 Python Script. “gest_models.py”. This program contains function definitions for various general types of models described in the manuscript.

S3 Python Script. “maternal_mass.py”. This program uses maternal mass vs. gestational age data (which are contained in the script) to perform model calibrations. It produces a plot of the data and various models for maternal mass vs. gestational age. The plot is saved in a subdirectory called “Output”.

S4 Python Script. “fetal_mass.py”. This program uses fetal mass vs. gestational age data (which are contained in the script) to perform model calibrations. It produces a plot of the data and various models for fetal mass vs. gestational age. The plot is saved in a subdirectory called “Output”.

S5 Python Script. “maternal_fat_mass.py”. This program uses maternal fat mass vs. gestational age data (which are contained in the script) to perform model calibrations. It produces a plot of the data and various models for maternal fat mass vs. gestational age. The plot is saved in a subdirectory called “Output”.

S6 Python Script. “maternal_plasma_volume.py”. This program uses maternal plasma volume vs. gestational age data (which are contained in the script) to perform model calibrations. It produces a plot of the data and various models for maternal plasma volume vs. gestational age. The plot is saved in a subdirectory called “Output”.

S7 Python Script. “maternal_plasma_vol.py”. This program uses maternal plasma volume vs. gestational age data (which are contained in the script) to perform model calibrations. It produces a plot of the data and various models for maternal plasma volume vs. gestational age. The plot is saved in a subdirectory called “Output”.

S8 Python Script. “maternal_rbc_vol.py”. This program uses maternal red blood cell volume vs. gestational age data (which are contained in the script) to perform model calibrations. It produces a plot of the data and various models for maternal red blood cell volume vs. gestational age. The plot is saved in a subdirectory called “Output”.

S9 Python Script. “maternal_placenta_volume.py”. This program uses placenta volume vs. gestational age data (which are contained in the script) to perform model calibrations. It produces a plot of the data and various models for placenta volume vs. gestational age. The plot is saved in a subdirectory called “Output”.

S10 Python Script. “maternal_amnf_vol.py”. This program uses amniotic fluid volume vs. gestational age data (which are contained in the script) to perform model calibrations. It produces a plot of the data and various models for amniotic fluid volume vs. gestational age. The plot is saved in a subdirectory called “Output”.

S11 Python Script. “maternal_mass_balance.py”. This program uses models derived in various other scripts to calculate a model for maternal “rest of body” volume. It produces a plot of the maternal rest of body volume vs. gestational age. The plot is saved in a subdirectory called “Output”.

S12 Python Script. “maternal_cardiac_output.py”. This program uses maternal cardiac output vs. gestational age data (which are contained in the script) to perform model calibrations. It produces a plot of the data and various models for maternal cardiac output vs. gestational age. The plot is saved in a subdirectory called “Output”.

S13 Python Script. “maternal_kidney_flow.py”. This program uses maternal kidney blood flow vs. gestational age data (which are contained in the script) to perform model calibrations. It produces a plot of the data and various models for maternal kidney blood flow vs. gestational age. The plot is saved in a subdirectory called “Output”.

S14 Python Script. “maternal_placenta_flow.py”. This program blood flow proportion estimates and other models to generate a model for maternal blood flow to the placenta vs. gestational age. It produces a plot of various models for maternal blood flow to the placenta vs. gestational age. The plot is saved in a subdirectory called “Output”.

S15 Python Script. “maternal_flow_balance.py”. This program uses models derived in various other scripts to calculate a model for blood flow to the maternal “rest of body”. It produces a plot of maternal blood flow to the rest of body vs. gestational age. The plot is saved in a subdirectory called “Output”.

S16 Python Script. “maternal_hematocrit.py”. This program uses maternal hematocrit vs. gestational age data (which are contained in the script) to perform model calibrations. It produces a plot of the data and various models for maternal hematocrit vs. gestational age. The plot is saved in a subdirectory called “Output”.

S17 Python Script. “maternal_gfr.py”. This program uses maternal glomerular filtration rate vs. gestational age data (which are contained in the script) to perform model calibrations. It produces a plot of the data and various models for maternal glomerular filtration rate vs. gestational age. The plot is saved in a subdirectory called “Output”.

S18 Python Script. “fetal_brain_mass.py”. This program uses fetal brain mass vs. gestational age data (which are contained in the script) to perform model calibrations. It produces a plot of the data and various models for fetal brain mass vs. gestational age. The plot is saved in a subdirectory called “Output”.

S19 Python Script. “fetal_liver_mass.py”. This program uses fetal liver mass vs. gestational age data (which are contained in the script) to perform model calibrations. It produces a plot of the data and various models for fetal liver mass vs. gestational age. The plot is saved in a subdirectory called “Output”.

S20 Python Script. “fetal_kidney_mass.py”. This program uses fetal kidney mass vs. gestational age data (which are contained in the script) to perform model calibrations. It produces a plot of the data and various models for fetal kidney mass vs. gestational age. The plot is saved in a subdirectory called “Output”.

S21 Python Script. “fetal_lung_mass.py”. This program uses fetal lung mass vs. gestational age data (which are contained in the script) to perform model calibrations. It produces a plot of the data and various models for fetal lung mass vs. gestational age. The plot is saved in a subdirectory called “Output”.

S22 Python Script. “fetal_thyroid_mass.py”. This program uses fetal thyroid mass vs. gestational age data (which are contained in the script) to perform model calibrations. It produces a plot of the data and various models for fetal thyroid mass vs. gestational age. The plot is saved in a subdirectory called “Output”.

S23 Python Script. “fetal_gut_mass.py”. This program uses fetal gut mass vs. gestational age data (which are contained in the script) to perform model calibrations. It produces a plot of the data and various models for fetal gut mass vs. gestational age. The plot is saved in a subdirectory called “Output”.

S24 Python Script. “fetal_mass_balance.py”. This program uses models derived in various other scripts to calculate a model for fetal “rest of body” volume. It produces a plot of the fetal rest of body volume vs. gestational age. The plot is saved in a subdirectory called “Output”.

S25 Python Script. “fetal_rvtl_flow.py”. This program uses fetal right ventricle blood flow vs. gestational age data (which are contained in a file called “Kiserud2006_Fig5A_Q_rvtl.csv” in a subdirectory called “Data”) to perform model calibrations. It produces a plot of the data and various models for fetal right ventricle blood flow data vs. gestational age. The plot is saved in a subdirectory called “Output”.

S26 Python Script. “fetal_rvtl_flow.py”. This program uses fetal left ventricle blood flow vs. gestational age data (which are contained in a file called “Kiserud2006_Fig4A_Q_lvtl.csv” in a subdirectory called “Data”) to perform model calibrations. It produces a plot of the data and various models for fetal left ventricle blood flow data vs. gestational age. The plot is saved in a subdirectory called “Output”.

S27 Python Script. “fetal_DA_flow.py”. This program uses fetal ductus arteriosus blood flow vs. gestational age data (which are contained in a file called “Mielke2001_Fig8_Q_DA.csv” in a subdirectory called “Data”) to perform model calibrations. It produces a plot of the data and various models for fetal ductus arteriosus blood flow data vs. gestational age. The plot is saved in a subdirectory called “Output”.

S28 Python Script. “fetal_plac_flow.py”. This program uses fetal umbilical vein blood flow vs. gestational age data (which are contained in a file called “Kiserud2000_Fig5A_Q_plac.csv” in a subdirectory called “Data”) to perform model calibrations. It produces a plot of the data and various models for fetal placenta blood flow data vs. gestational age. The plot is saved in a subdirectory called “Output”.

S29 Python Script. “fetal_DV_flow.py”. This program uses fetal ductus venosus blood flow vs. gestational age data (which are contained in a file called “Kiserud2000_Fig5B_Q_DV.csv” in a subdirectory called “Data”) to perform model calibrations. It produces a plot of the data and various models for fetal ductus venosus blood flow data vs. gestational age. The plot is saved in a subdirectory called “Output”.

S30 Python Script. “fetal_flow_balance.py”. This program uses models derived in various other scripts to calculate a model for blood flow to the fetal “rest of body”. It produces a plot of fetal blood flow to the rest of body vs. gestational age. The plot is saved in a subdirectory called “Output”.

S31 Python Script. “fetal_hematocrit.py”. This program uses fetal hematocrit vs. gestational age data (which are contained in the script) to perform model calibrations. It produces a plot of the data and various models for fetal hematocrit vs. gestational age. The plot is saved in a subdirectory called “Output”.

S32 Folder. “Output”. This folder must be in the directory or folder that contains the Python scripts. It is used for saving various types of output, including images (in PNG and TIF formats), and optimal model parameters (in “.npy” files).

S33 Folder. “Input”. This folder must be in the directory or folder that contains the Python scripts. It is used to contain raw data files.

S34 Data File. “Kiserud2006_Fig5A_Q_rvtl.csv”. This file contains data that were digitally extracted from Figure 5A of Kiserud et al. [36]. The file is located in a subdirectory called “Input”.

S35 Data File. “Kiserud2006_Fig4A_Q_lvtl.csv”. This file contains data that were digitally extracted from Figure 4A of Kiserud et al. [36]. The file is located in a subdirectory called “Input”.

S36 Data File. “Mielke2001_Fig8_Q_DA.csv”. This file contains data that were digitally extracted from Figure 4A of Mielke and Benda [37]. The file is located in a subdirectory called “Input”.

S37 Data File. “Kiserud2000_Fig5A_Q_plac.csv”. This file contains data that were digitally extracted from Figure 5A of Kiserud et al. [38]. The file is located in a subdirectory called “Input”.

S38 Data File. “Kiserud2000_Fig5B_Q_DV.csv”. This file contains data that were digitally extracted from Figure 5B of Kiserud et al. [38]. The file is located in a subdirectory called “Input”.

